# Group 2 innate lymphoid cells constrain type 3/17 lymphocytes in shared stromal niches to restrict liver fibrosis

**DOI:** 10.1101/2023.04.26.537913

**Authors:** Julia Sbierski-Kind, Kelly M Cautivo, Johanna C Wagner, Madelene W Dahlgren, Julia Nilsson, Maria Krasilnikov, Nicholas M. Mroz, Carlos O. Lizama, Anna Lu Gan, Peri R Matatia, Marcela T Taruselli, Anthony A Chang, Sofia Caryotakis, Claire E O’Leary, Maya Kotas, Aras N Mattis, Tien Peng, Richard M Locksley, Ari B Molofsky

**Affiliations:** Department of Laboratory Medicine, University of California San Francisco; Department of Surgery, Division of Transplantation, University of California San Francisco; Department of Medicine, Pulmonary Division, University of California San Francisco; Biomedical Sciences Graduate Program, University of California San Francisco; Cardiovascular Research Institute, University of California San Francisco; Department of Medicine, University of California San Francisco; Department of Pediatrics, School of Medicine and Public Health, University of Wisconsin – Madison, Madison Wisconsin; Liver Center, University of California San Francisco; Department of Pathology, University of California San Francisco; Department of Microbiology and Immunology, University of California San Francisco; Howard Hughes Medical Institute, University of California San Francisco; Diabetes Center, University of California San Francisco

**Keywords:** Type 2 immunity, allergic immunity, ILC2, type 3 immunity, type 17 immunity, γδ T cell, tissue immunology, fibrosis, stromal cells, myofibroblast, adventitial fibroblast, volumetric imaging, quantitative imaging, IL-5, IL-13, IL-17A, IL-33, tissue niches

## Abstract

Group 2 innate lymphoid cells (ILC2s) cooperate with adaptive Th2 cells as key organizers of tissue type 2 immune responses, while a spectrum of innate and adaptive lymphocytes coordinate early type 3/17 immunity. Both type 2 and type 3/17 lymphocyte associated cytokines are linked to tissue fibrosis, but how their dynamic and spatial topographies may direct beneficial or pathologic organ remodelling is unclear. Here we used volumetric imaging in models of liver fibrosis, finding accumulation of periportal and fibrotic tract IL-5^+^ lymphocytes, predominantly ILC2s, in close proximity to expanded type 3/17 lymphocytes and IL-33^high^ niche fibroblasts. Ablation of IL-5^+^ lymphocytes worsened carbon tetrachloride-and bile duct ligation-induced liver fibrosis with increased niche IL-17A^+^ type 3/17 lymphocytes, predominantly γδ T cells. In contrast, concurrent ablation of IL-5^+^ and IL-17A^+^ lymphocytes reduced this progressive liver fibrosis, suggesting a cross-regulation of type 2 and type 3 lymphocytes at specialized fibroblast niches that tunes hepatic fibrosis.

## HIGHLIGHTS

- IL-5^+^ and IL-17A^+^, but not IFNγ^+^, lymphocytes are enriched in periportal liver niches.
- Liver fibrosis drives IL-5^+^ and IL-17A^+^ lymphocyte accumulation at de novo fibrotic regions.
- Loss of IL-5^+^ type 2 lymphocytes heightened fibrosis-associated IL-17A^+^ type 3 lymphocytes and exacerbated liver fibrosis.
- Concurrent loss of IL-5^+^ and IL-17A^+^ lymphocytes attenuates liver inflammation and fibrosis.

## INTRODUCTION

Fibroproliferative diseases affect virtually all organ systems and are major drivers of worldwide morbidity and mortality, with particularly severe consequences in the liver (Gieseck et al., 2018a). Chronic liver (hepatic) injury and inflammation, driven by viral infections (*e.g.,* hepatitis B or C), obstruction or impairment of bile drainage (*e.g.,* gallstones, biliary/pancreatic cancer), or (non-)alcohol associated steatohepatitis (NASH/ASH), each increase the risk of chronic liver cirrhosis and downstream hepatocellular carcinoma (Hernandez-Gea and Friedman, 2011). <u>Fibrogenic</u> and <u>inflammatory</u> pathways synergistically regulate liver fibrosis through activation of resident stromal cells (*e.g.,* hepatic stellate cells and periportal fibroblasts), promoting the development of pathogenic, matrix-depositing myofibroblasts.

Tissue-resident lymphocytes (TRLs) are critical ‘upstream’ regulators of inflammatory pathways, secreting cytokines to coordinate discrete anti-microbial responses as well as aspects of organ development, remodeling, and repair (Fan and Rudensky, 2016; Annunziato et al., 2015). As the major TRLs, T cells and innate lymphoid cells (ILCs) are broadly categorized into three groups with associated canonical cytokines: type 1 lymphocytes and IFNγ; type 2 lymphocytes and IL-4, IL-5, IL-9, and IL-13; and type 3/17 lymphocytes and IL-17A/F and IL-22. These lymphocyte groups share regulation and effector function among CD4^+^ helper T cells (Th1, Th2, Th17), innate lymphoid cells (ILC1, ILC2, and ILC3), and unconventional/innate-like T cells (γδ, NK/T, MAIT T cells, etc). Although plasticity exists, and lymphocytes do not always fit precisely in such bins, these categories remain useful constructs to understand broad patterns of tissue immunity (Tortola et al., 2020).

Type 3/17 immunity restricts extracellular bacteria and fungi at barrier surfaces and is associated with beneficial tissue regeneration; in excess, it can drive fibrosis and inflammatory disease, often mediated by IL-17A and IL-22 acting on diverse non-hematopoietic tissue cell types (Molina et al., 2019; Fabre et al., 2014; Meng et al., 2012; Tedesco et al., 2018; Wree et al., 2018; Choy et al., 2015). Type 2 ‘allergic’ inflammation is canonically activated by multicellular helminths, protozoa, and other noxious stimuli. It is coordinated by tissue <u>type 2 lymphocytes</u> (ILC2s and Th2s) that secrete IL-4, IL-5, IL-9, and IL-13 (Fort et al., 2001; Hurst et al., 2002; Moro et al., 2010; Neill et al., 2010; Price et al., 2010; Ruterbusch et al., 2020; Zeis et al., 2020; Gasteiger et al., 2015; Molofsky et al., 2015). IL-4 and IL-13 signal through the shared IL-4Ra receptor and can also drive maladaptive type 2 inflammation (Gieseck et al., 2016; Wynn and Vannella, 2016; Saluzzo et al., 2017). Type 2 immunity is regulated by diverse upstream damage signals, including IL-33 which drives type 2 lymphocytes to proliferate and produce IL-13; in excess, both IL-33 and IL-13 are sufficient to drive liver fibrosis (Mchedlidze et al., 2013; Tan et al., 2018; Gao et al., 2016). However, type 2 immunity can also promote beneficial tissue development, remodeling, and repair (Allen and Sutherland, 2014; Lloyd and Snelgrove, 2018). ILC2s are implicated in both chronic inflammatory diseases and helminth-driven fibrosis (Mchedlidze et al., 2013; Hams et al., 2014; Forkel et al., 2017), but it is unknown whether, or how, they may impact the development of liver fibrosis in settings not intrinsically type 2 immune-skewed.

Type 2 lymphocytes are enriched in adventitial regions of larger vessels and other ‘border’ structures in multiple tissues, including periportal liver regions (Dahlgren et al., 2019; Cautivo et al., 2022). In naïve mice, type 2 lymphocytes (*e.g*., ILC2s and less frequent Th2s) have a shared transcriptomic and epigenomic signature, including expression of IL-5 and IL-13 (Lee et al., 2015; Nussbaum et al., 2013). Adventitial border regions are defined by a fibroblast subset, or state, that produces thymic stromal lymphopoietin (TSLP) and high levels of IL-33, designated adventitial fibroblasts (AFs), and supports IL-5^+^ type 2 lymphocytes (Dahlgren et al., 2019). AFs are found across organs as a universal state and have increased progenitor capacity, including the ability to differentiate into pathogenic myofibroblasts that drive organ fibrosis (Sbierski-Kind et al., 2021). However, the potential function and localization of AF-like IL-33^high^ ‘niche’ fibroblasts, type 2, and type 3/17 lymphocytes during liver fibrosis is unknown.

Here we found that hepatic fibrosis driven by carbon tetrachloride (CCl_4_)-or bile duct ligation (BDL) promotes expansion of IL-33^high^ AF-like fibroblasts in proximity to expanded IL-5^+^ lymphocytes (predominantly ILC2s) and type 3/17 lymphocytes (IL-17A^+^ Rorγt^+^, predominantly γδ T cells) in periportal adventitia regions, as well as de novo fibrotic tracts. Using selective ablation of IL-5^+^ lymphocytes, we found that loss of type 2 lymphocytes exacerbated liver fibrosis in both fibrotic models, associated with increasing type 3/17 inflammation. Co-depletion of IL-5^+^ and IL-17A^+^ lymphocytes attenuated the enhanced CCl_4_-induced hepatic fibrosis and liver inflammation. Our data suggest crosstalk between type 2 and type 3/17 lymphocytes in specialized fibroblast niches that tune the degree of liver damage and fibrosis.

## RESULTS

### Models of liver fibrosis

The liver has a unique tissue macrostructure, with *portal triads* comprised of portal veins carrying *first-pass* venous blood from the intestine, hepatic arteries with oxygenated blood, and ducts draining bile into the small intestine. Blood percolates from portal regions through the liver parenchyma, defined by hepatocytes, porous sinusoidal endothelium, and hepatic stellate cells (HSCs, *e.g.,* liver mural cells/pericytes), and is ultimately drained by central veins (**Fig. 1A**). We used three established mouse models to assess how diverse stromal cells and lymphocytes mapped onto this unique topography. (1) Repetitive doses of <u>carbon tetrachloride</u> (CCl_4_): liver toxin that causes hepatocyte injury and centrilobular liver fibrosis **(Fig. S1A)** (Tsuchida et al., 2018). These mice developed pericentral fibrosis, with liver damage (serum alanine transaminase), collagen deposition and fibrosis (Sirius Red staining, hydroxyproline levels), increased liver immune cells, and loss of pericentral hepatocytes (decreased glutamine synthetase) (Ghallab et al., 2019) (**Fig. S1B-F**). (2) <u>Bile duct ligation</u> (BDL): surgical model leading to acute obstructive jaundice and rapid progression to cholestasis-induced liver fibrosis focused in periportal regions **(Fig. 1A, S1G)** (Saito and Maher, 2000). At 14 days post BDL, mice had increased liver damage, cholestasis, liver collagen deposition at peri-portal bile duct-containing regions, and total liver fibrosis (**Fig. SH-K)**. To better mimic the metabolic derangements associated with NASH, we also used a (3) <u>fructose, palmitate and cholesterol-rich diet</u> <u>(FPC diet, 16 weeks)</u> (Wang et al., 2016) **(Fig. 1A, S1L)**. FPC-fed mice had increased weight with liver damage, patchy fibrosis, and fat droplet deposition; liver and visceral white adipose tissue (WAT) weights were both increased (**Fig. S1M-S)**. These initial data validated our models to reproduce distinct clinicopathologic and topographic features of liver disease and associated fibrosis.

**Figure 1:**
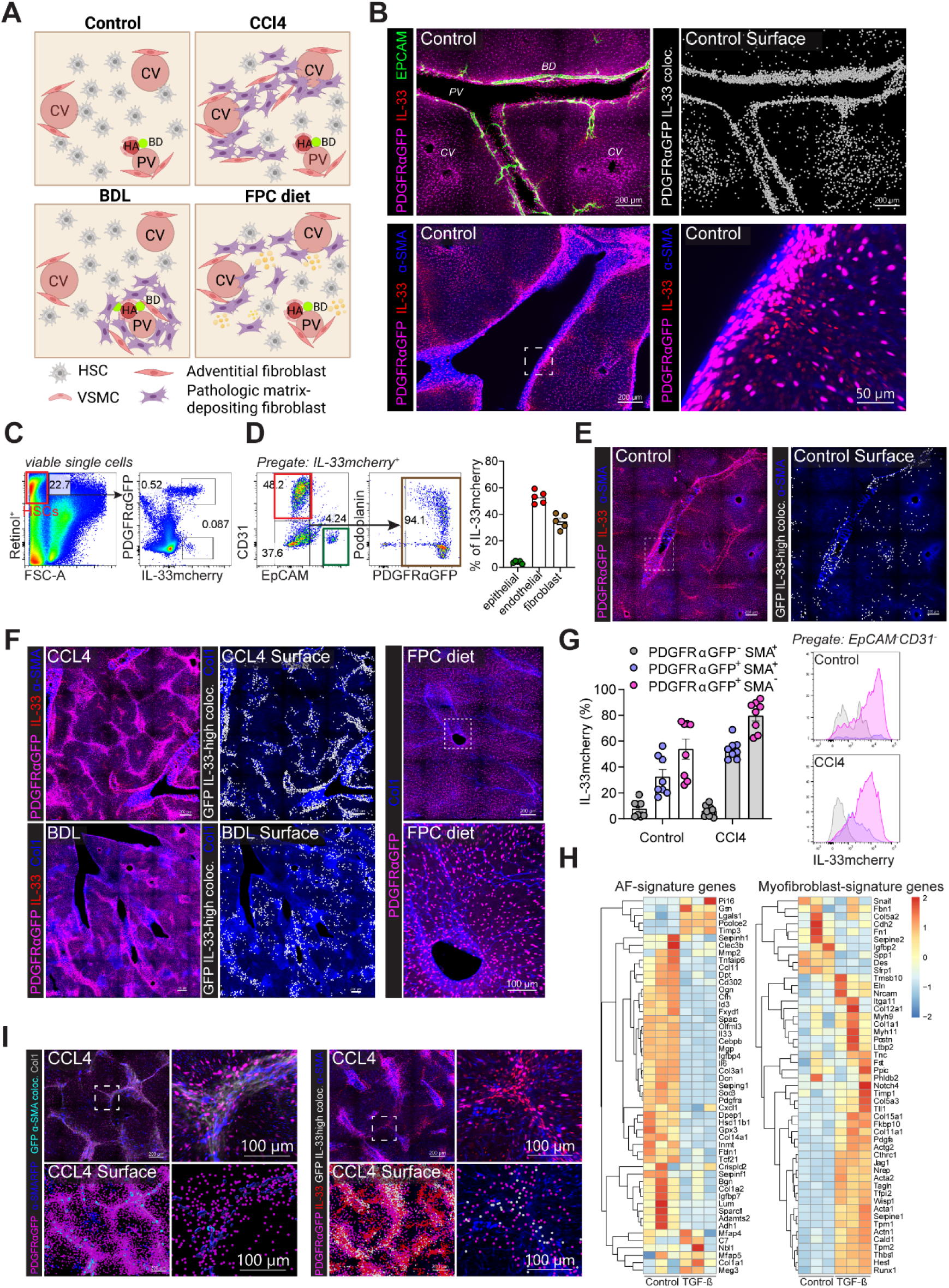
PDGFRαGFP^+^ fibroblasts express IL-33mcherry and localize to fibrotic tracts in hepatic fibrosis. (A) Schematic of the topography of heterogenous mesenchymal subpopulations in healthy liver lobule and in different models of advanced hepatic fibrosis. Carbon tetrachloride (CCl_4_), bile duct ligation (BDL), central vein (CV), portal vein (PV). (B) Confocal thick section images (200µm) from naïve livers of PDGFRαGFP; IL-33^mcherry/+^ mice with antibody stains indicated and PDGFRαGFP-IL-33^mcherry^ colocalization highlighted (white). (C) Representative flow cytometry plots from livers of 4w CCl_4_-treated PDGFRαGFP; IL-33^mcherry/+^ reporter mice, showing expression of PDGFRαGFP and IL-33mcherry in hepatic stellate cells (HSCs). (D) Representative flow cytometry plots and quantification from livers of 4w CCl_4_-treated PDGFRαGFP; IL-33^mcherry/+^ reporter mice, showing percent epithelial (EpCAM^+^), endothelial cells (CD31^+^), and fibroblasts (CD31^-^EpCAM^-^PDGFRαGFP^+^) of IL33mcherry^+^ non-hematopoietic CD45^-^ cells. (E and F) Representative thick section confocal images of livers from PDGFRαGFP; IL-33mcherry reporter mice at steady state (E) and after the indicated challenges (F). Antibody stains are indicated, and PDGFRαGFP-IL-33mcherry co-localization is highlighted in white. Images are representative of 1-4 independent experiments with n=6-13 mice/group. (G) Quantification and representative histograms of percentages IL-33mcherry expression of different fibroblast subsets defined by PDGFRαGFP and SMA. Histograms and bar graphs are representative of 4 repeat experiments with n=5-10 mice/group. (H) Heatmaps of the top adventitial stromal cell (ASC)-and myofibroblast-expressed genes in in vitro cultured lung stromal cells (CD45^-^CD31^-^EpCAM^-^PDGFRα^+^Sca-1^+^) stimulated with TGF-β for 48 hrs. (I) Representative thick section confocal images of livers from 4w CCl_4_-treated and PDGFRαGFP; IL-33mcherry reporter mice (right). Antibody stains are indicated, and PDGFRαGFP-IL-33mcherry co-localization is highlighted in white. Images are representative of 1-4 independent experiments with n=6-13 mice/group. Bar graphs indicate mean (±SE). See also Figure S1.

### Mapping IL-33^high^ fibroblasts and SMA^+^ (myo)fibroblasts in liver fibrosis

Next, we examined the spatial distribution of liver fibroblast-like stromal cells, including IL-33-high AFs previously shown to localize to tissue surface borders and vascular boundaries that commonly define lymphocyte niches (Dahlgren et al., 2019; Cautivo el al., 2022) **(Fig. 1A)**. Using thick-section confocal microscopy with quantitation, naïve liver stromal cells in periportal areas (e.g., periportal fibroblasts) expressed high IL-33 (PDGFRa^GFP+^ IL33^mcherry-high^) (Marvie et al., 2010); however, some parenchymal stromal cells (e.g., HSCs) also expressed low/variable IL-33 **(Fig. 1B; Video S1)**. Flow cytometry confirmed that HSCs expressed low levels of IL-33 compared to liver periportal fibroblasts (**Fig. 1C, D).** Liver venous endothelial cells and rare epithelial cells also expressed IL-33^mcherry^ **(Fig. 1D),** as described previously (Dahlgren et al., 2019).

Having established an approach to track adventitial-like ‘niche’ fibroblasts, we mapped their topography during liver fibrosis and compared with traditional SMA^+^ pathogenic fibroblasts. Central-vein associated HSCs are a major source of pathogenic myofibroblasts in centrilobular fibrosis such as CCl_4_ (Dobie et al., 2019a), whereas periportal fibroblasts are dominant drivers of periportal and cholestatic fibrosis, including in BDL models (Di Carlo and Peduto, 2018). CCl_4_ increased IL-33^high^ fibroblasts in both periportal regions and along de-novo fibrotic tracts associated with central veins **(Fig. 1E, F; Video S2)**. BDL surgery expanded IL-33^high^ periportal fibroblasts, whereas FPC diet promoted mild stromal cell/fibroblast increases across liver domains **(Fig. 1F)**. Flow cytometry confirmed myofibroblasts were also elevated in fibrotic livers (αSMA^+^, variable PDGFRαGFP), but expressed scant IL-33 **(Fig. 1G, Fig. S1T)** (Di Carlo and Peduto, 2018), suggesting myofibroblast and IL-33^high^ AF-like fibroblast states may be discrete.

Next, we hoped to better define the signature of pathogenic myofibroblasts compared to IL-33^high^ fibroblasts. As viable liver periportal fibroblasts were not readily isolated (not shown), we turned to lung AFs, which have conserved features with liver periportal fibroblasts and can differentiate into myofibroblasts (Buechler et al., 2021). We found that TGFβ treatment in primary AFs dramatically repressed the transcriptomic signature associated with ‘AF-like’ niche fibroblasts, including *Pdgfra*, several cytokines (*Il33, Il6*), chemokines (*Cxcl1, Ccl11*), growth factor-binding proteins (*Igfbp4, Igfbp7*), and transcription factors (*Cebpb*) (Dahlgren et al., 2019); instead, myofibroblast-associated genes (*Cthrc1, Col1a1, Acta2, Postn*) were upregulated **(Fig. 1H)** (Tsukui et al., 2020). Further, imaging of CCl_4_-treated mice identified putative myofibroblasts within fibrotic tracts (PDGFRαGFP^+/-^ SMA^+/-^) with low IL-33 expression; in contrast, IL-33^high^ AF-like fibroblasts were in proximity to large SMA^+^ myofibroblasts but were dispersed along the borders of fibrotic tracts **(Fig. 1I).** Together, these data suggest liver stromal heterogeneity during fibrosis **(Fig. S1U),** with IL-33^high^ fibroblasts enriched in periportal and *de novo* fibrotic regions that are in proximity to (but distinct from) ‘pathogenic’ myofibroblasts (Tsukui et al., 2020) that emerge in fibrotic zones after liver challenge.

### Type 2 and type 3/17 lymphocytes expand in liver damage and fibrosis

Next we evaluated type 2 lymphocytes in models of liver fibrosis. Liver ILC2s expanded over the course of CCl_4_ treatment (**Fig. 2A-C,** gating strategy in **Fig. S2A)**, similar to patients with NASH and hepatic fibrosis (Forkel et al., 2017). To further characterize and localize ILC2s, we used well-defined IL-5 cytokine reporter mice, often in combination with a lineage-tracker/ fate-mapper allele (Il5-TdT-Cre; R26-fsf-TdT) (Nussbaum et al., 2013; Molofsky et al., 2013; Dahlgren et al., 2019; Cautivo et al., 2022) **(Fig. S2B)**. ILC2s comprised ∼70-80% of IL-5^+^ liver cells, with the remainder identified as T cells (*i.e.,* Th2s and innate-like CD4^-^/CD8^-^ T cells; **Fig. 2D, E**). IL-5 production (TdT expression) was higher in liver ILC2s compared to T cells both at rest and after CCl_4_ treatment **(Fig. 2F)**. Both ILC2s and rare IL-5^+^ T cells were increased during CCl_4_ fibrosis; however, their expression of IL-5, IL-13, and CD25 (IL2Ra) was not changed (**Fig. S2C-G**), suggesting a type 2 lymphocyte expansion without overt changes in activation or cytokine production.

**Figure 2:**
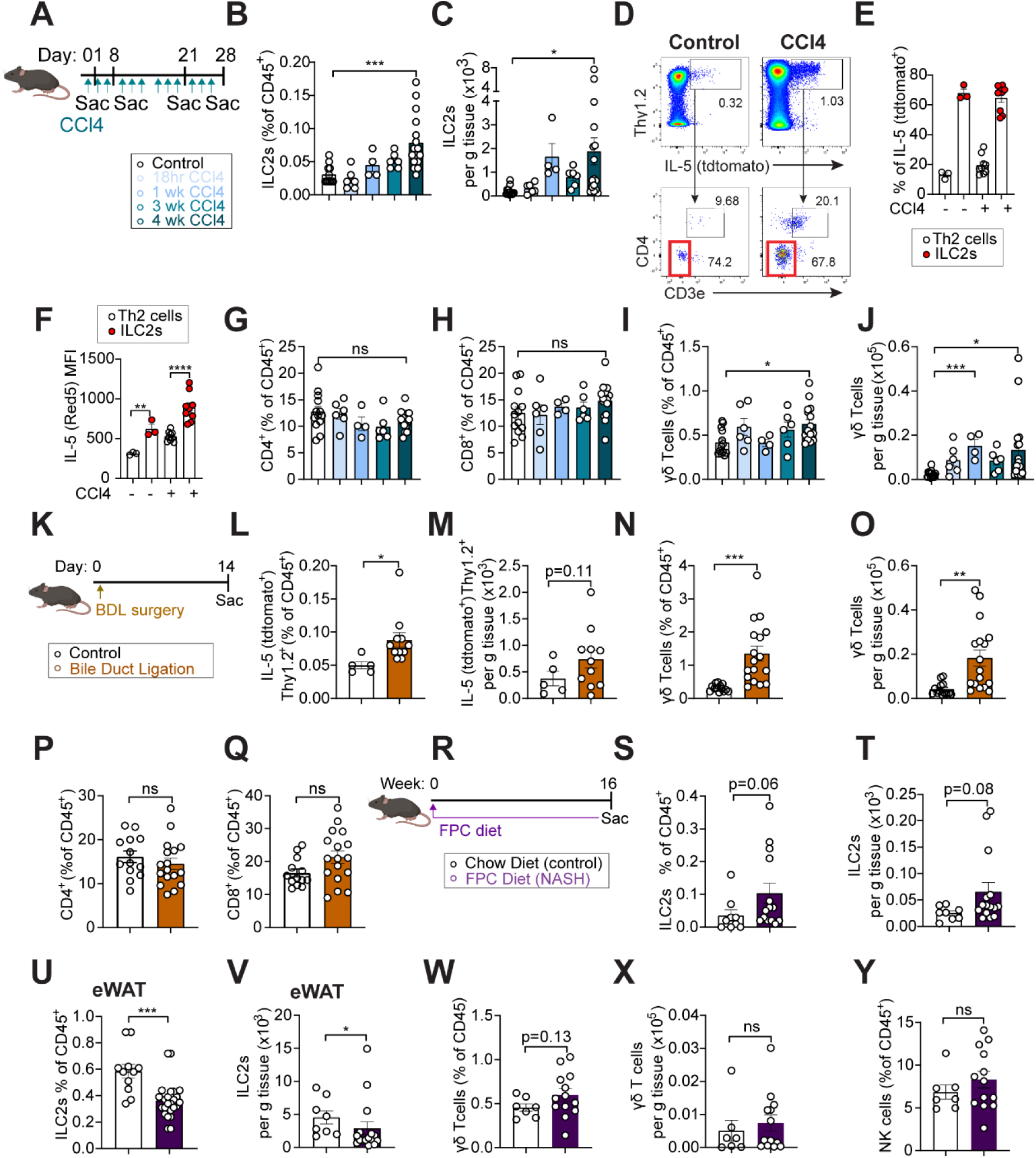
ILC2s and γδ T cells expand in different models of liver fibrosis. (A) Schematic showing CCl_4_ administration schedule in IL-33^mcherry/+^ reporter mice injected intraperitoneally (i.p.) with 1µl CCl_4_/g body weight (BW) once (18h) or with 0.5µl CCl_4_/g BW three times per week for 1, 3, or 4 weeks (1w, 3w, 4w), relevant to B, C and G-J. (B and C) Flow cytometry quantification, showing percent (B) and total numbers of ILC2s (C) in livers from IL-33^mcherry/+^ reporter mice treated with vehicle (corn oil) or CCl_4_ for the indicated duration. Pooled data from 3 independent experiments from 8-10 weeks-old age-and sex-matched mice for 4-week timepoint. (D and E) Flow cytometry plots (D) and quantification (E) of hepatic IL-5^+^ lymphocytes gated on CD45^+^CD11b^-^CD19^-^NK1.1^-^ Thy1.2^+^ and IL-5^+^ from control (treated with corn oil) and CCl_4_-treated Il5-tdtomato-Cre mice, highlighting ILC2s (IL-5^+^CD3^-^CD4^-^) and Th2s (IL-5^+^CD3^+^CD4^+^). (F) Flow quantification of Red5MFI of liver IL-5^+^ lymphocytes (Lin^-^Thy1.2^+^RFP^+^) from Il5-tdtomato-Cre mice mice at rest or treated with CCl_4_ for 4 weeks. (G-J) Flow cytometry quantification, showing percent CD4^+^ T cells (G), percent CD8^+^ T cells (H), and percent (I) and total numbers (J) of liver γδ T cells, from IL-33mchery reporter mice treated with vehicle or CCl_4_ for the indicated duration. Pooled data from 3 independent experiments from 8-10 weeks-old age-and sex-matched mice for 4-week timepoint. (K) Schematic showing schedule for bile duct ligation (BDL) surgery in IL-5^+^ lymphocyte lineage tracker mice (IL-5tdtomato-Cre; Rosa26RFP). (L-Q) Flow cytometry quantification, showing percent ILC2s (L) and γδ T cells (N), total numbers of ILC2s (M) and γδ T cells (O), and percent CD4^+^ (P) and CD8^+^ T cells (Q) in livers from IL-33^mcherry/+^ reporter mice 14 days post-BDL surgery. Pooled data from 3 independent experiments from 8-10-weeks-old age-and sex-matched mice. (R) Schematic showing administration of diet rich in fructose, palmitate, and cholesterol (FPC diet) for 16 weeks in IL-33^mcherry/+^ reporter mice. (S-Y) Flow cytometry quantification, showing percent (S) and total numbers (T) of liver ILC2s, percent (U) and total numbers (V) of ILC2s in epigonadal adipose tissue (eWAT), percent (W) and total numbers (X) of liver γδ T cells, and percent liver NK cells (Y) from IL-33^mcherry/+^ and PDGFRαGFP reporter mice fed with chow diet or cholesterol-enriched FPC diet for 16 weeks. Data pooled from 3 independent experiments from 6-8-weeks-old age-and sex-matched mice. Bar graphs indicate mean (±SE), Two-Way ANOVA with Sidak post-test (B, C, G-J) or unpaired t-test (F, L-Q, S-Y), *p ≤ 0.05, **p ≤ 0.01, ***p ≤ 0.001. See also Figure S2.

To determine if the expansion of liver IL-5^+^ lymphocytes was unique, we next profiled type 1 and type 3/17 cytokine producing lymphocytes. During CCl_4_ fibrosis, total liver CD4^+^ and CD8^+^ T cells expanded but with an unchanged ratio (**Fig. 2G, H**, not shown). However, total γδ T cells, Th17 cells, and relative IL-17A expression within γδ T cells all increased (**Fig. 2I-J, Fig S2H-I**), although IFNγ^+^ CD4^+^ T cells and IL-13^+^ T cells were relatively decreased (**Fig. S2J-K**). Tregs were also mildly elevated (**Fig. S2L-M**). The BDL model was also associated with increased IL-5^+^ lymphocytes, γδ T cells, and neutrophils at 14 days post-surgery, with modest relative increases in Th1, Th2, and Th17 cells (**Fig. 2K-Q**, **Fig. SH, J, K, N, O**). FPC-diet fed mice had modestly expanded liver ILC2s (**Fig. 2R-T**) with concurrent decreased ILC2s in expanded epididymal white adipose tissue (eWAT, **Fig 2U-V**) (Molofsky et al., 2013). FPC diet did not show significant changes in γδ T cells and NK cells, although they were trending upwards **(Fig. 2W-Y)**. Together, these results support a conserved and preferential liver fibrosis-associated increase in IL-5^+^ type 2 lymphocytes and IL-17A^+^ type 3/17 lymphocytes.

### Type 2 lymphocytes localize to periportal regions at steady state and expand into fibrotic tracts with liver fibrosis

Next we used volumetric confocal imaging to determine the positioning of liver type 2 lymphocytes, including their spatial relationship to fibroblasts. In naïve livers, IL-5^+^ lymphocytes were primarily in proximity to fibroblasts and collagen-dense networks, with enrichment in the periportal adventitia near bile ducts and hepatic arteries (Dahlgren et al., 2019; Cautivo et al., 2022) (**Fig. 3A-B, Fig. S3A-B; Video S3-S4)**. In all three fibrosis models, there were increased IL-5^+^ lymphocytes in collagen-dense perivascular (*i.e*., periportal and central vein associated) and *de novo* fibrotic tracts (**Fig. 3A-F, Fig. S3A-E; Video S5).** After BDL, a majority of expanded IL-5^+^ lymphocytes localized in collagen-dense regions near bile ducts, concomitant with ductal proliferation and portal fibrosis (**Fig. 3C, G, Fig. S3D**). IL-5^+^ lymphocytes were also closer to the nearest fibroblast after CCl_4_-induced fibrosis (**Fig. 3H-I)**. We also tested if IL-5^+^ lymphocyte expansion was a conserved feature of fibrosis in the lung, using the bleomycin model (Degryse et al., 2010) **(Fig. S3F-G**). Lung IL-5^+^ lymphocytes expanded modestly and were associated with collagen-dense adventitial and *de novo* fibrotic lung regions, but did not expand in the blood (**Fig. S3H-J**). Together, these data indicate that IL-5^+^ type 2 lymphocytes increase in multiple models of fibrosis, primarily residing in collagen-dense adventitial and *de novo* fibrotic regions in close proximity to a subset of expanded fibroblasts.

**Figure 3:**
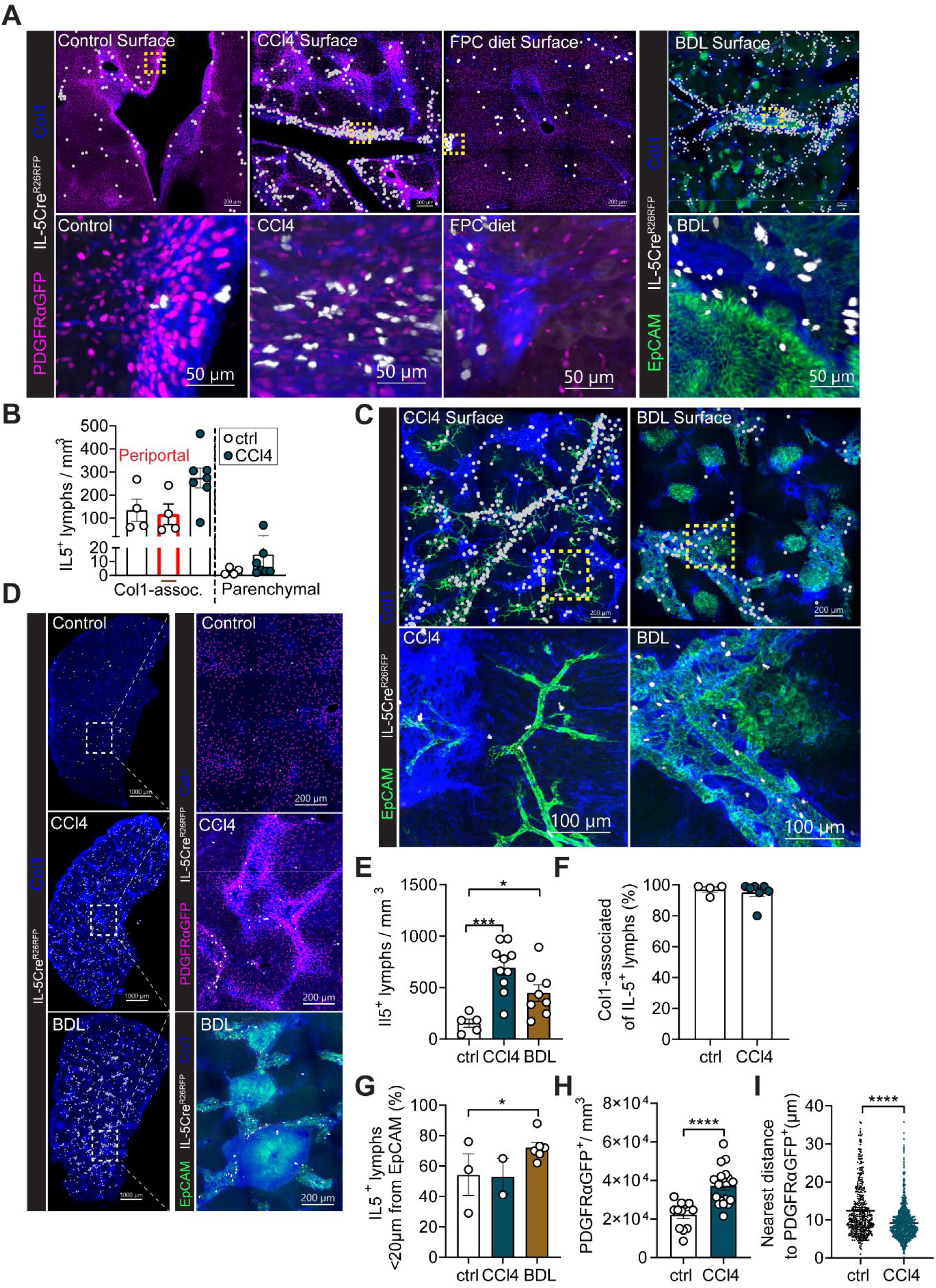
ILC2s localize to fibrotic tracts during the development of hepatic fibrosis. (A) Representative thick section confocal images or surface-rendered three-dimensional reconstruction (top row) from PDGFRαGFP; Il5tdtomato-^Cre/+;^ Rosa26^RFP/+^ mice at rest, treated with CCl_4_ for 4 weeks, fed with FPC diet for 16 weeks, or challenged with bile duct ligation (BDL), as indicated. Treatment schematics are shown in Figure S1A (0.5µl CCl_4_/g BW i.p. three times per week for 4 weeks), Figure S1G (BDL and harvest after 14 days) and Figure S1L (FPC diet for 16 weeks). All samples were prepared as coronal sections of 200µm thickness and stained for Collagen I (alpha-1 type I collagen). Images shown are representative of 3-4 mice/group, including both sexes and different littermates, with 2-3 different thick sections analyzed per mouse. (B) Quantitative analysis of liver thick-section confocal images as Collagen I-associated, periportal, and parenchymal IL-5^+^ lymphocyte numbers per tissue volume, total N = 4-7 mice/group. (C) Representative confocal liver tissue sections and surfacing analysis from IL-5^+^ lymphocyte lineage tracker mice (IL-5tdtomato-Cre; Rosa26RFP) after CCl_4_ treatment for 4 weeks or 14 days post BDL surgery. Images are representative of 3-4 mice per group. (D) Confocal thick section images of livers from PDGFRαGFP; Il5tdtomato-^Cre/+;^ Rosa26^RFP/+^ mice at rest (Control), or after 4-week CCl_4_ treatment, and from IL-5^+^ lymphocyte lineage tracker mice (IL-5tdtomato-Cre; Rosa26RFP) 14 days post BDL surgery, as indicated, with zoom of 200µm thick liver slices with IL-5^+^ lymphocytes (RFP^+^) and PDGFRαGFP^+^ fibroblasts visualized. Images are representative of 3 or more mice/group. (E) Quantification of IL-5^+^ lymphocyte numbers per tissue volume. Pooled data from 2-3 mice/group with 2-3 different thick sections analyzed per mouse. (F) Quantitative analysis of percent Collagen I-associated IL-5^+^ lymphocytes of total IL-5^+^ lymphocytes, total N = 4-7 mice/group. (G) Quantification of percent IL-5^+^ lymphocytes less than 20μm apart from epithelial duct cells (EpCAM^+^). Pooled data from 3-4mice/group with 1-3 different thick sections analyzed per mouse. (H and I) Quantitative imaging analysis of liver thick-section confocal images as PDGFRαGFP^+^ cell numbers per tissue volume (H), or distance from ILC2 to nearest PDGFRαGFP^+^ cell (I) in PDGFRαGFP; Il5tdtomato-^Cre/+;^ Rosa26^RFP/+^ mice at rest or treated with CCl_4_ for 4 weeks. Pooled data from n=3-4 mice/group with 1-3 different thick sections analyzed per mouse. Bar graphs indicate mean (±SE), unpaired t-test, *p ≤ 0.05, ***p ≤ 0.001, ****p ≤ 0.0001. See also Figure S3.

### Type 3/17 lymphocytes liver topography

As type 3/17 lymphocytes were also elevated during hepatic fibrosis (**Fig. 2I-J, Fig. S2H-I**), we determined their distribution using both RORγt-GFP (Lochner et al., 2008) and IL-17A fate mapper mice (Il17Cre; R26-fsf-TdT) (Hirota et al., 2011). In resting livers, type 3/17 lymphocytes resided in adventitial periportal areas, similar to IL-5^+^ lymphocytes (**Fig. 4A, D**). After induction of fibrosis with CCl_4_ or BDL, type 3/17 lymphocytes further accumulated in collagen dense areas of the adventitia as well as novo fibrotic tracts (**Fig. 4B-E**). In the <u>resting lung</u>, type 3/17 lymphocytes were also enriched in adventitial regions **(Fig. 4F)**, similar to IL-5^+^ type 2 lymphocytes (Dahlgren et al., 2019) but unlike IFNγ^+^ type 1 lymphocytes (Cautivo et al., 2022). Liver type 3/17 lymphocytes were comprised of γδ T cells, with additional subsets of CD4^+^ Th17 cells, ILC3s, and other innate-like T cells **(Fig. 4G-H)** and increased significantly with CCl4 fibrosis **(Fig. 4I)**. To test if AFs were sufficient to support both ILC2s and γδ T cells, we sorted lung AFs and co-cultured them with freshly sorted ILC2s or γδ T cells. AF co-cultures were sufficient to maintain both ILC2s and γδ T cells in the absence of cytokine supplement or TCR stimulation **(Fig. S3K),** as previously shown for ILC2s and IL-5^+^ Th2s (Dahlgren et al., 2019). We conclude that IL-33^high^ ‘niche’ fibroblasts are sufficient to support both ILC2s/Th2s and γδ T cells, and fibrosis-driven AF-like cell expansion *in vivo* was spatially and temporally associated with increased type 2 and type 3/17 lymphocytes.

**Figure 4:**
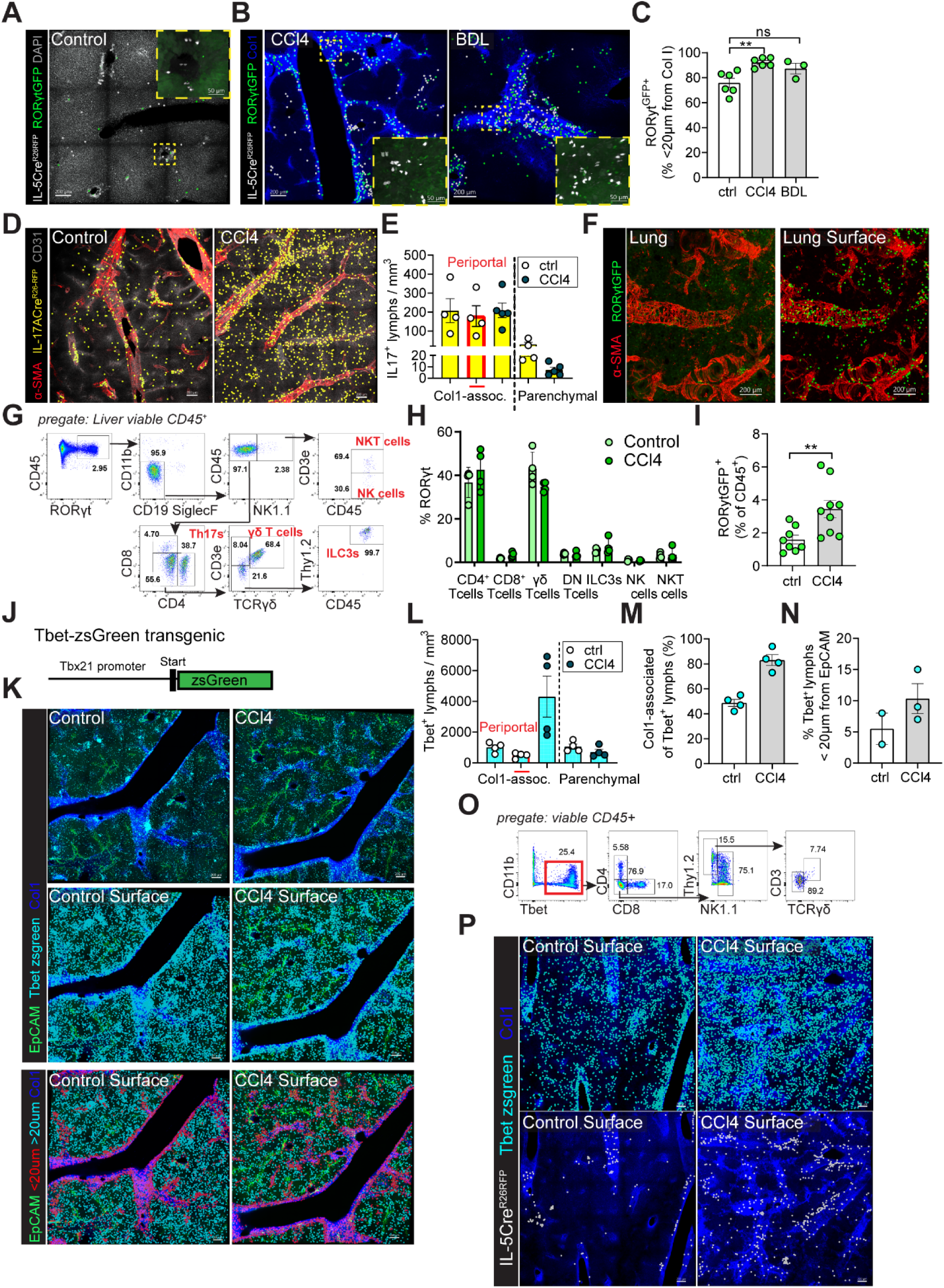
Type 3/17 lymphocytes have similar liver topography as IL-5^+^ type 2 lymphocytes. (A and B) Thick section confocal imaging from livers of RORγtGFP; Il5tdtomato-^Cre/+;^ Rosa26^RFP/+^ mice at steady state (A) or treated with CCl_4_ for 4 weeks or 14 days post bile duct ligation (BDL) surgery (B) with surfaces rendered on IL-5^+^ lymphocytes and RORytGFP^+^ cells. Higher magnification showing accumulation of RORγtGFP^+^ cells in collagen-rich regions. Image representative of 2 independent experiments with n=7-8 mice per group. (C) Quantification of percent RORγt^+^ cells less than 20μm apart from Collagen I. Pooled data from 3-4mice/group with 1-3 different thick sections analyzed per mouse. (D) Representative thick section confocal imaging of control and 4-week CCl_4_-treated IL-17A lineage tracker mice with surfaces rendered on IL-17A^+^ lymphocytes (Il17atdtomato-^Cre/+;^ Rosa26^RFP/+^). Images are representative of 2 mice with 2 sections each. (E) Quantitative analysis of liver thick-section confocal images as Collagen I-associated, periportal, and parenchymal IL-17^+^ lymphocyte numbers per tissue volume, total N = 4-5 mice/group. (F) Representative thick section confocal imaging of lung from RORγtGFP reporter mice at steady state. Image is representative of 3 mice with 2 sections each. (G) Representative flow gating scheme for liver RORγtGFP^+^ cells from RORγtGFP reporter mice treated with CCl_4_ for 4 weeks, including pre-gating strategy. (H) Flow cytometry quantification of percentages of RORγt expression in CD4^+^ T cells, γδ T cells, DN T cells, ILC3s, CD8^+^ T cells, NK cells, and NKT cells from RORγtGFP reporter mice treated with vehicle or CCl_4_ for 4 weeks. (I) Flow cytometry quantification of RORytGFP^+^ cells from livers of RORγtGFP reporter mice treated with vehicle or CCl_4_ for 4 weeks. Pooled data from 2 independent experiments with 6-12 weeks-old age-and sex-matched mice. (J) Schematic of Tbx21-ZsGreen “T-bet” reporter mouse, relevant to I-N. (K) Representative confocal thick section image from liver of T-bet (Tbx21)-zsGreen transcription factor reporter mice treated with vehicle or CCl_4_ for 4 weeks and stained with EpCAM and Collagen I. Images shown are representative of 2 mice with 2 sections each. (L) Quantitative analysis of liver thick-section confocal images as Collagen I-associated, periportal, and parenchymal Tbet^+^ lymphocyte numbers per tissue volume, total N = 4 mice/group. (M and N) Quantitative analysis of percent Collagen I-associated Tbet^+^ lymphocytes of total Tbet^+^ lymphocytes (M) and quantification of percent T-bet^+^ cells less than 20μm apart from epithelial duct cells (EpCAM^+^) (N). Pooled data from 4 mice/group. (O) Representative flow gating scheme for liver zsGreen expression in T-bet-zsGreen reporter mice treated with CCl_4_ for 4 weeks, including pre-gating strategy. (P) Representative confocal thick section image from liver of T-bet-zsGreen; Il5tdtomato-^Cre/+;^ Rosa26^RFP/+^ mice treated with vehicle or CCl_4_ for 4 weeks and stained with Collagen I. Images shown are representative of 2 mice with 2 sections each. Bar graphs indicate mean (±SE), unpaired t-test, ns= not significant, *p ≤ 0.05, **p ≤ 0.01.

To compare these findings with the topography of fibrosis-associated type 1 lymphocytes, we imaged Tbet (Tbx21) transgenic reporters, which mark cells with capacity to produce IFNγ **(Fig. 4J)** (Zhu et al., 2012). In contrast to type 2 and type 3/17 lymphocytes, type 1 lymphocytes are associated with <u>resistance</u> to NASH and liver fibrosis (Hart et al., 2017). At rest, type 1 lymphocytes (Tbet-zsgreen^+^) were more abundant than type 2 or type 3/17 lymphocytes and were broadly distributed across parenchymal sinusoidal and collagen-dense adventitial regions (Cautivo et al., 2022), with some enrichment near pericentral hepatic veins (**Fig. 4K, L**). CCl_4_-induced liver injury drove modest type 1 cell expansion and increased localization to collagen-rich regions (**Fig. 4K-M, P**); however, in contrast to type 2 or type 3/17 lymphocytes (**Fig. 3G**), few type 1 lymphocytes localized near periportal zones at rest or after fibrosis (5-10%; **Fig. 4N**). Flow cytometry confirmed that naïve liver Tbet^zsGreen+^ cells were a mixture of NK cells and ILC1s, as well as subsets of CD4^+^ and CD8^+^ T cells and innate-like T cells (**Fig. 4O**). This data, together with previous work (Cautivo et al., 2022), suggest the distribution of bulk liver type 1 lymphocytes is distinct from that of type 2 and type 3/17 lymphocytes, both at rest and in models of liver fibrosis.

### Loss of IL-5^+^ lymphocytes worsened hepatic fibrosis

Next we tested the function of IL-5^+^ lymphocytes during the development of liver fibrosis. Given the involvement of IL-13 in type 2 immune-driven fibrosis (Gieseck et al., 2016), we predicted type 2 lymphocyte deficiency would offer protection from liver fibrosis. Using IL-5 deleter mice, a well-established genetic approach that sensitively and specifically deletes most ILC2s and rare IL-5^+^ Th2 cells (Il5Cre; R26-DTA mice) (Van Dyken et al., 2014), we confirmed efficient deletion of IL-5^+^ lymphocytes, predominantly ILC2s, both at rest and after fibrosis (∼80% deletion, **Fig. 5A-C)**. Of note, the Il5Cre allele disrupts the endogenous *Il5* locus, and both deleter mice and littermate controls similarly lack functional IL-5. Unexpectedly, depletion of IL-5^+^ lymphocytes resulted in elevated liver CCl_4_ fibrosis, including increased hydroxyproline, liver profibrotic gene expression, periportal ductal expansion, collagen tract deposition, and CD34 expression, a marker of periportal liver fibroblasts (Ramachandran et al., 2019; Dobie et al., 2019b), whereas serum ALT (acute liver damage) was not different between groups (**Fig. 5D-I, Fig S4A**). IL-5 deleter mice had decreased pericentral hepatocyte markers (Cyp2E1 and glutamin synthetase (GS), **Fig. S4B, C**), consistent with more severe CCl_4_-induced pericentral loss and fibrosis. These results were recapitulated in an inducible model that allowed for depletion of IL-5^+^ lymphocytes in adult mice (Il5Cre^DTR^), with efficient and specific ILC2 deletion **(Fig. 5J, Fig. S4D-F)**. Fibrosis was also elevated (hydroxyproline levels), whereas weight loss and relative CD4 T cells and NK cells did not change **(Fig. 5K, Fig. S4G-I)**. To compare these results with a model of cholestasis-induced liver injury, we next performed BDL surgery. IL-5 deleter mice were similar in survival, weight loss, ALT levels and bilirubin levels **(Fig. 5L, Fig. S4J-M)**. However, they also demonstrated enhanced liver fibrosis, with elevated hydroxyproline levels, Sirius red and trichrome staining, and collagen type 1 staining (**Fig. 5M-P)**. Together, these findings suggest that loss of IL-5^+^ lymphocytes worsens liver fibrosis in both CCl_4_ and BDL surgery models, consistent with protective role(s) for type 2 lymphocytes such as ILC2s.

**Figure 5:**
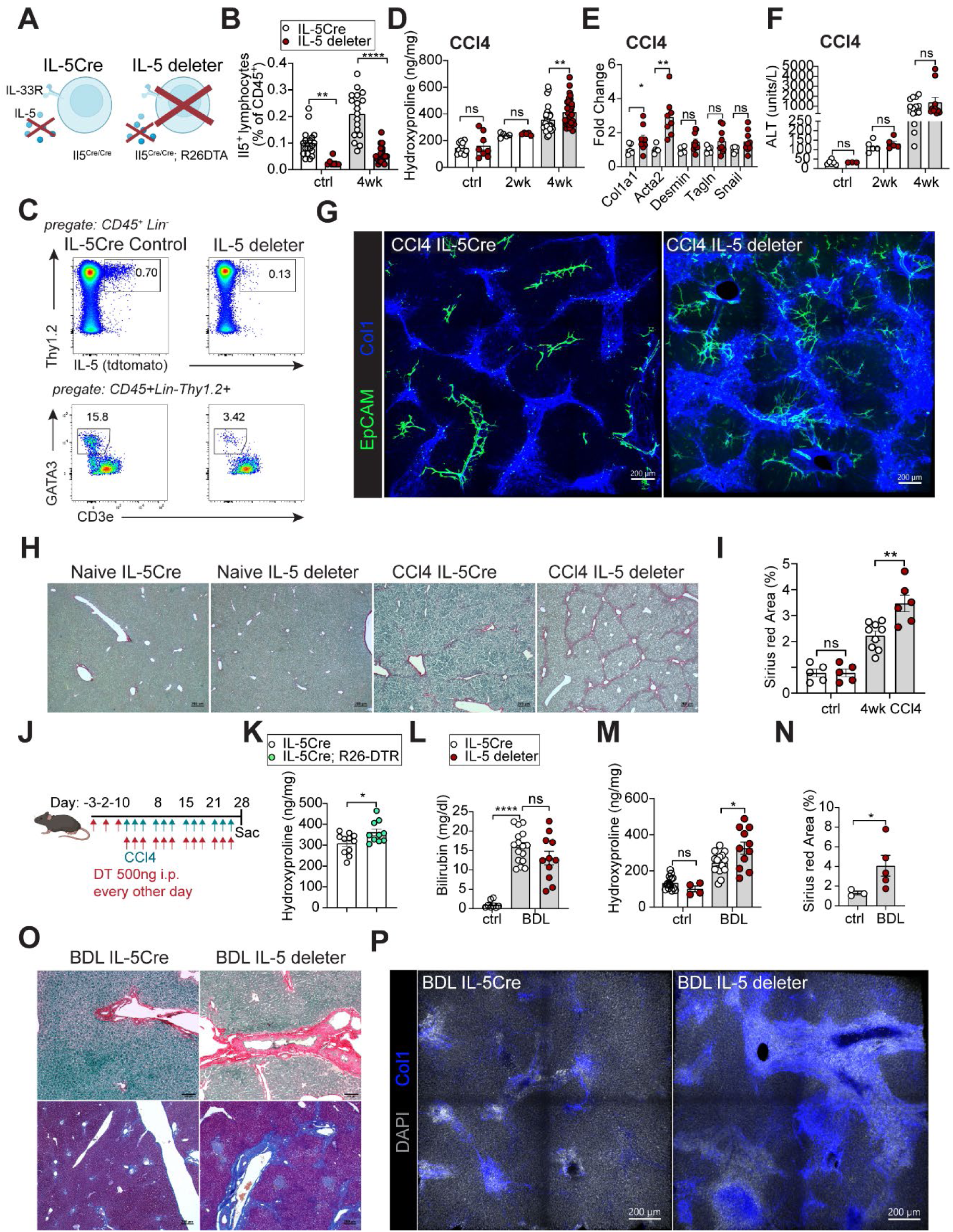
ILC2s regulate collagen deposition in different models of hepatic fibrosis. (A) Schematic of IL-5 deleter mice (RRDD; *Il5*^Red5/Red5^*Gt (Rosa)26*^DTA/DTA^) and IL-5tdtomato-Cre controls. (B) Flow cytometry quantification, showing percentage of liver IL-5^+^ lymphocytes from control or CCl_4_-treated Il5-tdtomato-Cre mice and IL-5 deleter (RRDD; *Il5*^Red5/Red5^*Gt (Rosa)26*^DTA/DTA^) mice. Pooled data from 3 independent experiments with n=5-8 8-10 weeks-old age-and sex-matched mice/group. (C) Representative flow cytometry plots of IL-5^+^ lymphocytes (Lin^-^Thy1.2^+^Red5^+^) and ILC2s (Lin^-^Thy1.2^+^GATA3^+^) in liver from either IL-5tdtomato-Cre or IL-5 deleter mice treated with CCl_4_ for 4 weeks. (D) Quantification of Hydroxyproline levels of the liver from control or 2-and 4-week CCl_4_-treated Il5-tdtomato-Cre mice and IL-5 deleter (RRDD; *Il5*^Red5/Red5^*Gt (Rosa)26*^DTA/DTA^) mice. Pooled data from 3 independent experiments from 8-10 weeks-old age-and sex-matched mice. (E) Total liver gene expression of *col1a1, acta2, desmin, tagln,* and *snail1*, normalized to *gapdh* expression in control or 4-week CCl_4_-treated Il5-tdtomato-Cre mice and IL-5 deleter (RRDD; *Il5*^Red5/Red5^*Gt (Rosa)26*^DTA/DTA^) mice. Pooled data from 2 independent experiments from 8-10 weeks-old age-and sex-matched mice. (F) Quantification of ALT levels of the liver from control or 2-and 4-week CCl_4_-treated Il5-tdtomato-Cre mice and IL-5 deleter (RRDD; *Il5*^Red5/Red5^*Gt (Rosa)26*^DTA/DTA^) mice. Pooled data from 3 independent experiments from 8-10 weeks-old age-and sex-matched mice. (G) Representative confocal liver tissue sections from control or 4-week CCl_4_-treated Il5-tdtomato-Cre mice and IL-5 deleter (RRDD; *Il5*^Red5/Red5^*Gt (Rosa)26*^DTA/DTA^) mice with staining for EpCAM (epithelial duct cells) and Collagen I. Images shown are representative of 3-4 mice/group. (H and I) Representative Sirius red staining images of 5µm paraffin liver sections (H) and quantification of Sirius red staining (I) from Il5-tdtomato-Cre and IL-5 deleter mice after 4-week CCl_4_ treatment and controls. (J) Schematic showing CCl_4_ administration schedule in IL-5^DTR^ (*Il5*^Red5/Red5^*Gt (Rosa)26*^DTR/DTR^ mice) injected intraperitoneally (i.p.) with 0.5µl CCl_4_/g BW three times per week 4 weeks, and with 500ng diphteria toxin (DT) every other day throughout the experiment, relevant to K. (K) Hydroxyproline levels of 4-week CCl_4_-treated IL-5tdtomato-Cre and IL-5^DTR^ mice. Pooled data from 2 independent experiments from 8-10 weeks-old age-and sex-matched mice. (L and M) Quantification of Bilirubin (L) and Hydroxyproline levels (M) for Il5-tdtomato-Cre mice and IL-5 deleter mice 14 days post BDL. Data are pooled from 3 independent experiments with n=5-13 8-10 weeks-old age-and sex-matched mice/group. (N and O) Quantification of Sirius red staining (N) and representative Sirius red staining (O) from Il5-tdtomato-Cre mice and IL-5 deleter mice 14 days post BDL. Images are representative of 3 independent experiments with n=5-13 mice per group. (P) Representative confocal liver tissue sections from Il5-tdtomato-Cre mice and IL-5 deleter mice 14 days post BDL. Samples were prepared as coronal sections of 200µm thickness and stained for Collagen I (a-Collagen I). Bar graphs indicate mean (±SE), unpaired t-test, *p ≤ 0.05, **p ≤ 0.01, ****p ≤ 0.0001. See also Figure S4.

### Neither ILC2 expansion, IL-33, nor IL-4/IL-13 signalling impacted degree of hepatic fibrosis

Given that fibrosis induced by CCl_4_ treatment and BDL surgery were impacted by IL-5^+^ type 2 lymphocytes, we next examined whether IL-33 played a role. IL-33 treatment drove liver ILC2 accumulation, as expected **(Fig. S5A-C)**. However, the pre-treatment with IL-33 and ILC2 expansion did not alter the subsequent course of CCl_4_-fibrosis, with similar levels of ALT, hydroxyproline, liver Tregs, and CD8^+^ T cells **(Fig. S5D-I)**. We also tested the role of endogenous IL-33, treating IL33-deficient mice (Vainchtein et al., 2018a) and IL-33 heterozygous littermates with CCl_4_ for 4 weeks. IL-33 deficient mice showed no significant difference in fibrosis or damage markers, and liver ILC2s were increased with fibrosis but unchanged with loss of IL-33 **(Fig. S5J-N, R)**. Similarly, IL-33 deficiency did not alter BDL-induced liver disease/fibrosis **(Fig. S5O-R)**. We conclude that loss of IL-33 does not alter CCl_4_-or BDL-related liver fibrosis or impair accumulation of liver ILC2s; further, driving ILC2 expansion did not alter subsequent liver fibrosis.

ILC2s produce constitutive IL-5, and produce IL-13 downstream of multiple activating signals (Nussbaum et al., 2013; Molofsky et al., 2013; Van Dyken et al., 2014), which can regulate distinct aspects of liver fibrosis, steatosis, cholestasis, and ductular reaction in type 2 cytokine-driven models of liver fibrosis (Gieseck et al., 2016). To directly test the possible role of IL-13 in liver fibrosis in our models, we first induced IL-13 overexpression (hydrodynamic tail vein injection, 10µg IL-13 overexpression plasmid IL-13OP) (Gieseck et al., 2016). This resulted in an inflammatory reaction with increased total liver leukocytes **(Fig. S6A-E)** and elevated collagen 1 **(Fig. S6F, G)**, consistent with prior work (Gieseck et al., 2016). We recapitulated these results with mice subjected to a lower dose of IL-13 overexpression **(Fig. S6H-I)**, although IL-5^+^ lymphocytes were not significantly elevated with this regimen **(Fig. S6J-K)**. As such, IL-13 excess is sufficient to drive liver inflammation and fibrosis. However, this did not address the role of endogenous IL-4/IL-13 signalling. As such, we tested liver fibrosis in mice with global loss of IL-4Ra, which is a required co-receptor component for both IL-4 and IL-13 (Mohrs et al., 1999). Following CCl_4_ treatment, there was no difference in liver fibrosis in IL4Ra^-/-^ mice relative to sex-and age-matched controls; similar to IL-33-deficient mice, we also did not find altered numbers of liver ILC2s **(Fig. S6L-M, P)**. BDL-induced fibrosis yielded similar results, with no alteration in survival/weight, liver damage, or fibrosis **(Fig. S6N-S**). These data suggest that neither IL-33 nor IL-4/IL-13 signalling are required for the development of hepatic fibrosis during CCl_4_ treatment or BDL surgery, although overexpression of IL-13 is sufficient to drive severe inflammation and multi-organ fibrosis.

### IL-5^+^ type 2 lymphocytes regulate the type 3/17 immune response in advanced hepatic fibrosis

To understand how type 2 lymphocytes impact liver fibrosis, we examined the liver fibrosis-associated immune landscape in the presence or absence of IL-5^+^ lymphocytes **(Fig. 6A)**. During CCl_4_-induced fibrosis, IL-5^+^ lymphocyte deficiency was accompanied by a further increase in γδ T cells; these results were recapitulated with inducible IL-5 lymphocyte depletion (Il5Cre^DTR^) **(Fig. 6B-E)**. Moreover, γδ T cells shifted towards an increased type 3/17 phenotype, including increased RORγt and IL-17A capacity **(Fig. 6F-H)**. In contrast, we found that loss of IL-5^+^ lymphocytes did not consistently change CD4^+^ IFNγ^+^ Th1 cells, IL-13^+^ Th2 cells, IL-17A^+^ Th17 cells, B cells, macrophages, CD8^+^ T cells, or neutrophils **(Fig. 6F, 6I-S)**. After BDL surgery, γδ T cells and their RORγt expression were also further increased in IL-5 deleter mice **(Fig. 6T-W)**. These data suggest a critical role of IL-5^+^ lymphocytes in restraining type 3/17 immunity during hepatic fibrosis.

**Figure 6:**
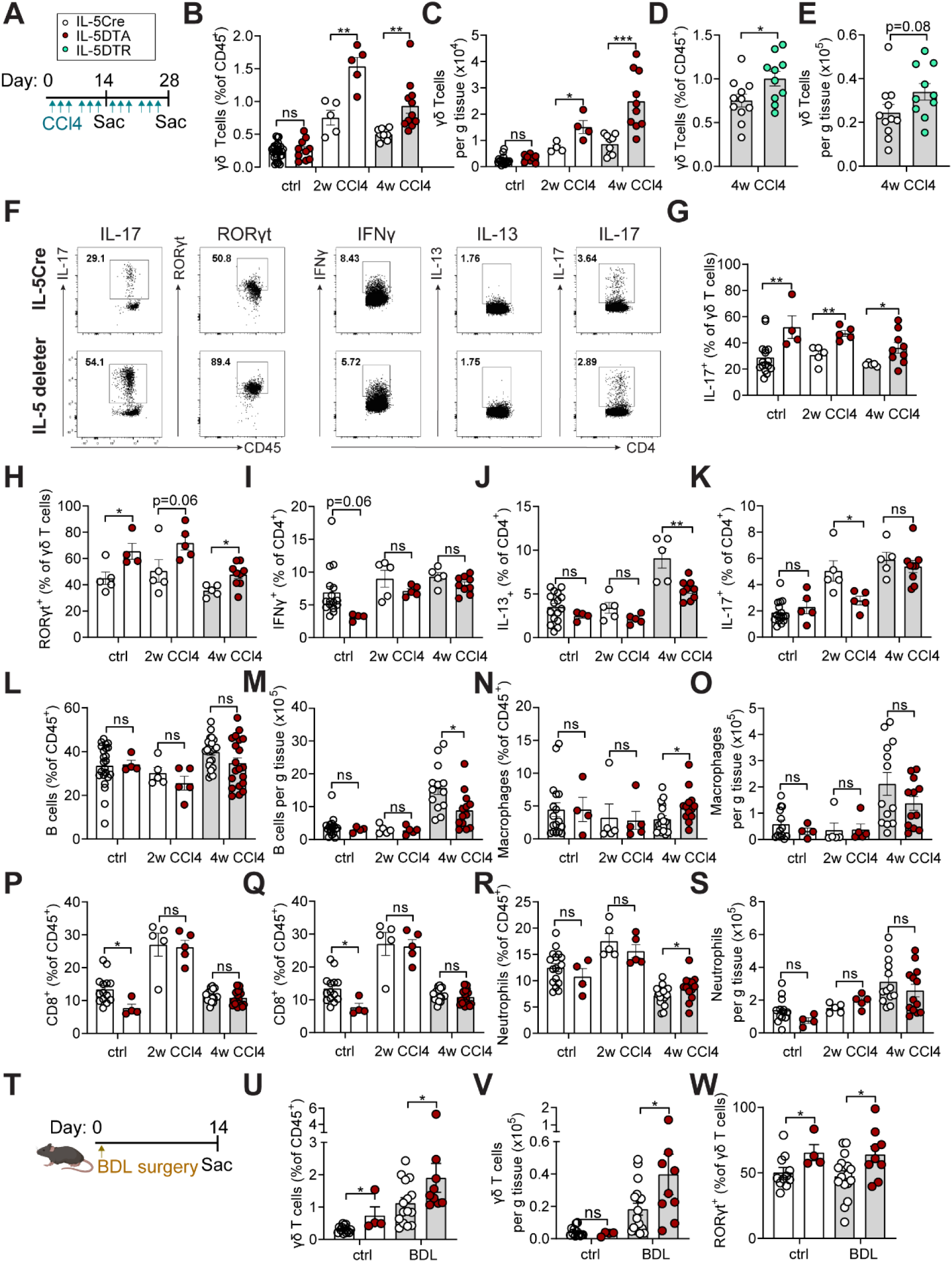
ILC2s regulate the type3/17 immune response in advanced hepatic fibrosis. (A) Schematic showing CCl_4_ administration schedule in Il5-tdtomato-Cre mice and IL-5 deleter (RRDD; *Il5*^Red5/Red5^*Gt (Rosa)26*^DTA/DTA^) mice injected intraperitoneally (i.p.) with 0.5µl CCl_4_/g BW three times per week for 2 or 4 weeks (2w, 4w), relevant to B, C and G-S. (B and C) Flow cytometry quantification, showing percent (B) and total numbers of γδ T cells (C) in livers from Il5-tdtomato-Cre mice and IL-5 deleter (RRDD; *Il5*^Red5/Red5^*Gt (Rosa)26*^DTA/DTA^) mice treated with vehicle or CCl_4_ for the indicated duration. Pooled data from 3 independent experiments from 8-10 weeks-old age-and sex-matched mice. (D and E) Flow cytometry quantification, showing percent (D) and total numbers of γδ T cells (E) in livers from Il5-tdtomato-Cre mice and IL-5^DTR^ (*Il5*^Red5/Red5^*Gt (Rosa)26*^DTR/DTR^ mice treated with CCl_4_ for the indicated duration. Mice were injected with 500ng diphteria toxin (DT) every other day as indicated in Supplementary Figure 4. Pooled data from 2 independent experiments from 8-10 weeks-old age-and sex-matched mice. (F-K) Representative flow cytometric plots and quantitation, showing cytokine expression of IL-17 (F, G) and RORγt (F, H) hepatic γδ T cells, and expression of IFNγ (F, I), IL-13 (F, J), and IL-17 (F, K) in hepatic CD4^+^ T cells from Il5-tdtomato-Cre mice and IL-5 deleter mice treated with vehicle or CCl_4_ for the indicated duration after stimulation with ionomycin (Ion) and phorbol 12-myristate 13-acetate (PMA). (L-S) Flow cytometry quantification of percent (L) or total numbers (M) of B cells, percent (N) or total numbers (O) of macrophages, percent (P) or total numbers (Q) of CD8^+^ T cells, and percent (R) or total numbers (S) of neutrophils in livers from Il5-tdtomato-Cre mice and IL-5 deleter mice treated with vehicle or CCl_4_ for the indicated duration. Pooled data from 3 independent experiments from 8-10 weeks-old age-and sex-matched mice. (T) Schematic showing schedule for bile duct ligation (BDL) surgery in Il5-tdtomato-Cre mice and IL-5 deleter mice. (U-W) Flow cytometry quantification of percent (U) or total numbers (V) of γδ T cells and percent of RORyt-expressing γδ T cells (W) in livers from Il5-tdtomato-Cre mice and IL-5 deleter (RRDD; *Il5*^Red5/Red5^*Gt (Rosa)26*^DTA/DTA^) mice 14 days post BDL surgery. Pooled data from 3 independent experiments with 8-10 weeks-old age-and sex-matched mice. Bar graphs indicate mean (±SE). unpaired t-test, *p ≤ 0.05, **p ≤ 0.01, ***p ≤ 0.001.

### Redundancy in type 3/17 lymphocytes during hepatic fibrosis

We next tested whether mice deficient in γδ T cells (*TCRd^-/-^*) would be protected against CCl_4_ driven hepatic fibrosis **(Fig. S7A-E)**. While T*CRd*^-/-^ mice had alower ALT levels **(Fig. S7F), neither** liver fibrosis nor ILC2s were altered **(Fig. S7G-I**), suggesting loss of γδ T cells alone was not sufficient to impact CCl_4_-induced liver fibrosis. However, levels of other IL-17A producing innate-like DN T cells were increased in CCl_4_-treated T*CRd*^-/-^ mice **(Fig. S7J)**, suggesting possible type 3/17 lymphocyte compensation in the absence of γδ T cells (e.g., MAIT cells, ILC3s). RORγt regulates the differentiation of type 3/17 immune cells and IL-17A production (Ivanov et al., 2006) and constitutes a therapeutic target to treat Th17-related autoimmune diseases (Withers et al., 2016). Antagonizing RORγt transcriptional activity could therefore limit excessive IL-17 responses associated with loss of IL-5^+^ lymphocytes. To test this, we treated IL-5 deleter mice daily with a RORγt antagonist (GSK 805), starting after 2 weeks of CCl_4_ treatment **(Fig. S7K)**. However, liver damage or fibrosis was not reduced, although Th17 frequencies were mildly decreased **(Fig. S7L-P)**. IL-17A^+^ unconventional DN T cells and γδ T cells were notably and surprisingly increased **(Fig. S7Q-T),** again suggesting potential compensation. To directly test the role of the IL-17A cytokine itself, we administered an anti-IL-17A neutralizing antibody; this led to reduction of the elevated neutrophil frequencies in IL-5 deleter mice, but IL-17 producing γδ T cells were further elevated and damage and fibrosis markers were comparable between the groups **(Fig. S7U-Y)**.

Our data suggested redundancy in type 3/17 lymphocytes and their signals that may mediate liver fibrosis, at least in the context of type 2 lymphocyte deficiency. To overcome this limitation and bypass compensatory feedback mechanisms, we next used a genetic approach to concurrently reduce all IL-17A^+^ type 3/17 lymphocytes with IL-5^+^ type 2 lymphocytes, using IL-17A^Cre^ mice (Hirota et al., 2011) intercrossed with IL-5 deleter mice (*Il5*^Cre/Cre^*; R26*^DTA/DTA^), hereafter referred to as ‘double deleter’ mice **(Fig. 7A)**. We subjected double deleter mice, and their IL-5 deleter controls, to 4 weeks of CCl_4_ treatment **(Fig. 7B)**. Liver fibrosis and inflammation markers were reduced as compared to IL-5 deleter only mice **(Fig. 7C-F),** as were total liver RORγt^+^ cells, neutrophils, and γδ T cells **(Fig. 7G-K)**. Together, our findings suggest a model in which type 2 and type 3/17 lymphocytes share resting and expanded post-fibrotic liver niches, defined at least in part by IL-33^high^ niche fibroblasts, and loss of type 2 lymphocytes leads to aggravated type 3/17 driven inflammation and fibrosis.

**Figure 7:**
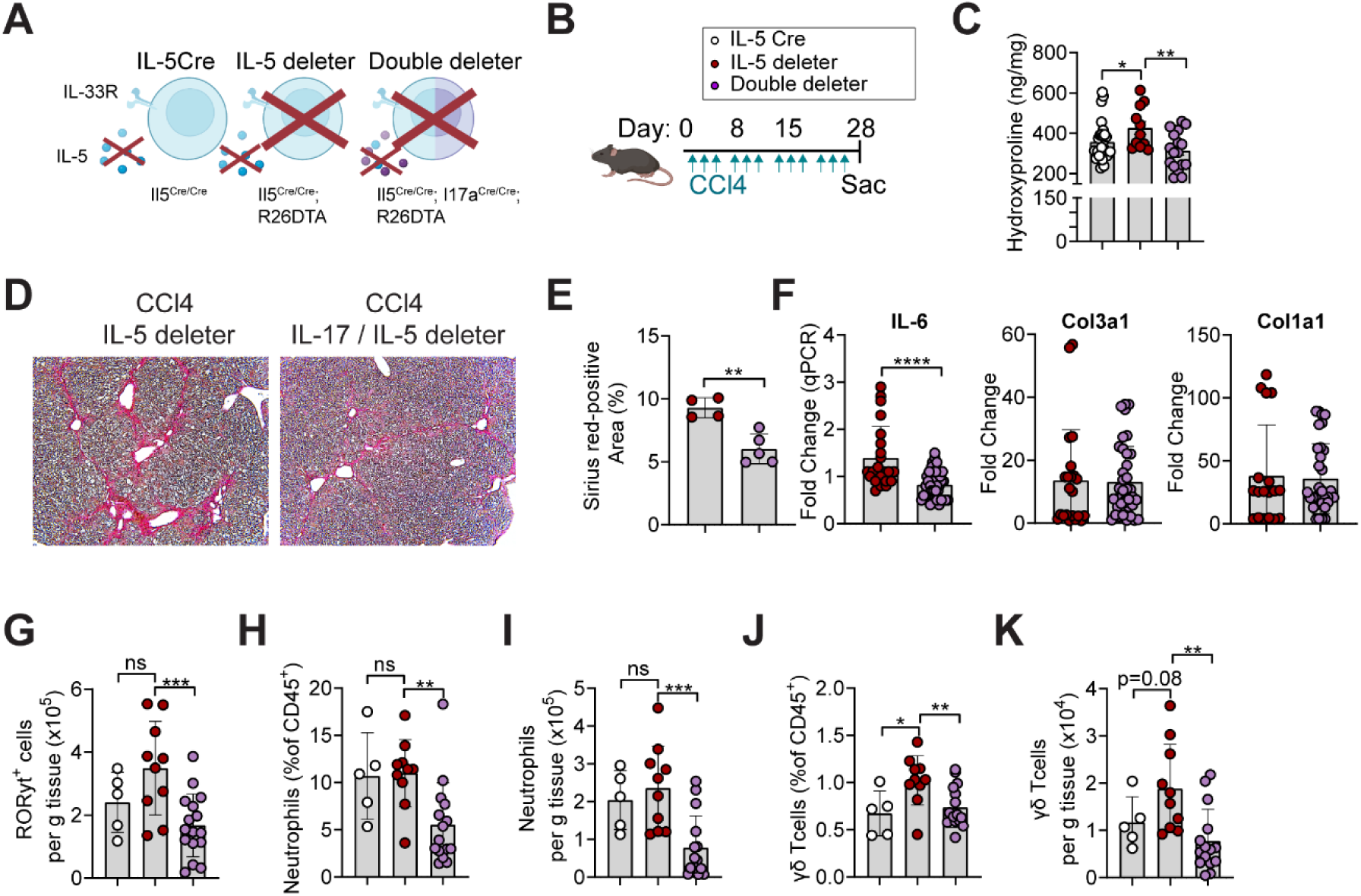
Depletion of IL-17 reduces advanced hepatic fibrosis. (A) Schematic of IL-17 IL-5 deleter mice (Il17aCre; *Il5*Cre; *(Rosa)26*^DTA/DTA^) and IL-5 deleter mice (RRDD; *Il5*^Red5/Red5^*Gt (Rosa)26*^DTA/DTA^) as controls. (B) Schematic showing treatment strategy for CCl_4_ injected intraperitoneally (i.p.) with 0.5µl CCl_4_/g BW three times per week for 4 weeks with Il5-tdtomato-Cre mice, IL-5 deleter (RRDD; *Il5*^Red5/Red5^*Gt (Rosa)26*^DTA/DTA^), and IL-17A IL-5 deleter mice (*Il17aCre; Il5Cre; (Rosa)26*^DTA/DTA^) relevant to data in (C-K). (C) Quantification of hepatic Hydroxyproline in Il5-tdtomato-Cre mice, IL-5 deleter mice, and IL-17A IL-5 deleter mice. Data pooled from 2 independent experiments with 8-10 weeks-old age-and sex-matched mice. (D and E) Representative Sirius red staining images of 5µm paraffin liver sections (D) and quantification of Sirius red staining (E) from Il5-tdtomato-Cre, IL-5 deleter mice, and IL-17A IL-5 deleter mice after 4-week CCl_4_ treatment. Images are representative of 2 independent experiments with 8-10 weeks old age-and sex-matched mice. (F) Total liver gene expression of IL6, Col1a1 and Col1a3 normalized to expression of housekeeping gene 36B4 in 4-week CCl_4_-treated IL-5 deleter mice and IL-17A IL-5 deleter mice. Pooled data from 2 independent experiments from 8-10 weeks-old age-and sex-matched mice (G-K) Flow cytometry quantification, showing total numbers of RORγt cells (G), percent (H) and total numbers (I) of neutrophils, and percent (J) and total numbers (K) of γδ T cells in livers from Il5-tdtomato-Cre mice, IL-5 deleter mice, and IL-17A IL-5 deleter mice treated with CCl_4_ for 4 weeks. Pooled data from 2 independent experiments from 8-10 weeks-old age-and sex-matched mice. Bar graphs indicate mean (±SE). Unpaired t-test for G and H, and one-Way ANOVA with multiple measurement for E and I-M, ns= not significant, **p ≤ 0.01, ***p ≤ 0.001, ****p ≤ 0.0001. See also Figure S7.

## DISCUSSION

Fibrosis is caused by excessive, chronic accumulation of collagen and other extracellular matrix components, typically driven by dysregulated tissue repair during repetitive perturbations. Although lymphocytes and their cytokines influence tissue fibrosis, the precise temporal and spatial contributions are poorly defined. We previously identified adventitial niches around larger vessels and other sterile tissue border and boundary sites, where IL-33^high^ fibroblasts locally regulate IL-5^+^ type 2 lymphocytes (Dahlgren and Molofsky, 2019; Dahlgren et al., 2019; Cautivo et al., 2022). Additional immune and non-immune subsets are also enriched at these regional immune hubs (Cautivo et al., 2020; Sbierski-Kind et al., 2021; Rustenhoven et al., 2021). Adventitial fibroblasts across multiple tissues share a universal fibroblast signature, defined by expression of Dpt and Pi16, with increased mesenchymal progenitor capacity suggesting a potential broadly conserved immunomodulatory role (Buechler et al., 2021). Here we found chronic liver injury and fibrosis cause expansion of AF-like IL-33^high^ fibroblasts that are tightly associated with expanded type 2 and type 3/17 lymphocytes. IL-33^high^ fibroblasts were distinct from αSMA^+^ and TGFβ-driven profibrotic ‘myofibroblasts’, although they co-existed within collagen-dense fibrotic areas. Fibrosis expanded IL-5^+^ lymphocytes that limited type 3/17 immune accumulation and response across multiple mouse models of liver fibrosis, and loss of type 2 lymphocytes worsened fibrosis that was dependent on the presence of increased type 3/17 lymphocytes.

Chronic and inappropriate activation of type 2 immunity drives allergic disease, but it can also synergize with fibrotic signals such as TGFβ to promote pathologic fibrosis. In type 2 immune-skewed settings, IL-13 drives hepatic fibrosis, steatosis, cholestasis and ductular reaction, simultaneously acting on distinct cellular targets (Gieseck et al., 2016). Excess IL-33, IL-25 and TSLP, additional tissue cytokines that drive type 2 immunity, can also lead to organ fibrosis (Hams et al., 2014; Kim et al., 2013; Mchedlidze et al., 2013), with combinatorial targeting of all three signals reducing type 2-driven inflammation and fibrosis in infectious models (Vannella et al., 2016; Van Dyken et al., 2014). In contrast, we found that limited exposure to IL-33, with concomitant ILC2 expansion, was not sufficient to alter CCl_4_-driven liver fibrosis; further, while we confirmed that IL-13 excess was sufficient to drive fibrosis (IL-13 OEP), lack of IL-4Rα did not alter liver fibrosis in CCl_4_-models. Type 2 immune activation can also represent an adaptive response to tissue perturbation, promoting resolution of inflammation and protecting against lung injury (Van Dyken and Locksley, 2013; Karo-Atar et al., 2016; Wilson et al., 2010; Van Dyken et al., 2014). As such, one parsimonious conclusion is that while excessive type 2 immunity can promote fibrotic disease, its role may be context and timing dependent.

Elevated type 3/17 lymphocytes and associated IL-17A and IL-22 have emerged as potential drivers of fibrosis in chronic or repetitive liver injuries (Fabre et al., 2018; Meng et al., 2012). IL-17A has profibrogenic functions through increased production of TGFβ (Tan et al., 2013), recruitment of pro-inflammatory neutrophils and monocytes, and activation of the STAT3 signaling that induces type 1 collagen production in liver stromal cells (Fabre et al., 2018). IL-17A can also enhance the sensitivity of fibroblasts to TGFβ by increasing the expression of TGFβRII (Fabre et al., 2014). These observations are consistent with our findings that IL-17A^+^ lymphocytes accumulated in collagen-dense fibrous bands and correlated with disease severity, and their loss attenuated the excessive hepatic fibrosis observed with loss of type 2 lymphocytes. Targeting the type 3/17 immune axis may represent a viable strategy in various liver diseases and clinical trials to evaluate the antifibrotic effects of IL-17 blocking antibody are underway (Magdaleno-Tapial et al., 2021).

Aside from lymphocytes, macrophages coordinate tissue response to injury and contribute both to fibrosis progression and resolution (Fabre et al., 2022; Hart et al., 2017; Guilliams et al., 2022; Remmerie et al., 2020). Both resident and recruited macrophages localize to distinct liver zones in contact with liver sinusoidal endothelial cells and hepatic stellate cells. Profibrotic CD9^+^ TREM2^+^ scar-associated macrophages (SAMs) were enriched in the fibrotic niche adjacent to activated stromal cells (Fabre et al., 2022), suggesting distinct macrophage states may be an additional critical fibrotic niche component. Moreover, commensal bacterial signals also organize liver immune spatial polarization, with Kupffer cells and NKT cells enriched in parenchymal zones closer to portal areas promoting host defense (Gola et al., 2020). It will be interesting to investigate how the microbiome influences immune zonation in the liver, including distribution of lymphocyte subsets, and to reassess the relationship between the localization of immune cells, host protection, and their impact on both tissue physiology and fibrosis.

Our results show a shared stromal niche where IL-5^+^ and IL-17^+^ innate lymphocytes are in close proximity, with possible similar niches in other tissues. In contrast, type 1 lymphocytes (Tbet^+^) were broadly distributed across domains both at rest and in fibrotic livers, and consistent with prior work (Cautivo et al., 2022). IL-5^+^ and IL-17^+^ lymphocytes accumulated in periportal regions and fibrous bands after CCl_4_-or BDL-induced liver injury, but they did not significantly accumulate in liver parenchymal sites, a region with abundant type 1 lymphocytes; nonetheless, it is possible that subsets of type 1 lymphocytes are topographically similar to ILC2s and γδ T cells and could locally impact this fibroblast-immune crosstalk. We also found that IL-5^+^ lymphocytes restrained type 3/17-associated responses to chronic liver injury. Similar results were previously seen in chitin-induced type 2 lung inflammation (Van Dyken et al., 2014), and IL-33R^+^ type 2-like Tregs suppress innate IL-17A^+^ γδ T cell responses to mucosal injury induced by environmental allergens (Faustino et al., 2020) and in mouse models of neuroinflammation (Hemmers et al., 2021). The cross-regulatory activity between flavors of cell-mediated effector immunity is well recognized, but the understanding of the complex cell-extrinsic niches that may regulate lymphocyte residence and accumulation is only emerging (Choy et al., 2015; Gieseck et al., 2018). While IL-33^high^ adventitial fibroblasts are sufficient to support both ILC2 and γδ T cell survival, we did not observe differences in their liver accumulation in the absence of IL-33. Early life ILC2s also associate intimately with AFs that produce both TSLP and IL-33, but neither signal was required for ILC2 tissue colonization or identity (Ricardo-Gonzalez et al., 2018). Further work is needed to understand these fibroblast-type 2 – type 3/17 cellular interactions, but these results suggest additional cells and signals are likely involved.

We found that both healthy and fibrotic livers contained a subset of fibroblasts that expressed high IL-33 and were enriched in periportal regions and fibrous bands. Similar AFs, that are primed to support type 2 immune responses (Dahlgren and Molofsky, 2019), were recently referred to as ‘border fibroblasts’ since they are located at the adventitial domain surrounding larger vessels and body-cavity linings and, thus, help maintain tissue zonation (Boothby et al., 2021). In contrast collagen-high myofibroblasts showed only weak or absent expression of IL-33. Similarly, treatment of AFs in vitro with the pro-fibrotic mediator TGF-β led to downregulation of AF-expressed *Il33* and associated immunomodulatory cytokines and receptors suggesting IL-33^+^ immunomodulatory fibroblasts enriched in adventitial niches represent largely distinct cell states from SMA^+^ stromal cells that classically associate with tissue fibrosis. scRNAseq of healthy and fibrotic mouse livers has revealed spatial zonation of pericyte-like HSCs (Dobie et al., 2019b). Although we isolated HSCs from fibrotic livers by flow cytometry, we were unable to dissect IL-33 expression in distinct HSC subsets and to verify their role in contributing to advanced hepatic fibrosis. Our work does not resolve the possible lineage relationships between HSCs, AFs, and fibrotic tract fibroblast states. Nonetheless, our findings suggest that particular fibroblast states are strongly associated with type 2 and type 3/17 immune topography and, in vitro, similar AFs are sufficient to support both ILC2s (Dahlgren et al., 2019) and γδ T cells without exogenous cytokines or T cell receptor activation. Future work will be required to elucidate the likely cross-regulatory signals between type 2 lymphocytes, type 3/17 lymphocytes, stromal cell subsets, and other potential niche occupants such as dendritic cells, macrophage subsets, and lymphatics. Further exploration of the healthy and fibrotic hepatic immune-stromal interactions may delineate topographically restricted mechanisms of crosstalk and lead to disease-relevant therapeutic strategies.

### Limitations of the study

(1) IL-5 deleter mice lack type 2 lymphocytes, predominantly ILC2s, in all tissues from early life; therefore, this genetic model does not reveal a specific contribution of the fibroblast-associated liver ILC2s. However, we were able to phenocopy our findings in an inducible genetic model, suggesting any impact is not secondary to developmental alterations. (2) IL-5 deficiency in IL-5 deleter mice leads to reduction of eosinophil levels across tissues, which could impact the accumulation of regenerative alternatively activated macrophages (Molofsky et al., 2013). Although we used IL-5-deficient homozygote mice as control groups for our studies, ruling out an impact of IL-5, our results may minimize additional impacts of eosinophils on hepatic fibrosis. (3) Our work does not dissect how γδ T cells, or other similar IL-17A^+^ type 3/17 lymphocytes, contribute mechanistically to centrilobular liver fibrosis or portal fibrosis. Although we found that genetic ablation of IL-17A-producing cells reduced the degree of CCl_4_-induced liver fibrosis, in line with previous studies (Fabre et al., 2014; Fabre et al., 2018; Rau et al., 2016), we did not determine relevant cytokines (e.g. IL-17A, IL-22) or requirements for specific type 3/17 lymphocytes (e.g. γδ T cells, CD4^+^ Th17 cells, subsets of MAIT or NKT cells), in part due to considerable redundancy in the type 3/17 lymphocyte pool.

## ACKNOWLEDGEMENTS

We thank the UCSF Parnassus Flow Cytometry Core (RRID:SCR_018206) for its assistance with flow cytometry analysis and cell sorting, which is supported in part by NIH P30 DK063720; the UCSF Biological Imaging and Development Core (BIDC) and members Austin Edwards and Kyle Marchuk (confocal microscopy and analysis). JSK is supported by the German Research Foundation (DFG, Deutsche Forschungsgemeinschaft), FoeFoLe, LMU Munich, and the German Society of Internal Medicine (DGIM, Deutsche Gesellschaft für Innere Medizin, Clinician Scientist Program). ABM is supported by the NLHBI (NIH R01HL142701 and R56HL142701), NIDDK (K08DK101604), Larry L. Hillblom Foundation Grant, Nina Ireland Program for Lung Health, Sandler Asthma Basic Research Center (SABRE), and the UCSF Liver Center, RAP and PBBR grants.

## AUTHOR CONTRIBUTIONS

Conceptualization, JSK, KCM and ABM; Methodology JSK, KMC and ABM; Investigation, JSK, KMC, JCW, MWD, JN, MK, NMM, COL, ALG, PRM, MTT, AAC, SC, CEO, MK, ANM, TP, RL; Data Curation, JSK and KMC; Writing-Original Draft, JSK and ABM; Writing-Editing and Revision, JSK, KCM, ABM; Supervision, ABM

## COMPETING INTERESTS

The authors declare no competing interests.

## STAR METHODS

### CONTACT FOR REAGENT AND RESOURCE SHARING

Further information and requests for resources and reagents should be directed to and will be fulfilled by the Lead Contact, Ari B. Molofsky.

### EXPERIMENTAL MODEL AND SUBJECT DETAILS

#### Study design

The objective of this study was to investigate the crosstalk of ILC2s with stromal cells during the development of hepatic fibrosis. We used flow cytometry, thick section confocal imaging and different experimental models (FPC diet, bile duct ligation and CCl_4_) of liver fibrosis to localize and define the role of ILC2s as critical mediators of inflammation. Sample sizes were determined by power analyses from pilot studies or previously published data. Mice of both sexes were used and randomly assigned to experimental groups. Data were pooled from multiple experiments unless otherwise specified. For imaging experiments, at least three mice were analyzed from at least two independent experiments, with two or more fields analyzed per mouse. No data were excluded.

#### Mice

Mice were bred and maintained in specific-pathogen-free conditions at the animal facilities of University of California San Francisco (UCSF) and were used in accordance with the guidelines established by the Institutional Animal Care and Use Committee and Laboratory Animal Resource Center. All mice were a minimum of six weeks of age unless otherwise noted. Experiments were performed with mixed sex mice backcrossed on C57BL/6 for at least 10 generations. Red5 (Il5-tdtomato-cre) cytokine reporter mice were used for tracking IL-5-producing type 2 lymphocytes (Jackson 030926) (Nussbaum et al., 2013). Imaging was performed in Red5 mice crossed to R26-CAG-RFP mice (Ai14) containing a flox-stop-flox sequence upstream of a CAG-RFP-WPRE-cassette in the constitutively expressed ROSA26 (R26) locus (Jackson 007914) or in Red5 mice crossed to R26-CAG-YFP mice (Ai3) (Madisen et al., 2010), serving as IL-5 lineage-reporters (Dahlgren et al., 2019). To delete ILC2s, Red5 (Il5-tdtomato-Cre) mice were intercrossed with R26-DTA, an approach that specifically deletes ∼70-80% of ILC2s, as described previously (Nussbaum et al., 2013). To conditionally delete ILC2s, Red5 (Il5-tdtomato-Cre) mice were intercrossed with R26-DTR mice, generously provided by Anna V. Molofsky and previously described (Buch et al., 2005). To concurrently delete IL-17A^+^ type 3/17 lymphocytes and IL-5^+^ type 2 lymphocytes, we used IL-17ACre mice (Hirota et al., 2011) crossed with IL-5 deleter mice (RRDD; *Il5*^Red5/Red5^*Gt (Rosa)26*^DTA/DTA^). Additional mice utilized include genetically targeted IL-33mcherry (Vainchtein et al., 2018b), PDGFRα-H2B-eGFP nuclear-localized GFP (Jackson 007669), RORγt-GFP(Lochner et al., 2008), *Arg1*^Yarg^ (Yarg; B6.129S4-*Arg1^tm1Lky^*/J; 015857; (Reese et al., 2007)), *Il13*^Smart^ (Smart13; B6.129S4[C]-*Il13^tm2.1Lky^*/J; 031367; (Liang et al., 2012), and *Arg1*^RFP-CreERT2^ (Schneider et al., 2019) mice. PDGFRα-H2B-eGFP; FoxP3DTR/αSMA-RFP mice were generated by crossing PDGFRα-H2B-eGFP to FoxP3DTR/αSMA-RFP mice which were kindly provided by Michael Rosenblum and were previously described (Kalekar et al., 2019). Tbet (Tbx21)-zsGreen transgenic mice were a kind gift of Jinfang Zhu, Lab of Immune System Biology, NIH, and were previously described (Zhu et al., 2012). IL4ra^-/-^ mice were generated as previously described (Mohrs et al., 1999) and were generously provided by F. Brombacher. *TCRd*^-/-^ mice (Jackson 002120) and C57BL/6J (Jackson 000664) were purchased from The Jackson Laboratory.

## METHOD DETAILS

### FPC diet

Mice were fed a fructose-palmitate-cholesterol (FPC) diet (Teklad, TD.140154) with 1.25% added cholesterol and with palmitic acid, anhydrous milk fat, and Primex as the sources of fat and with a ∼60% decrease in vitamin E and a ∼35% decrease in choline compared with typical mouse diets (Wang et al., 2016) for 16 weeks. Control groups were fed a chow diet (Picolab rodent diet 20, #5053).

### CCl4 treatment

Mice were treated with CCl_4_ (Sigma-Aldrich, Cat# 289116), resuspended in corn oil, as 0.5ml/kg or vehicle (corn oil) with three i.p. injections per week for the indicated duration.

### Bile duct ligation

All procedures were carried out under clean but non-sterile conditions following standard operating aseptic protocols. Mice were anesthetized with inhalation of isoflurane after i.p. injection of ketamine/xylazine (80-100 mg/kg Ketamine + 5-10 mg/kg Xylazine) and buprenorphine analgesia (0.05-0.1 mg/kg) administered via subcutaneous injection. The abdomen was opened by a midline laparotomy approximately 2 cm in length with a surgical scissor and the peritoneum cut open along the linea alba. The peritoneal cavity was enlarged by inserting a Colibri retractor and the hilum was revealed using a moisturized (0.9% NaCl solution) cotton swab. The bile duct was exposed and separated from the portal vein and hepatic artery using a micro-serrations forceps. The 7-0 suture was placed around the bile duct and secured with two surgical knots. A second cranial ligation was added in the same manner. Abdominal layers (peritoneum and cutis plus facia) were closed with running 6-0 sutures with absorbable suture material. Mice were allowed to recover in a cage warmed up by an infrared lamp until they were fully awake and active. Mice were weighed daily and monitored for humane endpoint. Mice were either euthanized after 7 or 14 days.

### Administration of diphteria toxin

Diphteria toxin (DT) (Sigma-Aldrich) was administered i.p. at a dose of 500ng in sterile PBS initially 2 to 1 day before CCl_4_ injections and then every other day throughout the experiment.

### Cytokines, neutralizing antibodies, and chemicals

For cytokine injections, Interleukin-33 (Biolegend) was given intraperitoneally as 500ng in 0.2ml PBS every day for three doses. For the therapeutic intervention, mice were placed on CCl_4_ (0.5ml/kg, i.p., three times per week) for 4 weeks and treated with 250 µg of mouse anti–IL-17a (clone 17F3, BioXcell) antibody intraperitoneally in a volume of 200 µL (diluted in PBS) at the beginning of CCl_4_ application and throughout the experiment or with the RORγt antagonist GSK805 (Sigma-Aldrich, Cat# 5313690001) from weeks 2 to 4 daily with 10mg/kg i.p. in corn oil. Vehicle (corn oil) was used as a control. For bleomycin injury, mice were given pharmaceutical grade bleomycin (Hospira) dissolved in PBS via intranasal application once a week for four weeks, as previously described (Cassandras et al., 2020). Mice were given a dose of 0.75U/kg per dose.

### Hydrodynamic delivery of IL-33

Mice were injected intravenously with 1µg or 10µg of a mammalian expression plasmid coding for murine IL-33 in 1ml of warm saline (Suda and Liu, 2007).

### Flow cytometry

Single cell suspensions were prepared from tissues including blood, liver, lung and epigonadal white adipose tissue (eWAT). After euthanizing the mice with CO2, peripheral blood was collected through cardiac punction, and mice were perfused with 10ml PBS via the left ventricle. Livers were weighed, cut in small pieces with an automated tissue dissociator (Gentle Macs; Milteny Biotec), and then digested in 7 ml Hank’s balanced salt solution (HBSS) with 0.5% bovine serum albumin (Sigma-Aldrich, Cat# A2153), 2% FCS, 40µl Liberase Tm (0.1 wU/ml, Roche, Cat# 5401127001), 40µl DNAse 1 (10mg/ml, Roche, Cat# 10104159001) for 30 min at 37°C with gentle agitation. Samples were subsequently processed on the GentleMacs using the “lung2” program, passed through 70µm filters, and centrifuged at 30 x g, 3min, 4°C to remove hepatocytes. Supernatant was centrifuged and leukocytes were further separated using 40% Percoll density gradient (GE Healthcare Cat# 17-0891-01) and centrifugation (1400 x g, 20 min, room temperature, no brake), followed by red blood cell lysis (PharmLyse; BS Biosciences) before final suspension in FACS buffer (PBS, 3% FCS, 0.05% NaN3). Whole lung single cell suspensions were prepared as previously described (Dahlgren et al., 2019) by harvesting lung lobes into 5 ml HBSS with 40µl Liberase Tm (0.1 wU/ml, Roche, Cat# 5401127001) and 20µl DNAse 1 (10mg/ml, Roche, Cat# 10104159001), followed by automated tissue dissociation (GentleMacs; Miltenyi Biotec) and tissue digestion for 30 min at 37°C on a shaker. Digested samples were processed on the GentleMacs using the “lung2” program, passed through 70µm filters, and washed, followed by red blood cell lysis and final suspension in FACS buffer. The eWAT was harvested, cut into small pieces with a scissor, digested in 10ml of low-glucose DMEM containing 0.2mg/ml Liberase Tm, 25µg/ml DNase, 0.2M HEPES and 10mg/ml BSA for 45 min at 37°C with gentle agitation, passed through 100µm filters and centrifuged at 1000 x g for 10 min. Red blood cells were lysed with PharmLyse and the remaining cell pellets were resuspended in FACS buffer. Blood samples were centrifuged for 5 min at 1500 x g and resuspended in PharmLyse for ∼10 min at RT, followed by centrifugation and final suspension in FACS buffer. Cells were counted using a NucleoCounter (Chemometic). All samples were stained in 96-well V-bottom plates. Single cell samples were first incubated with antibodies to surface antigens for 30 min at 4°C in 50µl staining volume. For intracellular staining, cells were fixed and permeabilized using FoxP3/Transcription Factor Staining Buffer Set (eBioscience, Cat# 00-5523-00). For cytokine staining, ∼2-5 x 10^6^ cells were plated in 96 well U bottom plates and stimulated ex vivo with 30ng/ml PMA (Sigma-Aldrich, Cat# 79346-1mG) and 500ng/ml Ionomycin (Sigma-Aldrich, Cat# IO634-5MG) in culture medium (RPMI 1640 + 10% FBS, 10% penicillin/streptomycin, 50mM 2-ME (Sigma-Aldrich, Cat# 60-24-2), and 1000x BD Golgi Plug containing Brefeldin A (Sigma-Aldrich, Cat# 555029) for 3 hours at 37°C. For the detection of vascular-associated lymphocytes, 3µg of anti-mouse fluorophore-conjugated CD45 antibody diluted in PBS were intravenously injected, 3-5 min before euthanization, as previously described (Anderson et al., 2014). Flow cytometry was performed on BD LSRFortessa X-20. Fluorochrome compensation was performed with single-stained UltraComp eBeads (Invitrogen, Cat# 01-2222-42). Samples were FSC-A/SSC-A gated to exclude debris, followed by FSC-H/FSC-A gating to select single cells and Zombie NIR fixable or DAPI to exclude dead cells. ILC2s were identified as lineage negative (CD11b^-^, CD3ε^-^, CD4^-^, CD8α^-^, CD19^-^, NK1.1^-^), CD45^+^, Thy1.2 (CD90.2)^+^, and Gata3^hi^, IL1RL1 (ST2)^+^, or CD25 (IL-2Rα)^+^, as indicated. Th2 cells were identified as CD45^+^, CD3ε^+^, CD4^+^, and FoxP3^-^. For some analyses, IL-5 lineage tracker mice (Il5-tdtomato-Cre; R26-RFP) were used to gate ILC2s as lineage negative, Thy1.2 (CD90.2^+^), RFP^+^ and Th2 cells as CD45^+^, CD3ε^+^, CD4^+^, RFP^+^. Data were analyzed using FlowJo software (TreeStar, USA) and compiled using Prism (GraphPad Software).

### Flow cytometry Antibodies

Monoclonal antibodies used for flow cytometry include anti-CD45 (30-F11, Biolegend), anti-CD90.2 (Thy1.2) (53-2.1, Biolegend), anti-CD3 (17A2, Biolegend), anti-CD4 (RM4-5, Biolegend), anti-CD8 (53-6.7 Biolegend), anti-CD11b (M1/70, Biolegend), anti-CD11c (N418, Biolegend), anti-NK1.1 (PK136, Biolegend), anti-CD19 (6D5, Biolegend), anti-TCRγ/δ (GL3, Biolegend), anti-T1/ST2 (DJ8, MD BioSciences), anti-KLRG1 (2F1, Biolegend), anti-IL13 (eBio13A, eBioscience), anti-MerTK (DS5MMER, eBiosciences), anti-64 (X54-5/7, eBiosciences), anti-Ly6C (HK1.4, Biolegend), anti-Ly6G (RB6-8C5), anti-SiglecF (E50-2440), anti-I-A/I-E (MHCII) (M5/114.15.2, Biolegend), anti-CD25 (PC61, Biolegend), anti-IFNγ (XMG1.2, Biolegend), anti-IL17A (TC11-18H10.1, Biolegend), and anti-RORγt (B2D, eBioscience). Anti-FoxP3 (FJK-16S, eBiosciences), anti-Ki-67 (16A8, Biolegend) and anti-GATA3 (TWAJ, eBiosciences) were utilized after first using a fixable live/dead stain (Invitrogen), then fixing and permeabilizing cells per manufacturer’s instructions. For non-hematopoietic cells, antibodies used include anti-CD31 (390, Biolegend), anti-EpCAM (CD326, G8.8, eBiosciences), anti-Gp38 (podoplanin) (8.1.1, Biolegend), anti-PDGFRα (CD140a, APA5, Biolegend), and anti-Ly6A/E (Sca1, D7, eBioscience).

### Imaging antibodies

Primary antibodies used for imaging include Living Colors anti-DsRed Rabbit Polyclonal Pan Antibody (1:500; TaKaRa), Chicken Polyclonal anti-GFP (1:300, Aves labs), Alexa Fluor 488 anti-aSMA monoclonal Antibody (IA4, 1:200; eBioscience), eFluor 660 anti-LYVE1 monoclonal Antibody (ALY7, 1:500, eBioscience), Goat Monoclonal anti-mouse Collagen Type 1 (1:200, Southern Biotech), Rat Monoclonal anti-mouse CD326 (G8.8, 1:200, BD Pharmingen), Rabbit Monoclonal anti-mouse CD34 (EP373Y, 1:200, Abcam), Mouse Monoclonal anti-mouse Glutamin Synthetase (GS-6, 1:250, Millipore), Rabbit Polyclonal anti-SDS (PA5-58704), eBiosciences), Goat Monoclonal anti-mouse Desmin (GWB-EV0472, 1:200, Genway Biotech), Goat Polyclonal anti-VEGFR3 (R&D Systems), Rabbit Monoclonal anti-mouse Vimentin (EPR3776, 1:200, Abcam), and Rabbit Polyclonal anti-Cytochrome P450 2E1 (1:250, Abcam). The following secondary antibodies were used at 1:300 dilution: Alexa Fluor 555 donkey anti-rabbit IgG (H+L) cross-adsorbed (ThermoFisher Scientific), Alexa Fluor 647 donkey anti-rat IgG (H+L) cross-adsorbed (Abcam), Alexa Fluor 647 donkey anti-rabbit IgG (H+L) cross-adsorbed (Life Technologies), Alexa Fluor 488 donkey anti-rat IgG (H+L) cross-adsorbed (Abcam), Alexa Fluor 488 donkey anti-chicken IgG (H+L) cross-adsorbed (Sigma-Aldrich), Alexa Fluor 647 donkey anti-goat IgG (H+L) cross-adsorbed (ThermoFisher Scientific), Alexa Fluor 555 donkey anti-mouse IgG (H+L) cross-adsorbed (ThermoFisher Scientific).

### 3D Tissue Preparation and Imaging

Animals were sacrificed with CO2, followed by transcardial perfusion with 20ml PBS and 4% paraformaldehyde (PFA) (Thermo Scientific, Cat# 28906). Tissues (liver, adipose tissue, gall bladder, and lung) were harvested and fixed in fresh 4% PFA overnight at 4°C. Fixed tissues were cut into ∼200µm thick sections using a vibratome (Leica VT 1000S or Precisionart Compresstome VF-310-0Z). Tissue slices were permeabilized (PBS/0.2%TritonX-100/0.3M glycine), then blocked in PBS, 0.2% Triton-X-100, 10% FBS, 1% BSA, 5% serum (appropriate species serum for secondary antibodies) at 4°C overnight. Samples were then incubated with primary antibodies in PBS, 0.2% TritonX-100, 3% serum at 4°C until the next day. Next, samples were washed in PBS, 0.2% TritonX-100 for 30 min, 3-4 times, then incubated with secondary antibodies diluted in PBS, 0.2% Triton-X-100, 10% FBS, 1% BSA, 5% serum at 4°C overnight. Samples were then washed in PBS, 0.2% TritonX-100 for 30 min, 3-4 times, dehydrated in an ascending ethanol series (20, 30, 50, 70, 95, 100%), 10 min each step, and cleared in methyl salicylate. For identification of ILC2s, liver, and lung tissues were stained with anti-tdtomato (1:200). For localization of ILC2s to anatomical structures, samples were stained with anti-aSMA (1:200), anti-Lyve 1(1:500), anti-type 1 collagen (1:200), anti-CD326 (1:200), anti-CD34 (1:200), anti-Desmin (1:200), and anti-Vimentin (EPR3776, 1:200, Abcam). All samples were scanned using a Nikon A1R laser scanning confocal including 405, 488, 561, and 650 laser lines for excitation and imaged with 16X/0.8, NA Plan Apo long working distance water immersion objective. Z-steps were acquired every 2µm with 120-200µm of each slice imaged. Images were acquired in galvo mode with pixel sizes of 300nm and 2X frame averaging.

### Histopathology

Mice were euthanized by CO2 and then perfused with 20ml 1X PBS, followed by 20ml 4% PFA/PBS. Livers and lungs were harvested and post fixed overnight in 4% PFA/PBS on a shaker. Formalin-fixed samples were embedded in paraffin, cut in 5µm sections, and stained with hematoxylin & eosin (H&E), Picrosirius red, and Trichrome by Peninsula Histopathology Laboratory. The Picrosirius red positive area was morphometrically quantified with FIJI image analysis software (ImageJ). Images were converted to grayscale and the fibrotic area (Sirius red) was calculated using total liver section area and Sirius red positive area calculated by the thresholding method. H&E, Sirius Red and Trichrome stains were reviewed and evaluated by a blinded UCSF liver pathologist (A.N.M.) for steatosis and lobular inflammation using a histological scoring system for nonalcoholic fatty liver disease. Livers from FPC-fed mice were evaluated and scored according to methods by Kleiner and Brunt (Kleiner and Brunt, 2012).

### Hydroxyproline Assay

For fibrosis quantification, hydroxyproline assay was performed as described previously (Fabre et al., 2018). Briefly, 80-200 mg liver tissue were hydrolyzed in 1.5 ml of 6 N HCl at 110**°**C overnight. 10 µl of standards or the hydrolyzed samples were pipetted as triplets into a 96-well optically clear plate with 30 µl of citric acid buffer. 100 µl of Chloramine T solution was added and allowed to oxidize for 20 min at room temperature. 100 µl of Ehrlich’s Reagent was mixed with the oxidized samples or standards and incubated at 65**°**C for 20 min. Absorbance was read at 550nm in spectrophotometer and samples were quantified by comparison to the standard curve.

### Serum analysis

Blood was harvested after cardiac puncture and processed for collection of serum, snap frozen, and stored at-80°C until further testing. Bilirubin was measured using the Bilirubin Assay Kit following the manufacturer’s instructions (Sigma-Aldrich, Cat# MAK126-1KT). Alanine aminotransferase (ALT) was assayed on Roche Diagnostic’s Cobas Mira.

### Total tissue RNA extraction and qPCR

RNA from mouse liver tissue was obtained by homogenizing in trizol (ThermoFisher, Cat#15596018) and extracting RNA using the E.Z.N.A. Total RNA Kit (Omega Bio-Tek, Cat#R6834-01) following manufacturer’s instructions. RNA was reverse transcribed using SuperScript III cDNA synthesis kit (ThermoFisher) and cDNA was used as template for quantitative PCR (aPCR) using Power SYBR Green PCR master mix (ThermoFisher). Transcripts were normalized to *Gapdh* expression and relative expression shown as 2-ΔΔCt compared to the average ΔCt of experiment-matched controls. The following primers were used:

*Desmin: 5’-GTTTCAGACTTGACTCAGGCA-3’, 5’-TCTCGCAGGTGTAGGACTGG-3’*

*Acta2: 5’-CCCAAAGCTAACCGGGAGAAG-3’, 5’-GACAGCACCGCCTGGATA-3’*

*Snail: 5’-CACACGCTGCCTTGTGTCT-3’, 5’-GGTCAGCAAAAGCACGGT T-3’*

*Col1a1: 5’-CCAAGAAGACATCCCTGAAGTCA-3’, 5’-TGCACGTCATCGCACACA-3’*

*Epcam: 5’-GCGGCTCAGAGAGACTCT-3’, 5’-CCAAGCATTTAGACGCCAGTTT-3’*

*Tgfb1: 5′-TGGAGCAACATGTGGAACTC-3′, 5′-GTCAGCAGCCGGTTACCA-3′*

### Mesenchymal-ILC2-γδ T cell-co-cultures

Lungs from wildtype C57Bl/6 mice were harvested, manually dissociated with a razorblade, and subsequently digested with 7.5U/ml Dispase II, 112.5U/ml Collagenase I and 40μg/ml DNase I in PBS for 30 min at 37°C with gentle agitation. Samples were subsequently passed through 70μm filters, washed, subjected to red blood cell lysis (PharmLyse; BD Biosciences), and enriched for stromal cells through magnetic bead separation of CD45^+^ and CD31^+^ cells before final suspension in FACS buffer (PBS, 3% FCS, 50U/mL penicillin, and 50μg/mL streptomycin). Sorted adventitial fibroblasts (CD45^-^, EpCAM^-^, PDGFRα^+^ Sca1^+^) were seeded in flat bottomed 96-well plates in 200μl DMEM (supplemented with 10% FBS, 1X Glutamax, 50U/mL penicillin and 50 μg/mL streptomycin) at a density of 10,000 cells per well and allowed to form monolayers over 6 days. Lungs from IL-33-injected, Il5tdtomato-^Cre/+;^ Rosa26^RFP/+^ mice were harvested for ILC2 purification and lungs from wild-type C57Bl/6 mice were harvested for γδ T cell purification, subsequently manually dissociated (Miltenyi GentelMacs) and digested with 0.2 wU/mL Liberase TM and 40 µg/mL DNase I in PBS for 30 min at 37°C with gentle agitation. Samples were manually dissociated (Miltenyi GentelMacs) a second time following digestion, passed through 70μm filters, washed, and subjected to red blood cell lysis (PharmLyse; BD Biosciences) before final suspension in FACS buffer (PBS, 3% FCS, 50U/mL penicillin, and 50μg/mL streptomycin). Sorted ILC2s (lin^−^, CD45^+^, CD3^-^, CD4^-^, RFP^+^) and/or γδ T cells (lin^-^, CD45^+^, CD3^+^, TCRβ^-^, TCRγδ^+^), were seeded onto the stromal monolayers or cultured with only DMEM in a total volume of 200μl, at a density of 7,000 ILC2s and/or γδ T cells per well. After 6 days of culture, supernatants were collected, and cells were liberated, stained for viability dye, CD45, CD3, TCRγδ, Sca-1 and PDGFRα, analyzed and enumerated using CountBright Absolute counting beads (Life Technologies) by flow cytometry.

### In vitro cultures and 3’ Tag RNAseq

12 000 sorted (viable, CD45^-^, CD31^-^, EpCAM^-^, Pdgfra^+^, Sca1^+^) primary lung fibroblasts were seeded per well in flat bottomed 96-well plates (DMEM +10% FBS, +1% Penicillin/Streptomycin) and grown to confluence over 6 days. Cells were washed with pre-warmed serum-free media and TGF-β (Biolegend) was added in a final concentration of 1 ng/mL. Fibroblasts were harvested after 48hrs of stimulation and total RNA was isolated using QIAGEN RNeasy Plus Micro Kit according to the manufacturer’s instructions. RNA quality and quantitation were assessed on a Bioanalyzer (Agilent). 3’-Tag RNAseq library preparation and sequencing was done by DNA Technologies and Expression Analysis Core at the UC Davis Genome Center as previously described (Cautivo et al., 2022). Normalized reads were visualized using pheatmap (v1.0.12).

## QUANTIFICATION AND STATISTICAL ANALYSIS

### Image analysis and quantification

Imaris Bitplane 9.7.2 software package (Andor Technology PLC, Belfast, N. Ireland) was used for all 3D image analysis and Z-stacks images were rendered in 3D dimensions. To parse PDGFRα subsets, a colocalization channel was made between PDGFRα and anti-IL-33. PDGFRαGFP^+^ stromal cells, IL-5^+^, IL-17A and RORγt^+^ lymphocytes were annotated using the Imaris spot’s function based on the fluorescent reporter signal. To localize IL-5^+^, IL-17A^+^ and RORγt^+^ lymphocytes with different stromal cell types and bile ducts, 3D reconstructions of structures labeled for Collagen Type 1 and CD326 were generated, followed by calculation of three-dimensional distances between lymphocytes and Collagen Type 1^+^ surfaces using the Imaris Distance Transform Matlab XTension and volumetric decile calculations were performed using a Matlab-based Imaris XTension.

### Statistical analysis

All data are expressed as means ± standard error of the mean (SEM) unless otherwise noted. Comparisons between two groups were analyzed by using unpaired two-tailed Student’s t-tests, and multiple comparisons were analyzed by one-way analysis of variance (ANOVA) with Tukey’s multiple comparisons test (Prism, GraphPad Software, La Jolla, CA), with * = p<0.05, ** = p<0.01, *** =p<0.001, **** =p<0.0001. Each symbol reflects individual biological mouse replicates for flow analysis, or individual tissue slices analyzed for confocal imaging

## Supplemental information

### Supplementary Figure 1

**Figure S1:**
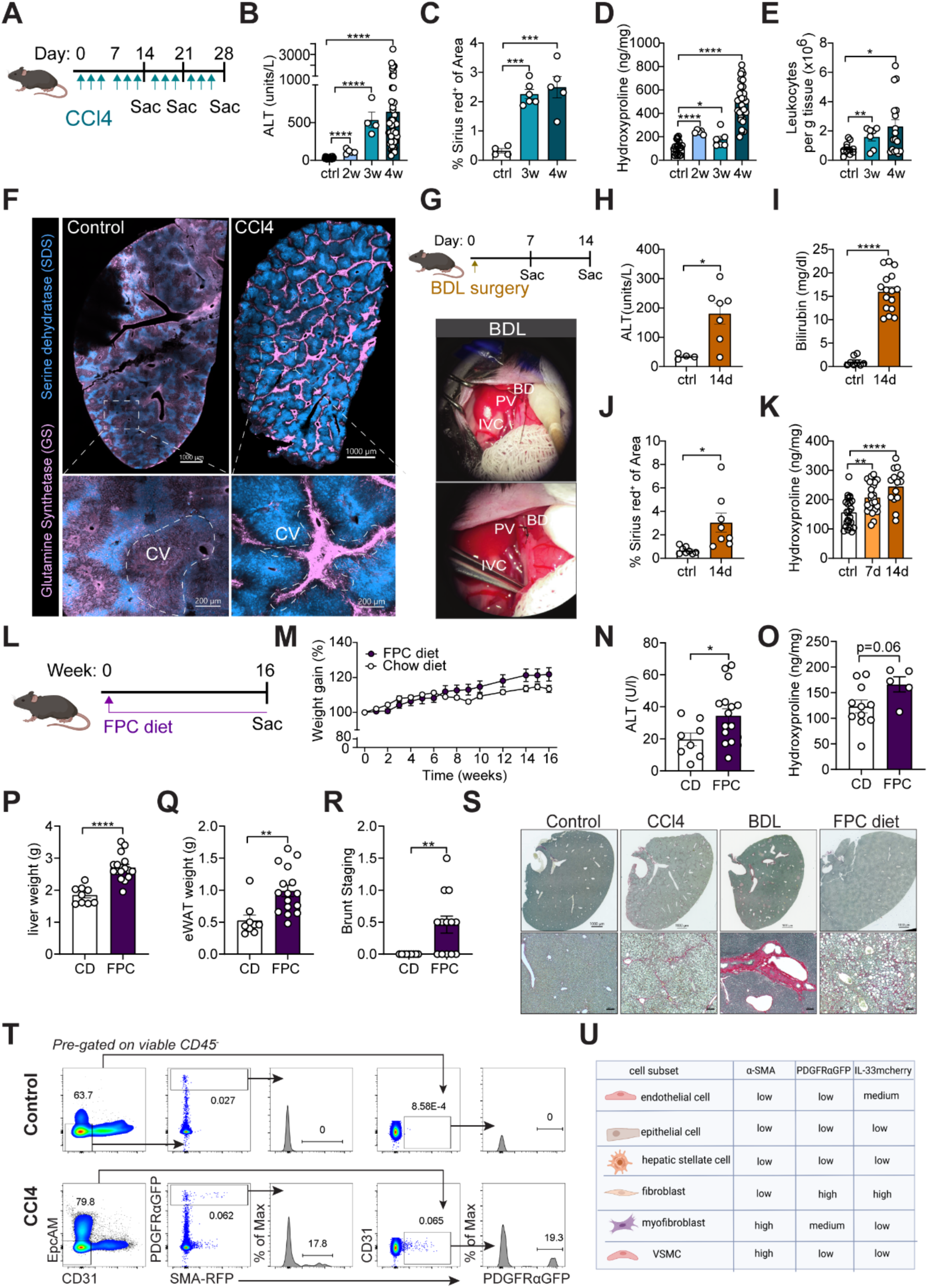
Models of liver fibrosis and impact on stromal cell topography (related to **Figure 1).** **(A)** Schematic showing CCl_4_ administration schedule in IL-33^mcherry/+^ reporter mice injected intraperitoneally (i.p.) with 0.5µl CCl_4_/g BW three times per week for 2, 3, or 4 weeks (2w, 3w, 4w), relevant to B-E. **(B-E)** Quantification of ALT **(B)**, Sirius red positive staining **(C)**, hepatic Hydroxyproline levels **(D),** total leukocytes **(E)** in control mice and CCl_4_-treated mice at the indicated timepoints. Pooled data from 8-10 weeks old age-and sex-matched mice for 4-week timepoint. **(F)** Representative thick section liver image from control and 4-week CCl_4_ treated IL-33^mcherry/+^ reporter mice with staining for glutamin synthetase (GS), a marker for pericentral hepatocytes, and sodium dodecyl sulphate (SDS) surrounding portal veins. Higher magnification image highlighting severe structural changes in liver morphology with CCl_4_-induced fibrosis. **(G)** Schematic of bile duct ligation (BDL) surgery in PDGFRαGFP; IL33^mcherry/+^ reporter mice for time course analysis, relevant to H-K. Bile duct (BDL), Portal vein (PV), Inferior vena cava (IVC). **(H-K)** Quantification of ALT **(H)**, Bilirubin **(I)**, Sirius red positive staining **(J)**, and hepatic Hydroxyproline levels **(K)** in control mice and bile duct ligated mice at the indicated timepoints. Pooled data from 3 independent experiments from 8-10-weeks-old age-and sex-matched mice. **(L)** Schematic of fructose-palmitate-cholesterol (FPC) diet in IL-33^mcherry/+^ mice, relevant to M-S. **(M-R)** Body weight gain **(M)**, quantification of ALT **(N)** and hepatic Hydroxyproline levels **(O)**, liver weight **(P)**, epigonadal white adipose tissue (eWAT) weight **(Q)** and Brunt staging **(R)** in 16w chow diet-fed and 16w FPC diet-fed mice. Data pooled from 3 independent experiments from 6-8-weeks-old IL-33^mcherry/+^ reporter age-and sex-matched mice. **(S)** Representative Sirius Red staining of 5µm paraffin liver sections from IL-33 ^mcherry/+^ reporter mice at steady state, at day 28 of CCl_4_ treatment, at day 14 post BDL surgery, and at week 16 of FPC diet feeding. Higher magnification images show detailed morphology of collagen deposition. Scale Bars: 100µm (10X). Images are representative of 3 repeat experiments with n=3-10 mice per group. **(T)** Representative flow cytometry plots from liver of PDGFRαGFPSMARFP reporter mice treated with vehicle or CCl_4_ for 4 weeks and representative histograms of αSMA expression on PDGFRαGFP^+^ fibroblasts. Data are representative of 2 independent experiments. **(U)** Schematic showing expression of α-SMA, PDGFRαGFP and IL-33mcherry in different stromal cell subsets. Bar graphs indicate mean (±SE). Unpaired t-test, *p ≤ 0.05, **p ≤ 0.01, ***p ≤ 0.001, ****p ≤ 0.0001.

### Supplementary Figure 2

**Figure S2:**
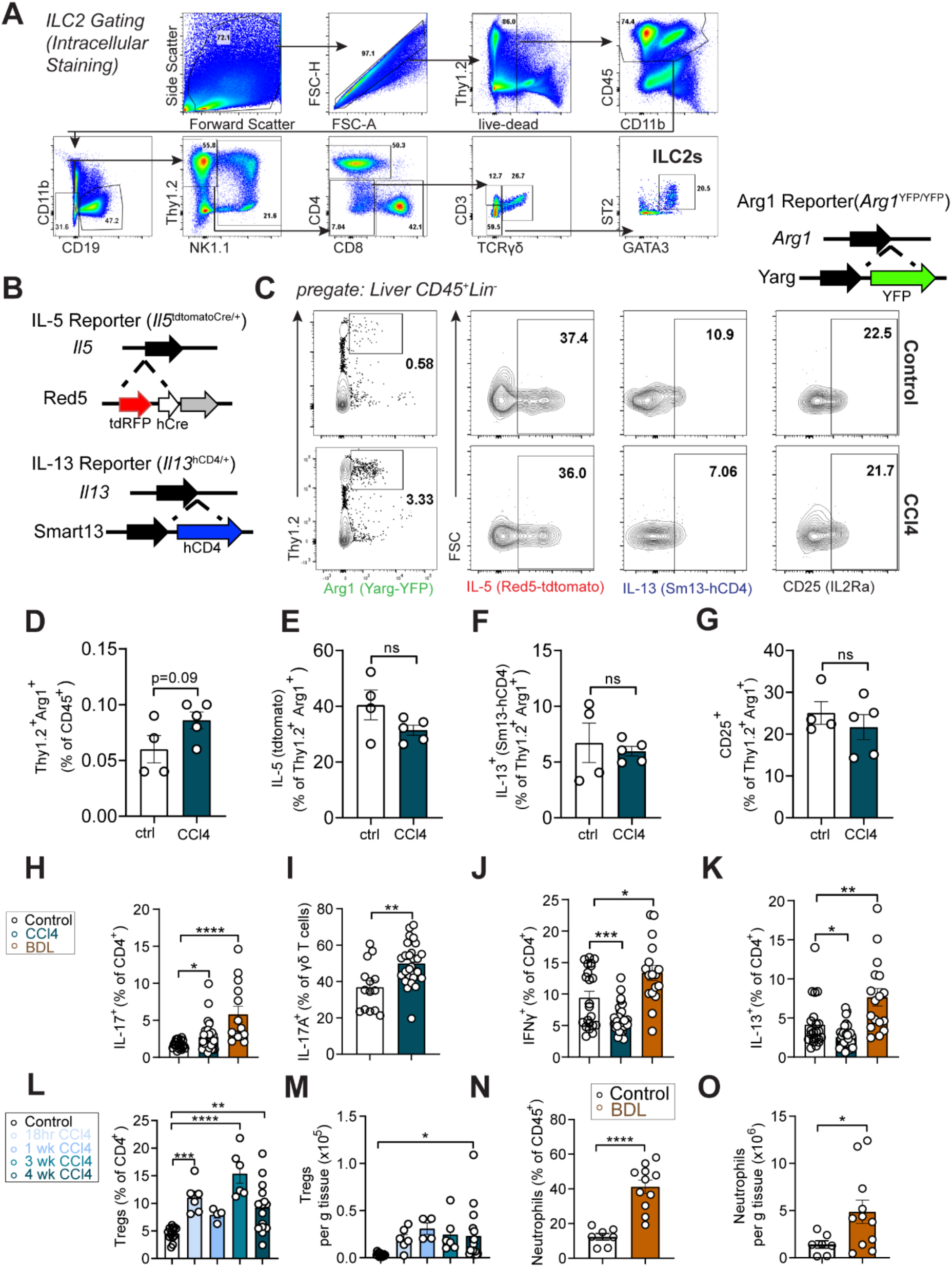
Type 2 and type 3/17 lymphocytes expand in liver damage and fibrosis (related to **Figure 2).** (A) Representative flow gating scheme for liver ILC2s (viable CD45^+^Lin^-^GATA3^+^ST2^+^) and T cells. (B) Schematic of the IL-5^+^ lymphocyte lineage tracker mice (IL-5tdtomato-Cre; Rosa26RFP), *Il13*^Smart^ (Smart13; B6.129S4[C]-*Il13^tm2.1Lky^*/J; 031367), and *Arg1*^RFP-^ _CreERT2 mice._ (C-G) Representative flow cytometry plots (C) and quantification of ILC2 (Lin^-^ Thy1.2^+^Arg^+^) (D) expression of IL-5RFP (E), IL-13 (F), and CD25 (G) in livers of control mice and 4-week CCl_4_-treated mice on Arg1 (Yarg); R5 (IL-5); S13 (IL-13) combined triple-reporter (YRS) background. (H-K) Flow cytometry quantification showing percentages of IL-17^+^CD4^+^ T cells (H), IL-17^+^ γδ T cells (I), IFNγ^+^CD4^+^ T cells (J), and IL-13^+^CD4^+^ T cells (K) in livers from C57BL/6 wildtype mice treated with vehicle or CCl_4_ for 4 weeks. Pooled data from 4 independent experiments from 8-10 weeks-old age-and sex-matched mice. (L and M) Flow cytometry quantification showing percentage (L) and total numbers (M) of Tregs in livers from IL-33^mcherry/+^ reporter mice treated with vehicle or CCl_4_ for the indicated duration. Pooled data from 3 independent experiments from 8-10 weeks-old age-and sex-matched mice for 4-week timepoint. (N and O) Flow cytometry quantification of percentage (N) and total numbers (O) of neutrophils in livers from IL-33^mcherry/+^ reporter mice 14 days post-BDL surgery. Pooled data from 3 independent experiments from 8-10-weeks-old age-and sex-matched mice. Bar graphs indicate mean (±SE), unpaired t-test, *p ≤ 0.05, **p ≤ 0.01, ***p ≤ 0.001, ****p ≤ 0.0001.

### Supplementary Figure 3

**Figure S3:**
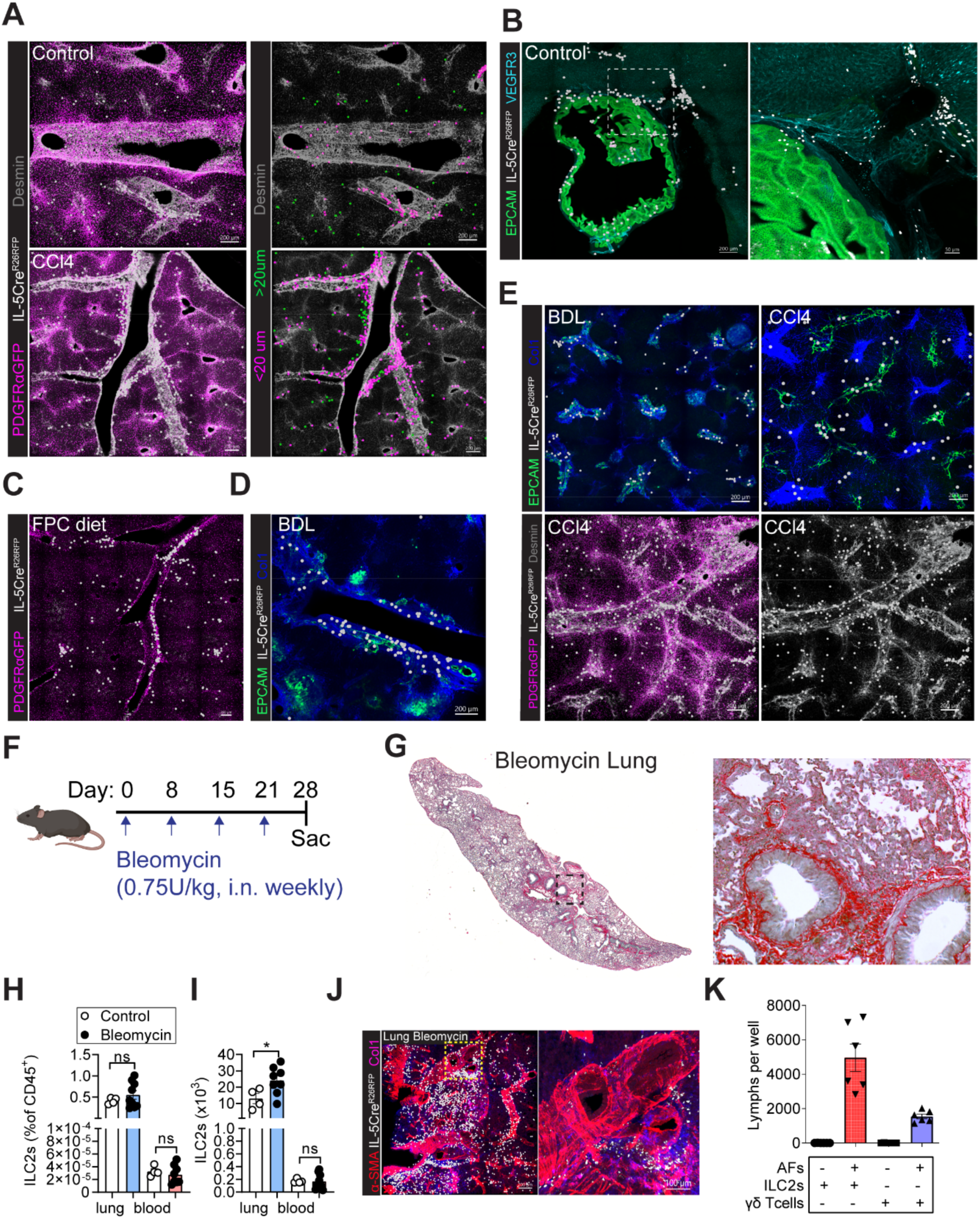
Type 2 lymphocytes localize to periportal regions at steady state and expand into fibrotic tracts with liver fibrosis (related to **Figure 3).** (A) Representative confocal imaging of liver from IL-5^+^ lineage tracker mice (IL-5tdtomato-Cre; Rosa26RFP) treated with vehicle or CCl_4_ for 4 weeks with surface rendering of IL-5^+^ lymphocytes and indicated staining. Images are representative of three or more mice. (B) Representative confocal imaging of gallbladder and liver from naïve IL-5^+^ lymphocyte lineage tracker mice with staining for EpCAM (epithelial duct cells) and VEGFR3 (lymphatics). Higher magnification highlighting accumulation of IL-5^+^ lymphocytes around the bile ducts. Image is representative of n=3 mice. (C-E) Representative confocal thick section imaging of liver from IL-5^+^ lineage tracker mice fed with FPC diet for 16 weeks (C), 14 days post BDL surgery (D, E), treated with CCl_4_ for 4 weeks (E), and surface rendering of IL-5^+^ lymphocytes and indicated staining. Images are representative of three or more mice. (F) Schematic showing Bleomycin administration schedule in IL-5 reporter mice subjected to weekly intranasal applications of Bleomycin (0.75U/kg), relevant to G-J. (G) Representative Sirius red staining from IL-5tdtomato-Cre mice treated with Bleomycin for 4 weeks. Images are representative of one experiment with n=11 8-10 weeks-old age-and sex-matched mice. (H and I) Flow cytometry quantitation of percent (H) and total numbers (I) of ILC2s in lung and blood from PDGFRαGFP; IL-33^mcherry/+^ mice treated weekly with Bleomycin or PBS (intranasal application) for 4 weeks. (J) Representative confocal imaging of cleared thick lung sections with surface analysis for IL-5RFP^+^ lymphocytes from Il5-tdtomato-Cre mice treated weekly with Bleomycin (intranasal application) for 4 weeks. Images are representative of one experiment with n=11 Il5-tdtomato-Cre mice. (K) PDGFRα^+^Sca1^+^ lung adventitial fibroblasts (AFs) were cultured with lung ILC2s and γδ T cells for 7 days; ILC2s and γδ T cells were counted. Pooled data from 2 independent experiments. Bar graphs indicate mean (±SE). Unpaired t-test for D, ns= not significant, *p ≤ 0.05.

### Supplementary Figure 4

**Figure S4:**
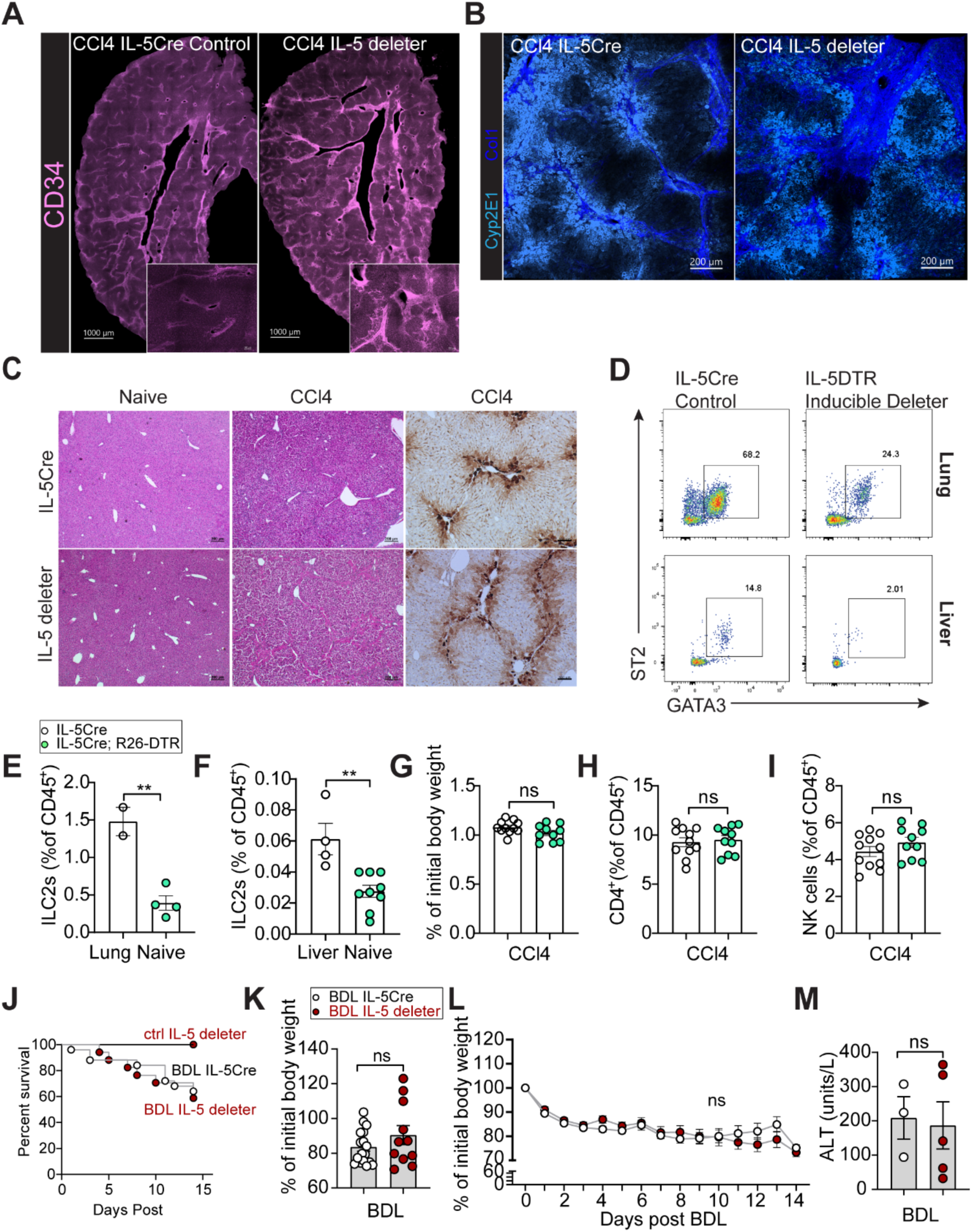
Loss of IL-5^+^ lymphocytes worsened hepatic fibrosis (related to **Figure 5).** (A) Representative confocal liver tissue sections from control or 4-week CCl_4_-treated Il5-tdtomato-Cre mice and IL-5 deleter (RRDD; *Il5*^Red5/Red5^*Gt (Rosa)26*^DTA/DTA^) mice with staining for CD34. Images shown are representative of 3-4 mice/group. (B) Representative confocal thick section imaging of liver from IL-5tdtomato-Cre and IL-5 deleter mice with staining for Collagen I and Cyp2E1 (marker of pericentral hepatocytes). Images are representative of three or more mice. (C) Representative Sirius red staining and glutamin synthetase (GS) immunohistochemistry from IL-5tdtomato-Cre and IL-5 deleter (RRDD; *Il5*^Red5/Red5^*Gt (Rosa)26*^DTA/DTA^) mice treated with vehicle or CCl_4_ for 4 weeks. Images are representative of 3 independent experiments with 8-10 weeks-old age-and sex-matched mice. (D-F) Representative flow cytometry plots of ILC2s (Lin^-^Thy1.2^+^GATA3^+^ST2^+^) (D) and flow quantitation of ILC2 percentages in lung (E) or liver (F) from either IL-5tdtomato-Cre or IL-5^DTR^ treated with CCl_4_ for 4 weeks. (G-I) Percent of initial body weight before CCl_4_ treatment (G) and percent CD4^+^ T cells (H) and percent NK cells (I) in livers of 4-week CCl_4_-treated IL-5tdtomato-Cre and IL-5^DTR^ mice. Pooled data from 2 independent experiments from 8-10 weeks-old age-and sex-matched mice. (J-M) Kaplan-Meier-survival curves (J), body weight loss in percentage of initial body weight (K and L) after bile duct ligation (BDL), and ALT levels (M) from bile duct ligated Il5-tdtomato-Cre mice and IL-5 deleter mice. Data for survival curves are pooled from 3 independent experiments with n=25 BDL Il5-tdtomato-Cre mice, n=17 BDL IL-5 deleter mice, n=10 control Il5-tdtomato-Cre mice, and n=4 IL-5 deleter mice. Other data are pooled from 3 independent experiments with n=5-13 8-10 weeks-old age-and sex-matched mice/group. Bar graphs indicate mean (±SE), Kaplan-Meier survival curves are compared using the log-rank (Mantel-Cox) analysis for J. unpaired t-test, **p ≤ 0.01.

### Supplementary Figure 5

**Figure S5:**
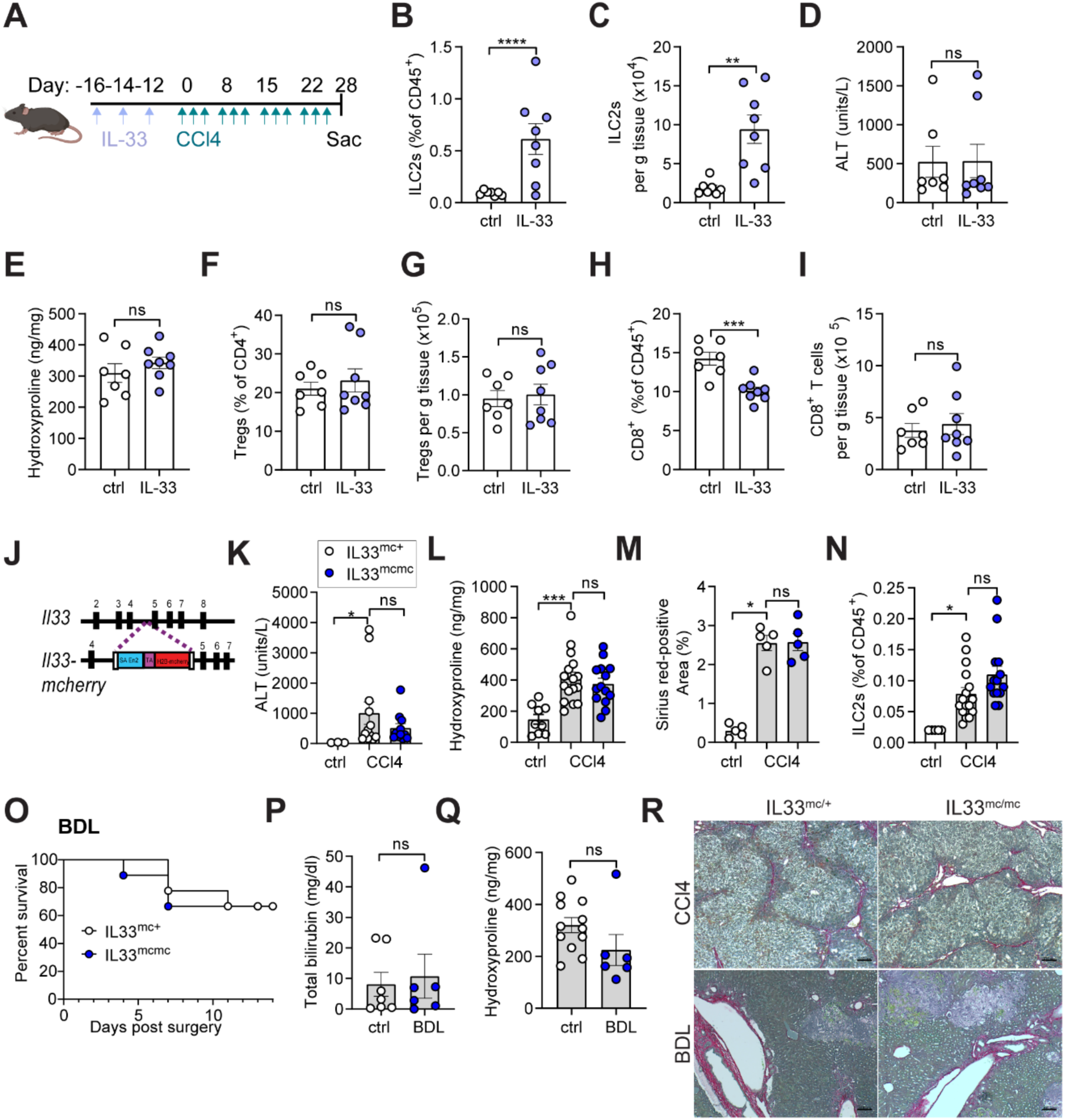
Neither ILC2 expansion nor IL-33 signalling impacted degree of hepatic fibrosis. (A) Schematic of CCl_4_ and IL-33 treatment (500ng i.p.) for IL-^33mcherry/+^ mice. (B and C) Flow cytometry quantitation of percent (B) and total numbers (C) of ILC2s in control and IL-33-treated IL-33^mcherry/+^ mice after 4 weeks of CCl_4_ treatment. (D and E) Quantification of ALT (D) and hepatic Hydroxyproline levels (E) in control and IL-33-treated IL-33^mcherry/+^ mice after 4 weeks of CCl_4_ treatment. (F-I) Flow cytometry quantitation of percent (F) or total numbers (G) of Tregs and percent (H) or total numbers (I) of CD8^+^ T cells from control and IL-33-treated IL-33^mcherry/+^ mice after 4 weeks of CCl_4_ treatment. (J) Diagram of the construction for the IL-33-H2B-mcherry nuclear localization IL-33 reporter. (K-M) Quantification of ALT (K), hepatic Hydroxyproline (L), and Sirius red staining (M) from IL-33^mcherry/+^ and IL-33^mcherry/mcherry^ mice treated with CCl_4_ for 4 weeks or controls. Pooled data from 3 independent experiments from 6-12 weeks-old age-and sex-matched mice. (N) Flow cytometry quantification, showing percent of ILC2s in livers from IL-33^mcherry/+^ and IL-33^mcherry/mcherry^ mice treated with CCl_4_ for 4 weeks or controls. Pooled data from 3 independent experiments from 6-12 weeks-old age-and sex-matched mice. (O) Kaplan–Meier survival curves after bile duct ligation (BDL). Data are pooled from 2 independent experiments with n=9 IL-33^mcherry/+^ mice and n=9 IL-33^mcherry/mcherry^ mice. (P and Q) Quantification of total bilirubin (P) and Hydroxyproline (Q) from IL-33^mcherry/+^ and IL-33^mcherry/mcherry^ mice 14 days post BDL surgery. Data are pooled from 2 independent experiments. (R) Representative Sirius red staining from IL-33^mcherry/+^ and IL-33^mcherry/mcherry^ mice treated with CCl_4_ for 4 weeks or 14 days post BDL surgery, as indicated. Bar graphs indicate mean (±SE), unpaired t-test, *p ≤ 0.05, **p ≤ 0.01, ***p ≤ 0.001, ****p ≤ 0.0001.

### Supplementary Figure 6

**Figure S6:**
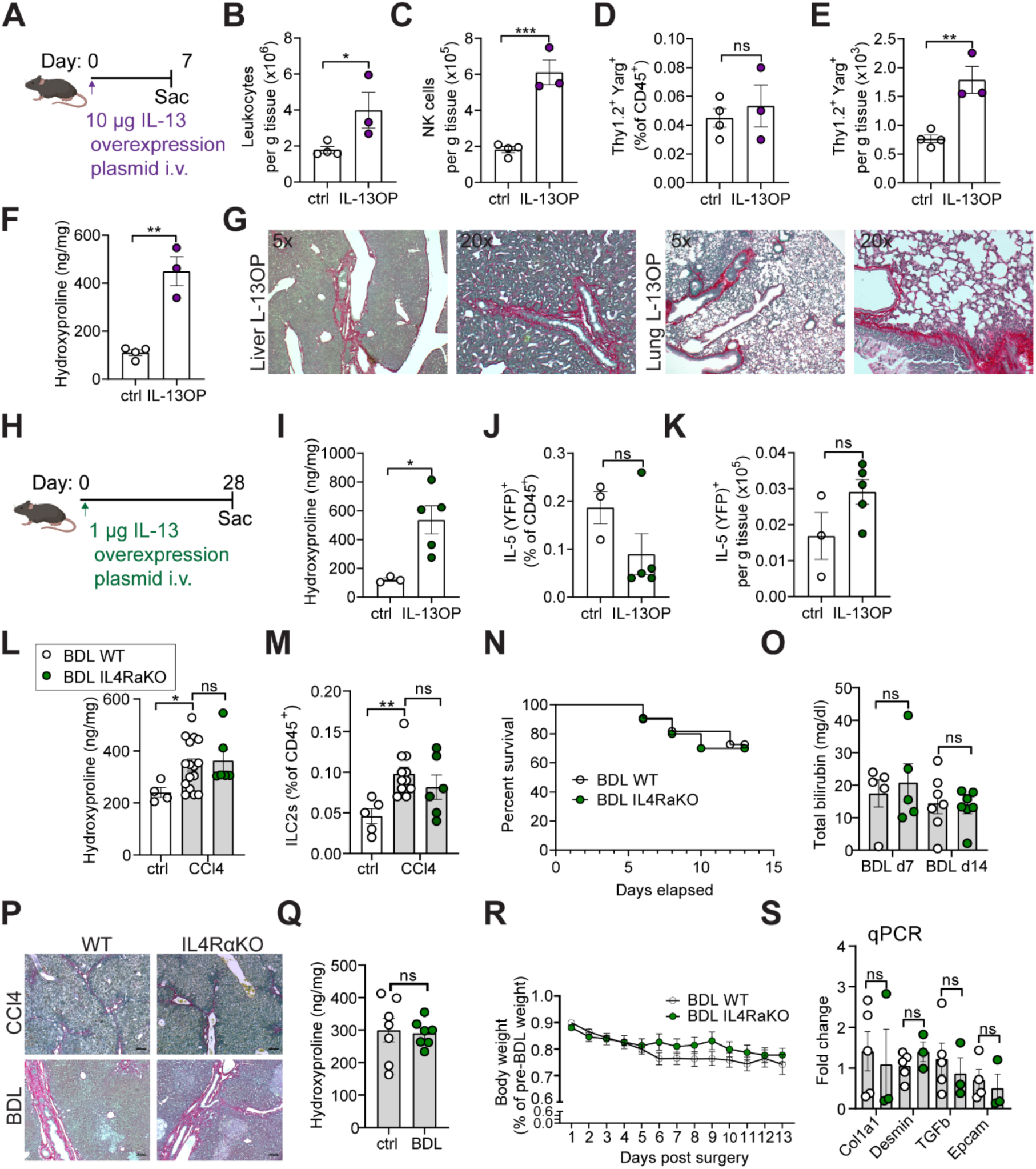
The IL-4/IL-13 receptor does not regulate hepatic fibrosis in different models of liver injury. (A) Schematic showing intravenous (i.v.) treatment with 10µg IL-13 overexpressing plasmid in Arg1 (Yarg); R5 (IL-5); S13 (IL-13) combined triple-reporter (YRS) mice. Mice were harvested after 7 days. (B-E) Flow cytometry quantitation of total leukocyte numbers (B), total numbers of NK cells (C), percent (D), and total numbers of ILC2s (Lin^-^Thy1.2^+^Arg1^+^) (E) in control and _plasmid_IL-13-treated YRS mice. (F and G) Quantification of Hydroxyproline in control and _plasmid_IL-13-treated YRS mice (F) and representative Sirius red staining from liver and lung of _plasmid_IL-13-treated YRS mice (G). Images are representative of three or more mice. (H) Schematic showing intravenous (i.v.) treatment with 1µg IL-13 overexpressing plasmid in IL-5^+^ lymphocyte lineage tracker (IL-5tdtomato-Cre; RosaCAGYFP); IL-33^mcherry/+^ double reporter mice. Mice were harvested after 4 weeks. (I) Quantification of Hydroxyproline levels in control and _plasmid_IL-13-treated IL-5^+^ lymphocyte lineage tracker; IL-33^mcherry/+^ double reporter mice. (J and K) Flow cytometry quantification of percent (J) and total numbers (K) of IL-5^+^ lymphocytes in control and _plasmid_IL-13-treated IL-5^+^ lymphocyte lineage tracker; IL-13^mcherry/+^ double reporter mice. (L and M) Quantification of Hydroxyproline (L) and flow cytometry quantification, showing percent of ILC2s (M) in liver from wild-type (WT) or IL-4/IL-13-deficient mice treated with vehicle or CCl_4_ for 4 weeks. (N) Kaplan–Meier survival curves after bile duct ligation (BDL) from wild-type and IL-4/IL-13-deficient mice. Data are pooled from 2 independent experiments with n=11 WT mice and n=10 IL-4/IL-13-deficient mice. (O) Quantification of bilirubin from WT and IL-4/IL-13-deficient mice 14 days post BDL surgery. Pooled data from 2 independent experiments from 8-10 weeks-old age-and sex-matched mice. (P) Representative Sirius red staining from WT and IL-4/IL-13-deficient mice treated with CCl_4_ for 4 weeks or 14 days post BDL surgery, as indicated. (Q) Quantification of Hydroxyproline from WT and IL-4/IL-13-deficient mice 14 days post BDL surgery. Pooled data from 2 independent experiments from 8-10 weeks-old age-and sex-matched mice. (R) Body weight loss of WT and IL-4/IL-13-deficient mice post BDL surgery. (S) Total liver gene expression of *COL1A1, DES, TGFB1*, and *EPCAM* normalized to *GAPDH* expression in control or bile duct ligated WT or IL_4/IL-13-deficient mice. Pooled data from 2 independent experiments from 8-10 weeks-old age-and sex-matched mice. Bar graphs indicate mean (±SE). Kaplan-Meier survival curves are compared using the log-rank (Mantel-Cox) analysis for N. Unpaired t-test for B-F, I-M, O, Q, and S, and two-Way ANOVA with Sidak post-test for R, *p ≤ 0.05, **p ≤ 0.01, ***p ≤ 0.001.

### Supplementary Figure 7

**Figure S7:**
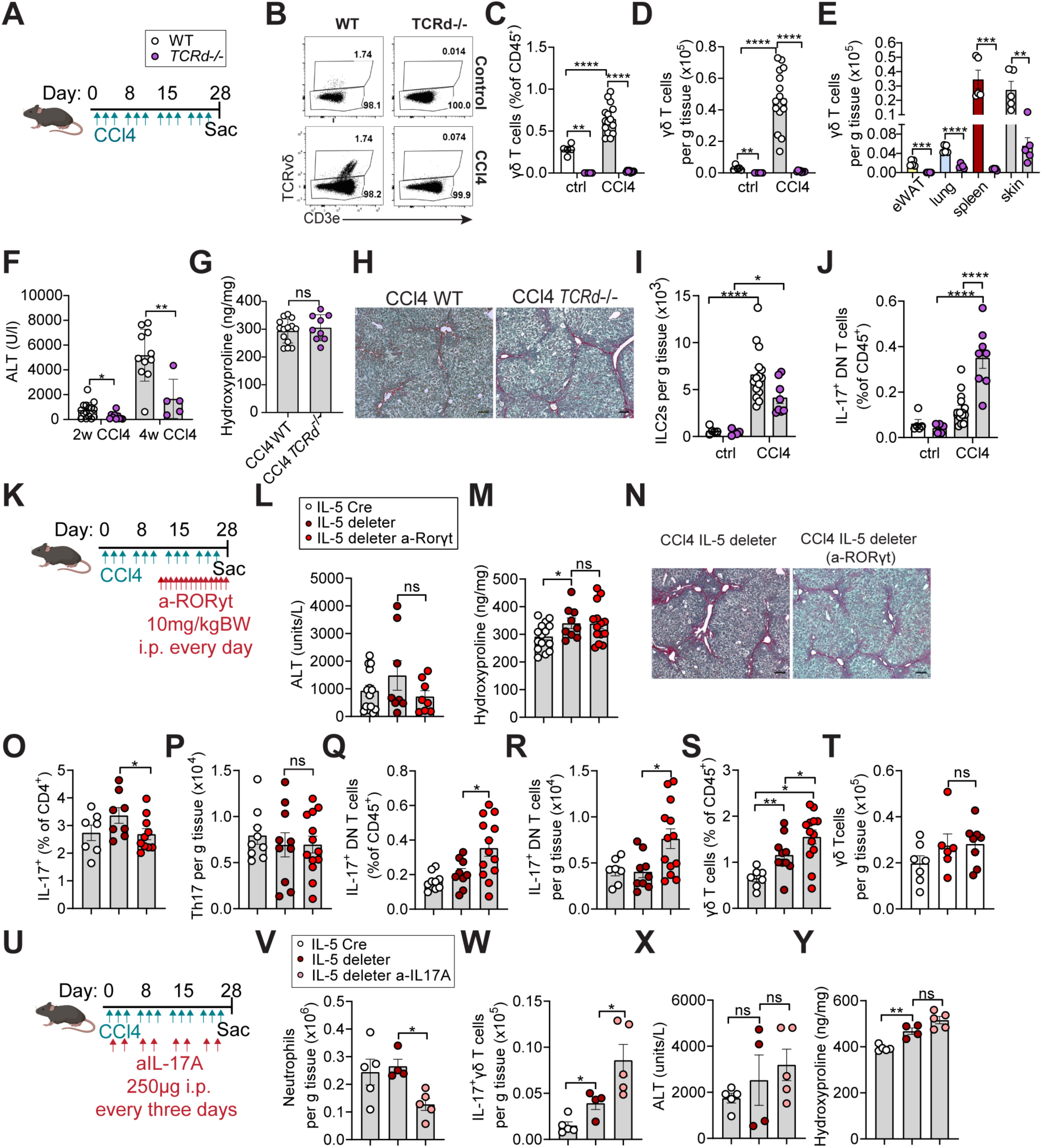
Redundancy in type 3/17 lymphocytes during hepatic fibrosis (related to **Figure 7).** (A) Schematic showing CCl_4_ administration schedule in WT and *TCRd^-/-^* mice injected intraperitoneally (i.p.) with 0.5µl CCl_4_/g BW three times per week for 4 weeks (4w), relevant to B-J. (B-D) Representative flow cytometry plots of γδ T cells (B) and quantification of percent (C) or total numbers (D) of γδ T cells in wild-type (WT) and *TCRd^-/-^* mice treated with vehicle or CCl_4_ for 4 weeks. (E) Quantification of total γδ T cell numbers in eWAT, lung, spleen, and skin in naïve adult WT and *TCRd^-/-^* mice. (F and G) Quantification of ALT levels (F) and hepatic Hydroxyproline (G) in wild-type (WT) and *TCRd*^-/-^ mice treated with CCl_4_ for indicated duration. Data pooled from 2 independent experiments with 8-10 weeks-old age-and sex-matched mice. (H) Representative Sirius red staining from WT and TCRd^-/-^ mice treated with CCl_4_ for 4 weeks. Images are representative of 2 independent experiments with 8-10 weeks old age-and sex-matched mice. (I and J) Flow cytometry quantification, showing percent (I) and total numbers of ILC2s (J) in livers from WT and TCRd^-/-^ mice treated with vehicle or CCl_4_ for 4 weeks. Pooled data from 2 independent experiments from 8-10 weeks-old age-and sex-matched mice. (K) Schematic showing treatment strategy for CCl_4_ injected intraperitoneally (i.p.) with 0.5µl CCl_4_/g BW three times per week for 4 weeks and RORγt antagonist injected with 10mg/kg BW i.p. every day for the last 2 weeks of the experiment with Il5-tdtomato-Cre mice and IL-5 deleter (RRDD; *Il5*^Red5/Red5^*Gt (Rosa)26*^DTA/DTA^), relevant to data in L-S. (L and M) Quantification of ALT levels (L) and Hydroxyproline (M) in 4-week CCl_4_-treated Il5-tdtomato-Cre mice, IL-5 deleter mice and IL-5 deleter mice treated with RORγt antagonist. Pooled data from 3 independent experiments with 8-10 weeks-old-age-and sex-matched mice. (N) Representative Sirius red staining from 4-week CCl_4_-treated IL-5 deleter and IL-5 deleter mice treated with RORγt antagonist. Images are representative of 3 independent experiments with 8-10 weeks-old age-and sex-matched mice. (O-T) Flow cytometry quantification, showing percent (O) and total numbers (P) of Th17 cells, percent (Q) and total numbers (R) of IL-7^+^ DN T cells, and percent (S) and total numbers (T) of γδ T cells in livers from Il5-tdtomato-Cre mice, IL-5 deleter, and IL-5 deleter mice treated with RORγt antagonist. Pooled data from 3 independent experiments with 8-10 weeks-old age-and sex-matched mice. (U) Schematic showing CCl_4_ and neutralizing anti-IL17A administration in Il5-tdtomato-Cre mice and IL-5 deleter mice, injected intraperitoneally (i.p.) with 0.5µl/g BW three times per week and with anti-IL17A antibody every three days throughout the experiment, relevant to U-X. (V-Y) Flow cytometry quantification of total numbers of neutrophils (V) and IL-17^+^ γδ T cells (W), and quantification of ALT levels (X) and Hydroxyproline (Y) in livers from Il5-tdtomato-Cre mice, IL-5 deleter mice, and IL-5 deleter mice treated with RORγt antagonists. Pooled data are representative of 3 independent experiments with 8-10 weeks-old age-and sex-matched mice. Bar graphs indicate mean (±SE), unpaired t-test for C-G, I, J, L, M, and one-Way ANOVA for O-S and U-X, ns= not significant, *p ≤ 0.05, **p ≤ 0.01, ***p ≤ 0.001, ****p ≤ 0. 0001.

## Notes

### Competing Interest Statement

The authors have declared no competing interest.

## REFERENCES AND NOTES

Allen, J.E., and Sutherland, T.E. (2014). Host protective roles of type 2 immunity: parasite killing and tissue repair, flip sides of the same coin. Semin. Immunol. 26, 329–340.

Anderson, K.G., Mayer-Barber, K., Sung, H., Beura, L., James, B.R., Taylor, J.J., Qunaj, L., Griffith, T.S., Vezys, V., Barber, D.L., et al. (2014). Intravascular staining for discrimination of vascular and tissue leukocytes. Nat. Protoc. 9, 209–222.

Annunziato, F., Romagnani, C., and Romagnani, S. (2015). The 3 major types of innate and adaptive cell-mediated effector immunity. J. Allergy Clin. Immunol. 135, 626–635.

Boothby, I.C., Kinet, M.J., Boda, D.P., Kwan, E.Y., Clancy, S., Cohen, J.N., Habrylo, I., Lowe, M.M., Pauli, M., Yates, A.E., et al. (2021). Early-life inflammation primes a T helper 2 cell–fibroblast niche in skin. Nature 599, 667–672.

Buch, T., Heppner, F.L., Tertilt, C., Heinen, T.J.A.J., Kremer, M., Wunderlich, F.T., Jung, S., and Waisman, A. (2005). A Cre-inducible diphtheria toxin receptor mediates cell lineage ablation after toxin administration. Nat. Methods 2, 419–426.

Buechler, M.B., Pradhan, R.N., Krishnamurty, A.T., Cox, C., Calviello, A.K., Wang, A.W., Yang, Y.A., Tam, L., Caothien, R., Roose-Girma, M., et al. (2021). Cross-tissue organization of the fibroblast lineage. Nature 593, 575–579.

Di Carlo, S.E., and Peduto, L. (2018). The perivascular origin of pathological fibroblasts. J. Clin. Invest. 128, 54–63.

Cassandras, M., Wang, C., Kathiriya, J., Tsukui, T., Matatia, P., Matthay, M., Wolters, P., Molofsky, A., Sheppard, D., Chapman, H., et al. (2020). Gli1+ mesenchymal stromal cells form a pathological niche to promote airway progenitor metaplasia in the fibrotic lung. Nat. Cell Biol. 22, 1295–1306.

Cautivo, K.M., Steer, C.A., and Molofsky, A.B. (2020). Immune outposts in the adventitia: One foot in sea and one on shore. Curr. Opin. Immunol. 64, 34–41.

Cautivo, K.M., Matatia, P.R., Lizama, C.O., Mroz, N.M., Dahlgren, M.W., Yu, X., Sbierski-Kind, J., Taruselli, M.T., Brooks, J.F., Wade-Vallance, A., et al. (2022). Interferon gamma constrains type 2 lymphocyte niche boundaries during mixed inflammation. Immunity 55, 254–271.e7.

Choy, D.F., Hart, K.M., Borthwick, L.A., Shikotra, A., Nagarkar, D.R., Siddiqui, S., Jia, G., Ohri, C.M., Doran, E., Vannella, K.M., et al. (2015). T^H^2 and T^H^17 inflammatory pathways are reciprocally regulated in asthma. Sci. Transl. Med. 7, 301ra129–LP-301ra129.

Dahlgren, M.W., and Molofsky, A.B. (2019). Adventitial Cuffs: Regional Hubs for Tissue Immunity. Trends Immunol. 40, 877–887.

Dahlgren, M.W., Jones, S.W., Cautivo, K.M., Dubinin, A., Ortiz-Carpena, J.F., Farhat, S., Yu, K.S., Lee, K., Wang, C., Molofsky, A. V., et al. (2019). Adventitial Stromal Cells Define Group 2 Innate Lymphoid Cell Tissue Niches. Immunity 50, 707–722.e6.

Degryse, A.L., Tanjore, H., Xu, X.C., Polosukhin, V. V, Jones, B.R., McMahon, F.B., Gleaves, L.A., Blackwell, T.S., and Lawson, W.E. (2010). Repetitive intratracheal bleomycin models several features of idiopathic pulmonary fibrosis. Am. J. Physiol. Lung Cell. Mol. Physiol. 299, L442–L452.

Dobie, R., Wilson-Kanamori, J.R., Henderson, B.E.P., Smith, J.R., Matchett, K.P., Portman, J.R., Wallenborg, K., Picelli, S., Zagorska, A., Pendem, S. V, et al. (2019a). Single-Cell Transcriptomics Uncovers Zonation of Function in the Mesenchyme during Liver Fibrosis. Cell Rep. 29, 1832–1847.e8.

Dobie, R., Wilson-Kanamori, J.R., Henderson, B.E.P., Smith, J.R., Matchett, K.P., Portman, J.R., Wallenborg, K., Picelli, S., Zagorska, A., Pendem, S. V., et al. (2019b). Single-Cell Transcriptomics Uncovers Zonation of Function in the Mesenchyme during Liver Fibrosis. Cell Rep. 29, 1832–1847.e8.

Van Dyken, S.J., and Locksley, R.M. (2013). Interleukin-4-and interleukin-13-mediated alternatively activated macrophages: roles in homeostasis and disease. Annu. Rev. Immunol. 31, 317–343.

Fabre, T., Kared, H., Friedman, S.L., and Shoukry, N.H. (2014). IL-17A Enhances the Expression of Profibrotic Genes through Upregulation of the TGF-β Receptor on Hepatic Stellate Cells in a JNK-Dependent Manner. J. Immunol. 193, 3925–3933.

Fabre, T., Molina, M.F., Soucy, G., Goulet, J.P., Willems, B., Villeneuve, J.P., Bilodeau, M., and Shoukry, N.H. (2018). Type 3 cytokines IL-17A and IL-22 drive TGF-–dependent liver fibrosis. Sci. Immunol. 3, 1–16.

Fabre, T., Barron, A.M.S., Christensen, S.M., Asano, S., Wadsworth, M.H., Chen, X., Wang, J., McMahon, J., Schlerman, F., White, A., et al. (2022). Identification of a Broadly Fibrogenic Macrophage Subset Induced by Type 3 Inflammation in Human and Murine Liver and Lung Fibrosis. BioRxiv 2022.07.01.498017.

Fan, X., and Rudensky, A.Y. (2016). Hallmarks of Tissue-Resident Lymphocytes. Cell 164, 1198–1211.

Faustino, L.D., Griffith, J.W., Rahimi, R.A., Nepal, K., Hamilos, D.L., Cho, J.L., Medoff, B.D., Moon, J.J., Vignali, D.A.A., and Luster, A.D. (2020). Interleukin-33 activates regulatory T cells to suppress innate γδ T cell responses in the lung. Nat. Immunol. 21, 1371–1383.

Forkel, M., Berglin, L., Kekäläinen, E., Carlsson, A., Svedin, E., Michaëlsson, J., Nagasawa, M., Erjefält, J.S., Mori, M., Flodström-Tullberg, M., et al. (2017). Composition and functionality of the intrahepatic innate lymphoid cell-compartment in human nonfibrotic and fibrotic livers. Eur. J. Immunol. 47, 1280–1294.

Fort, M.M., Cheung, J., Yen, D., Li, J., Zurawski, S.M., Lo, S., Menon, S., Clifford, T., Hunte, B., Lesley, R., et al. (2001). IL-25 Induces IL-4, IL-5, and IL-13 and Th2-Associated Pathologies In Vivo. Immunity 15, 985–995.

Gao, Y., Liu, Y., Yang, M., Guo, X., Zhang, M., Li, H., Li, J., and Zhao, J. (2016). IL-33 treatment attenuated diet-induced hepatic steatosis but aggravated hepatic fibrosis. Oncotarget 7, 33649–33661.

Gasteiger, G., Fan, X., Dikiy, S., Lee, S.Y., and Rudensky, A.Y. (2015). Tissue residency of innate lymphoid cells in lymphoid and nonlymphoid organs. Science (80-.). 350, 981 LP – 985.

Ghallab, A., Myllys, M., Holland, C.H., Zaza, A., Murad, W., Hassan, R., Ahmed, Y.A., Abbas, T., Abdelrahim, E.A., Schneider, K.M., et al. (2019). Influence of Liver Fibrosis on Lobular Zonation. Cells 8, 1556.

Gieseck, R.L., Ramalingam, T.R., Hart, K.M., Vannella, K.M., Cantu, D.A., Lu, W.Y., Ferreira-González, S., Forbes, S.J., Vallier, L., and Wynn, T.A. (2016). Interleukin-13 Activates Distinct Cellular Pathways Leading to Ductular Reaction, Steatosis, and Fibrosis. Immunity 45, 145–158.

Gieseck, R.L., Wilson, M.S., and Wynn, T.A. (2018a). Type 2 immunity in tissue repair and fibrosis. Nat. Rev. Immunol. 18, 62–76.

Gieseck, R.L., Wilson, M.S., and Wynn, T.A. (2018b). Type 2 immunity in tissue repair and fibrosis. Nat. Rev. Immunol. 18, 62–76.

Gola, A., Dorrington, M.G., Speranza, E., Sala, C., Shih, R.M., Radtke, A.J., Wong, H.S., Baptista, A.P., Hernandez, J.M., Castellani, G., et al. (2020). Commensal-driven immune zonation of the liver promotes host defence. Nature.

Guilliams, M., Bonnardel, J., Haest, B., Vanderborght, B., Wagner, C., Remmerie, A., Bujko, A., Martens, L., Thoné, T., Browaeys, R., et al. (2022). Spatial proteogenomics reveals distinct and evolutionarily conserved hepatic macrophage niches. Cell 185, 379–396.e38.

Hams, E., Armstrong, M.E., Barlow, J.L., Saunders, S.P., Schwartz, C., Cooke, G., Fahy, R.J., Crotty, T.B., Hirani, N., Flynn, R.J., et al. (2014). IL-25 and type 2 innate lymphoid cells induce pulmonary fibrosis. Proc. Natl. Acad. Sci. U. S. A. 111, 367– 372.

Hart, K.M., Fabre, T., Sciurba, J.C., Gieseck, R.L., Borthwick, L.A., Vannella, K.M., Acciani, T.H., De Queiroz Prado, R., Thompson, R.W., White, S., et al. (2017). Type 2 immunity is protective in metabolic disease but exacerbates NAFLD collaboratively with TGF-b. Sci. Transl. Med. 9.

Hemmers, S., Schizas, M., and Rudensky, A.Y. (2021). T reg cell-intrinsic requirements for ST2 signaling in health and neuroinflammation. J. Exp. Med. 218.

Hernandez-Gea, V., and Friedman, S.L. (2011). Pathogenesis of Liver Fibrosis. Annu. Rev. Pathol. Mech. Dis. 6, 425–456.

Hirota, K., Duarte, J.H., Veldhoen, M., Hornsby, E., Li, Y., Cua, D.J., Ahlfors, H., Wilhelm, C., Tolaini, M., Menzel, U., et al. (2011). Fate mapping of IL-17-producing T cells in inflammatory responses. Nat. Immunol. 12, 255–263.

Hurst, S.D., Muchamuel, T., Gorman, D.M., Gilbert, J.M., Clifford, T., Kwan, S., Menon, S., Seymour, B., Jackson, C., Kung, T.T., et al. (2002). New IL-17 Family Members Promote Th1 or Th2 Responses in the Lung: In Vivo Function of the Novel Cytokine IL-25. J. Immunol. 169, 443 LP – 453.

Ivanov, I.I., McKenzie, B.S., Zhou, L., Tadokoro, C.E., Lepelley, A., Lafaille, J.J., Cua, D.J., and Littman, D.R. (2006). The Orphan Nuclear Receptor RORγt Directs the Differentiation Program of Proinflammatory IL-17^+^ T Helper Cells. Cell 126, 1121–1133.

Kalekar, L.A., Cohen, J.N., Prevel, N., Sandoval, P.M., Mathur, A.N., Moreau, J.M., Lowe, M.M., Nosbaum, A., Wolters, P.J., Haemel, A., et al. (2019). Regulatory T cells in skin are uniquely poised to suppress profibrotic immune responses. Sci. Immunol. 4, eaaw2910.

Karo-Atar, D., Bordowitz, A., Wand, O., Pasmanik-Chor, M., Fernandez, I.E., Itan, M., Frenkel, R., Herbert, D.R., Finkelman, F.D., Eickelberg, O., et al. (2016). A protective role for IL-13 receptor α 1 in bleomycin-induced pulmonary injury and repair. Mucosal Immunol. 9, 240–253.

Kim, B.S., Siracusa, M.C., Saenz, S.A., Noti, M., Monticelli, L.A., Sonnenberg, G.F., Hepworth, M.R., Van Voorhees, A.S., Comeau, M.R., and Artis, D. (2013). TSLP elicits IL-33-independent innate lymphoid cell responses to promote skin inflammation. Sci. Transl. Med. 5, 170ra16.

Kleiner, D.E., and Brunt, E.M. (2012). Nonalcoholic Fatty Liver Disease: Pathologic Patterns and Biopsy Evaluation in Clinical Research. Semin Liver Dis 32, 3–13.

Lee, M.W., Odegaard, J.I., Mukundan, L., Qiu, Y., Molofsky, A.B., Nussbaum, J.C., Yun, K., Locksley, R.M., and Chawla, A. (2015). Activated type 2 innate lymphoid cells regulate beige fat biogenesis. Cell 160, 74–87.

Liang, H.-E., Reinhardt, R.L., Bando, J.K., Sullivan, B.M., Ho, I.-C., and Locksley, R.M. (2012). Divergent expression patterns of IL-4 and IL-13 define unique functions in allergic immunity. Nat. Immunol. 13, 58–66.

Lloyd, C.M., and Snelgrove, R.J. (2018). Type 2 immunity: Expanding our view. Sci. Immunol. 3, eaat1604.

Lochner, M., Peduto, L., Cherrier, M., Sawa, S., Langa, F., Varona, R., Riethmacher, D., Si-Tahar, M., Di Santo, J.P., and Eberl, G. (2008). In vivo equilibrium of proinflammatory IL-17+ and regulatory IL-10+ Foxp3+ RORgamma t+ T cells. J. Exp. Med. 205, 1381–1393.

Madisen, L., Zwingman, T.A., Sunkin, S.M., Oh, S.W., Zariwala, H.A., Gu, H., Ng, L.L., Palmiter, R.D., Hawrylycz, M.J., Jones, A.R., et al. (2010). A robust and high-throughput Cre reporting and characterization system for the whole mouse brain. Nat. Neurosci. 13, 133–140.

Magdaleno-Tapial, J., López-Martí, C., Ortiz-Salvador, J.M., Hernández-Bel, P., Tamarit-García, J.J., Diago-Madrid, M., Sánchez-Carazo, J.L., and Pérez-Ferriols, A. (2021). Can secukinumab improve liver fibrosis? A pilot prospective study of 10 psoriatic patients. Dermatol. Ther. 34, e15065.

Marvie, P., Lisbonne, M., L’Helgoualc’h, A., Rauch, M., Turlin, B., Preisser, L., Bourd-Boittin, K., Théret, N., Gascan, H., Piquet-Pellorce, C., et al. (2010). Interleukin-33 overexpression is associated with liver fibrosis in mice and humans. J. Cell. Mol. Med. 14, 1726–1739.

Mchedlidze, T., Waldner, M., Zopf, S., Walker, J., Rankin, A.L., Schuchmann, M., Voehringer, D., McKenzie, A.N.J., Neurath, M.F., Pflanz, S., et al. (2013). Interleukin-33-dependent innate lymphoid cells mediate hepatic fibrosis. Immunity 39, 357–371.

Meng, F., Wang, K., Aoyama, T., Grivennikov, S.I., Paik, Y., Scholten, D., Cong, M., Iwaisako, K., Liu, X., Zhang, M., et al. (2012). Interleukin-17 signaling in inflammatory, Kupffer cells, and hepatic stellate cells exacerbates liver fibrosis in mice. Gastroenterology 143, 765–776.e3.

Mohrs, M., Ledermann, B., Köhler, G., Dorfmüller, A., Gessner, A., and Brombacher, F. (1999). Differences Between IL-4-and IL-4 Receptor α-Deficient Mice in Chronic Leishmaniasis Reveal a Protective Role for IL-13 Receptor Signaling. J. Immunol. 162, 7302 LP – 7308.

Molina, M.F., Abdelnabi, M.N., Fabre, T., and Shoukry, N.H. (2019). Type 3 cytokines in liver fibrosis and liver cancer. Cytokine 124, 154497.

Molofsky, A.B., Nussbaum, J.C., Liang, H.E., Dyken, S.J.V., Cheng, L.E., Mohapatra, A., Chawla, A., and Locksley, R.M. (2013). Innate lymphoid type 2 cells sustain visceral adipose tissue eosinophils and alternatively activated macrophages. J. Exp. Med.

Molofsky, A.B., Savage, A.K., and Locksley, R.M. (2015). Interleukin-33 in Tissue Homeostasis, Injury, and Inflammation. Immunity.

Moro, K., Yamada, T., Tanabe, M., Takeuchi, T., Ikawa, T., Kawamoto, H., Furusawa, J., Ohtani, M., Fujii, H., and Koyasu, S. (2010). Innate production of TH2 cytokines by adipose tissue-associated c-Kit+Sca-1+ lymphoid cells. Nature 463, 540–544.

Neill, D.R., Wong, S.H., Bellosi, A., Flynn, R.J., Daly, M., Langford, T.K.A., Bucks, C., Kane, C.M., Fallon, P.G., Pannell, R., et al. (2010). Nuocytes represent a new innate effector leukocyte that mediates type-2 immunity. Nature 464, 1367–1370.

Nussbaum, J.C., Van Dyken, S.J., von Moltke, J., Cheng, L.E., Mohapatra, A., Molofsky, A.B., Thornton, E.E., Krummel, M.F., Chawla, A., Liang, H.-E., et al. (2013). Type 2 innate lymphoid cells control eosinophil homeostasis. Nature 502, 245—248.

Price, A.E., Liang, H.-E., Sullivan, B.M., Reinhardt, R.L., Eisley, C.J., Erle, D.J., and Locksley, R.M. (2010). Systemically dispersed innate IL-13-expressing cells in type 2 immunity. Proc. Natl. Acad. Sci. U. S. A. 107, 11489–11494.

Ramachandran, P., Dobie, R., Wilson-Kanamori, J.R., Dora, E.F., Henderson, B.E.P., Luu, N.T., Portman, J.R., Matchett, K.P., Brice, M., Marwick, J.A., et al. (2019). Resolving the fibrotic niche of human liver cirrhosis at single-cell level. Nature 575, 512–518.

Rau, M., Schilling, A.-K., Meertens, J., Hering, I., Weiss, J., Jurowich, C., Kudlich, T., Hermanns, H.M., Bantel, H., Beyersdorf, N., et al. (2016). Progression from Nonalcoholic Fatty Liver to Nonalcoholic Steatohepatitis Is Marked by a Higher Frequency of Th17 Cells in the Liver and an Increased Th17/Resting Regulatory T Cell Ratio in Peripheral Blood and in the Liver. J. Immunol. 196, 97–105.

Reese, T.A., Liang, H.-E., Tager, A.M., Luster, A.D., Van Rooijen, N., Voehringer, D., and Locksley, R.M. (2007). Chitin induces accumulation in tissue of innate immune cells associated with allergy. Nature 447, 92–96.

Remmerie, A., Martens, L., Thoné, T., Castoldi, A., Seurinck, R., Pavie, B., Roels, J., Vanneste, B., De Prijck, S., Vanhockerhout, M., et al. (2020). Osteopontin Expression Identifies a Subset of Recruited Macrophages Distinct from Kupffer Cells in the Fatty Liver. Immunity 53, 641–657.e14.

Ricardo-Gonzalez, R.R., Van Dyken, S.J., Schneider, C., Lee, J., Nussbaum, J.C., Liang, H.-E., Vaka, D., Eckalbar, W.L., Molofsky, A.B., Erle, D.J., et al. (2018). Tissue signals imprint ILC2 identity with anticipatory function. Nat. Immunol. 19, 1093–1099.

Rustenhoven, J., Drieu, A., Mamuladze, T., Lopes, M., and Herz, J. (2021). Functional characterization of the dural sinuses as a neuroimmune interface. Cell 1– 17.

Ruterbusch, M., Pruner, K.B., Shehata, L., and Pepper, M. (2020). In Vivo CD4+ T Cell Differentiation and Function: Revisiting the Th1/Th2 Paradigm. Annu. Rev. Immunol. 38, 705–725.

Saito*, J.M., and Maher‡, J.J. (2000). Bile duct ligation in rats induces biliary expression of cytokine-induced neutrophil chemoattractant. Gastroenterology 118, 1157–1168.

Saluzzo, S., Gorki, A.-D., Rana, B.M.J., Martins, R., Scanlon, S., Starkl, P., Lakovits, K., Hladik, A., Korosec, A., Sharif, O., et al. (2017). First-Breath-Induced Type 2 Pathways Shape the Lung Immune Environment. Cell Rep. 18, 1893–1905.

Sbierski-Kind, J., Mroz, N., and Molofsky, A.B. (2021). Perivascular stromal cells: Directors of tissue immune niches. Immunol. Rev.

Schneider, C., Lee, J., Koga, S., Ricardo-Gonzalez, R.R., Nussbaum, J.C., Smith, L.K., Villeda, S.A., Liang, H.E., and Locksley, R.M. (2019). Tissue-Resident Group 2 Innate Lymphoid Cells Differentiate by Layered Ontogeny and In Situ Perinatal Priming. Immunity 50, 1425–1438.e5.

Suda, T., and Liu, D. (2007). Hydrodynamic Gene Delivery: Its Principles and Applications. Mol. Ther. 15, 2063–2069.

Tan, Z., Qian, X., Jiang, R., Liu, Q., Wang, Y., Chen, C., Wang, X., Ryffel, B., and Sun, B. (2013). IL-17A Plays a Critical Role in the Pathogenesis of Liver Fibrosis through Hepatic Stellate Cell Activation. J. Immunol. 191, 1835 LP – 1844.

Tan, Z., Liu, Q., Jiang, R., Lv, L., Shoto, S.S., Maillet, I., Quesniaux, V., Tang, J., Zhang, W., Sun, B., et al. (2018). Interleukin-33 drives hepatic fibrosis through activation of hepatic stellate cells. Cell. Mol. Immunol. 15, 388–398.

Tedesco, D., Thapa, M., Chin, C.Y., Ge, Y., Gong, M., Li, J., Gumber, S., Speck, P., Elrod, E.J., Burd, E.M., et al. (2018). Alterations in Intestinal Microbiota Lead to Production of Interleukin 17 by Intrahepatic γδ T-Cell Receptor-Positive Cells and Pathogenesis of Cholestatic Liver Disease. Gastroenterology 154, 2178–2193.

Tortola, L., Jacobs, A., Pohlmeier, L., Obermair, F.-J., Ampenberger, F., Bodenmiller, B., and Kopf, M. (2020). High-Dimensional T Helper Cell Profiling Reveals a Broad Diversity of Stably Committed Effector States and Uncovers Interlineage Relationships. Immunity 53, 597–613.e6.

Tsuchida, T., Lee, Y.A., Fujiwara, N., Ybanez, M., Allen, B., Martins, S., Fiel, M.I., Goossens, N., Chou, H.-I., Hoshida, Y., et al. (2018). A simple diet-and chemical-induced murine NASH model with rapid progression of steatohepatitis, fibrosis and liver cancer. J. Hepatol. 69, 385–395.

Tsukui, T., Sun, K.H., Wetter, J.B., Wilson-Kanamori, J.R., Hazelwood, L.A., Henderson, N.C., Adams, T.S., Schupp, J.C., Poli, S.D., Rosas, I.O., et al. (2020). Collagen-producing lung cell atlas identifies multiple subsets with distinct localization and relevance to fibrosis. Nat. Commun. 11, 1–16.

Vainchtein, I.D., Chin, G., Cho, F.S., Kelley, K.W., Miller, J.G., Chien, E.C., Liddelow, S.A., Nguyen, P.T., Nakao-Inoue, H., Dorman, L.C., et al. (2018a). Astrocyte-derived interleukin-33 promotes microglial synapse engulfment and neural circuit development. Science (80-.). 359, 1269–1273.

Vainchtein, I.D., Chin, G., Cho, F.S., Kelley, K.W., Miller, J.G., Chien, E.C., Liddelow, S.A., Nguyen, P.T., Nakao-Inoue, H., Dorman, L.C., et al. (2018b). Astrocyte-derived interleukin-33 promotes microglial synapse engulfment and neural circuit development. Science (80-.). 359, 1269 LP – 1273.

Van Dyken, S.J., Mohapatra, A., Nussbaum, J.C., Molofsky, A.B., Thornton, E.E., Ziegler, S.F., McKenzie, A.N.J., Krummel, M.F., Liang, H.-E., and Locksley, R.M. (2014). Chitin Activates Parallel Immune Modules that Direct Distinct Inflammatory Responses via Innate Lymphoid Type 2 and γδ T Cells. Immunity 40, 414–424.

Vannella, K.M., Ramalingam, T.R., Borthwick, L.A., Barron, L., Hart, K.M., Thompson, R.W., Kindrachuk, K.N., Cheever, A.W., White, S., Budelsky, A.L., et al. (2016). Combinatorial targeting of TSLP, IL-25, and IL-33 in type 2 cytokine-driven inflammation and fibrosis. Sci. Transl. Med. 8, 1–13.

Wang, X., Zheng, Z., Caviglia, J.M., Corey, K.E., Herfel, T.M., Cai, B., Masia, R., Chung, R.T., Lefkowitch, J.H., Schwabe, R.F., et al. (2016). Hepatocyte TAZ/WWTR1 Promotes Inflammation and Fibrosis in Nonalcoholic Steatohepatitis. Cell Metab. 24, 848–862.

Wilson, M.S., Madala, S.K., Ramalingam, T.R., Gochuico, B.R., Rosas, I.O., Cheever, A.W., and Wynn, T.A. (2010). Bleomycin and IL-1beta-mediated pulmonary fibrosis is IL-17A dependent. J. Exp. Med. 207, 535–552.

Withers, D.R., Hepworth, M.R., Wang, X., Mackley, E.C., Halford, E.E., Dutton, E.E., Marriott, C.L., Brucklacher-Waldert, V., Veldhoen, M., Kelsen, J., et al. (2016). Transient inhibition of ROR-γt therapeutically limits intestinal inflammation by reducing TH17 cells and preserving group 3 innate lymphoid cells. Nat. Med. 22, 319–323.

Wree, A., McGeough, M.D., Inzaugarat, M.E., Eguchi, A., Schuster, S., Johnson, C.D., Peña, C.A., Geisler, L.J., Papouchado, B.G., Hoffman, H.M., et al. (2018). NLRP3 inflammasome driven liver injury and fibrosis: Roles of IL-17 and TNF in mice. Hepatology 67, 736–749.

Wynn, T.A., and Vannella, K.M. (2016). Macrophages in Tissue Repair, Regeneration, and Fibrosis. Immunity 44, 450–462.

Zeis, P., Lian, M., Fan, X., Herman, J.S., Hernandez, D.C., Gentek, R., Elias, S., Symowski, C., Knöpper, K., Peltokangas, N., et al. (2020). In Situ Maturation and Tissue Adaptation of Type 2 Innate Lymphoid Cell Progenitors. Immunity 53, 775–792.e9.

Zhu, J., Jankovic, D., Oler, A.J., Wei, G., Sharma, S., Hu, G., Guo, L., Yagi, R., Yamane, H., Punkosdy, G., et al. (2012). The transcription factor T-bet is induced by multiple pathways and prevents an endogenous Th2 cell program during Th1 cell responses. Immunity 37, 660–673.

