## Supplementary Information for "Group 2 innate lymphoid cells constrain type 3/17 lymphocytes in shared stromal niches to restrict liver fibrosis"

### Supplemental information

#### Supplementary Figure 1

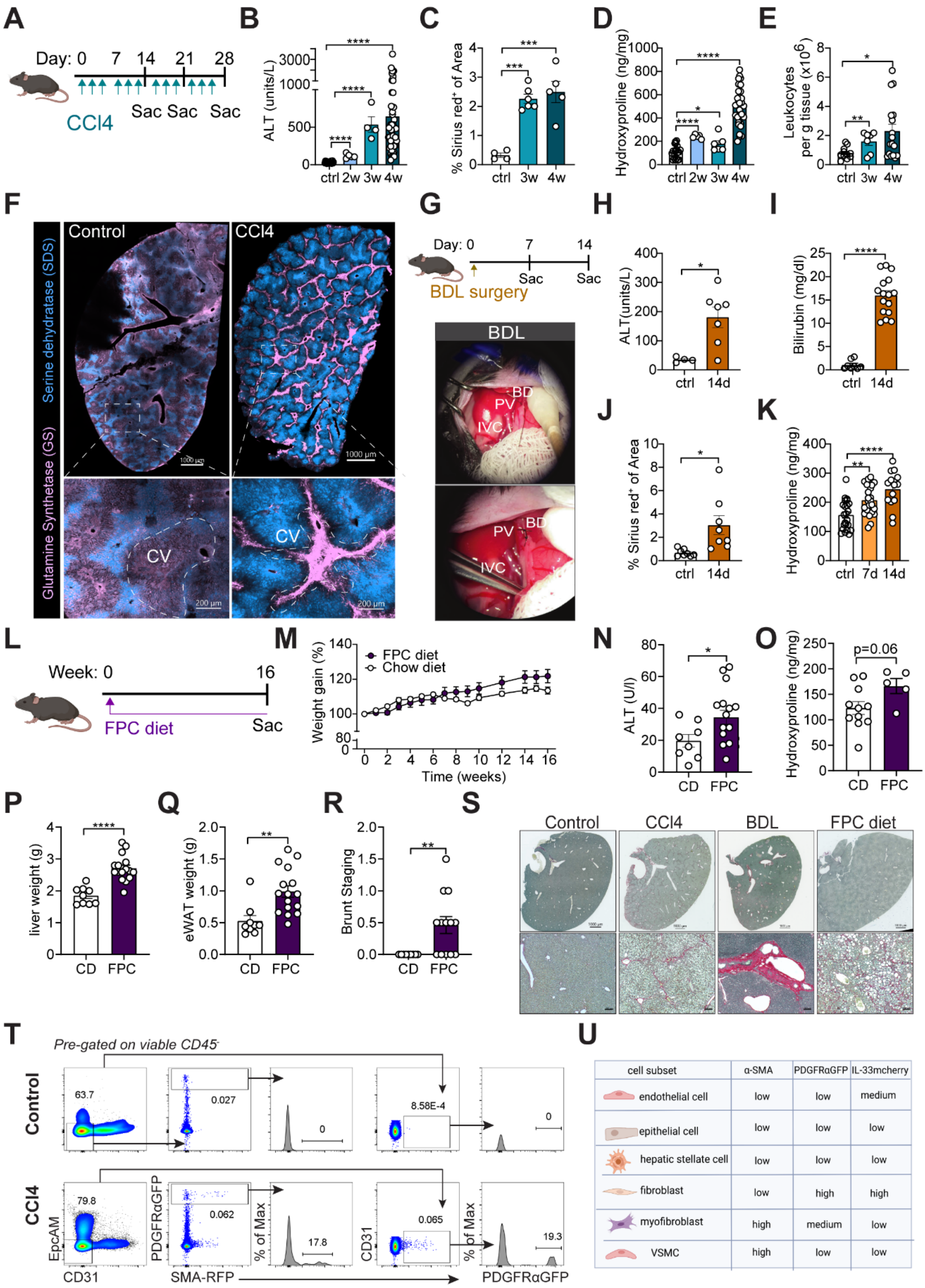


| cell subset | α-SMA | PDGFRαGFP | IL-33mcherry |
|---|---|---|---|
| endothelial cell | low | low | medium |
| epithelial cell | low | low | low |
| hepatic stellate cell | low | low | low |
| fibroblast | low | high | high |
| myofibroblast | high | medium | low |
| VSMC | high | low | low |

**Figure S1: Models of liver fibrosis and impact on stromal cell topography (related to Figure 1).**

**(A)** Schematic showing $CCl_4$ administration schedule in IL-33$^{mcherry/+}$ reporter mice injected intraperitoneally (i.p.) with 0.5µl $CCl_4$/g BW three times per week for 2, 3, or 4 weeks (2w, 3w, 4w), relevant to B-E.

**(B-E)** Quantification of ALT **(B)**, Sirius red positive staining **(C)**, hepatic Hydroxyproline levels **(D),** total leukocytes **(E)** in control mice and $CCl_4$-treated mice at the indicated timepoints. Pooled data from 8-10 weeks old age- and sex-matched mice for 4-week timepoint.

**(F)** Representative thick section liver image from control and 4-week $CCl_4$ treated IL-33$^{mcherry/+}$ reporter mice with staining for glutamin synthetase (GS), a marker for pericentral hepatocytes, and sodium dodecyl sulphate (SDS) surrounding portal veins. Higher magnification image highlighting severe structural changes in liver morphology with $CCl_4$-induced fibrosis.

**(G)** Schematic of bile duct ligation (BDL) surgery in PDGFRαGFP; IL33$^{mcherry/+}$ reporter mice for time course analysis, relevant to H-K. Bile duct (BDL), Portal vein (PV), Inferior vena cava (IVC).

**(H-K)** Quantification of ALT **(H)**, Bilirubin **(I)**, Sirius red positive staining **(J)**, and hepatic Hydroxyproline levels **(K)** in control mice and bile duct ligated mice at the indicated timepoints. Pooled data from 3 independent experiments from 8-10-weeks-old age- and sex-matched mice.

**(L)** Schematic of fructose-palmitate-cholesterol (FPC) diet in IL-33$^{mcherry/+}$ mice, relevant to M-S.

**(M-R)** Body weight gain **(M)**, quantification of ALT **(N)** and hepatic Hydroxyproline levels **(O)**, liver weight **(P)**, epigonadal white adipose tissue (eWAT) weight **(Q)** and Brunt staging **(R)** in 16w chow diet-fed and 16w FPC diet-fed mice. Data pooled from 3 independent experiments from 6-8-weeks-old IL-33$^{mcherry/+}$ reporter age- and sex-matched mice.

**(S)** Representative Sirius Red staining of 5µm paraffin liver sections from IL-33$^{mcherry/+}$ reporter mice at steady state, at day 28 of $CCl_4$ treatment, at day 14 post BDL surgery, and at week 16 of FPC diet feeding. Higher magnification images show detailed morphology of collagen deposition. Scale Bars: 100µm (10X). Images are representative of 3 repeat experiments with n=3-10 mice per group.

**(T)** Representative flow cytometry plots from liver of PDGFRαGFPSMARFP reporter mice treated with vehicle or $CCl_4$ for 4 weeks and representative histograms of αSMA expression on PDGFRαGFP$^+$ fibroblasts. Data are representative of 2 independent experiments.

**(U)** Schematic showing expression of α-SMA, PDGFRαGFP and IL-33mcherry in different stromal cell subsets.

Bar graphs indicate mean (±SE). Unpaired t-test, *$p \le 0.05$, **$p \le 0.01$, ***$p \le 0.001$, ****$p \le 0.0001$.

#### Supplementary Figure 2

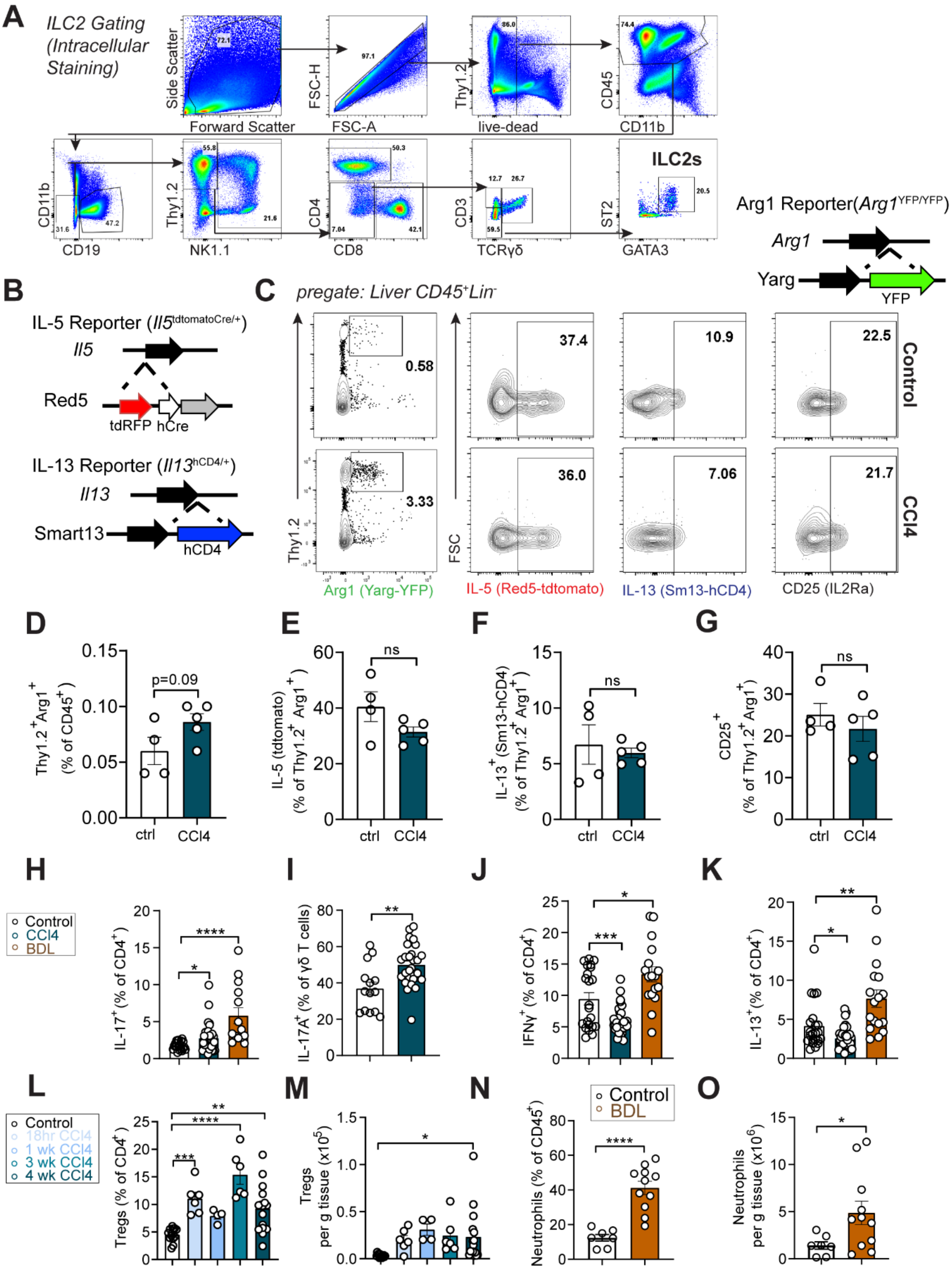


44

45

**Figure S2: Type 2 and type 3/17 lymphocytes expand in liver damage and fibrosis (related to Figure 2).**

(A) Representative flow gating scheme for liver ILC2s (viable $CD45^+Lin^-GATA3^+ST2^+$) and T cells.

(B) Schematic of the IL-$5^+$ lymphocyte lineage tracker mice (IL-5tdtomato-Cre; Rosa26RFP), *Il13*$^{Smart}$ (Smart13; B6.129S4[C]-*Il13*$^{tm2.1Lky}$/J; 031367), and *Arg1*$^{RFP\text{-}CreERT2}$ mice.

(C-G) Representative flow cytometry plots (C) and quantification of ILC2 ($Lin^-$ $Thy1.2^+Arg^+$) (D) expression of IL-5RFP (E), IL-13 (F), and CD25 (G) in livers of control mice and 4-week $CCl_4$-treated mice on Arg1 (Yarg); R5 (IL-5); S13 (IL-13) combined triple-reporter (YRS) background.

(H-K) Flow cytometry quantification showing percentages of IL-$17^+CD4^+$ T cells (H), IL-$17^+$ γδ T cells (I), $IFN\gamma^+CD4^+$ T cells (J), and IL-$13^+CD4^+$ T cells (K) in livers from C57BL/6 wildtype mice treated with vehicle or $CCl_4$ for 4 weeks. Pooled data from 4 independent experiments from 8-10 weeks-old age- and sex-matched mice.

(L and M) Flow cytometry quantification showing percentage (L) and total numbers (M) of Tregs in livers from IL-$33^{mcherry/+}$ reporter mice treated with vehicle or $CCl_4$ for the indicated duration. Pooled data from 3 independent experiments from 8-10 weeks-old age- and sex-matched mice for 4-week timepoint.

(N and O) Flow cytometry quantification of percentage (N) and total numbers (O) of neutrophils in livers from IL-$33^{mcherry/+}$ reporter mice 14 days post-BDL surgery. Pooled data from 3 independent experiments from 8-10-weeks-old age- and sex-matched mice.

Bar graphs indicate mean (±SE), unpaired t-test, *$p \leq 0.05$, **$p \leq 0.01$, ***$p \leq 0.001$, ****$p \leq 0.0001$.

### Supplementary Figure 3

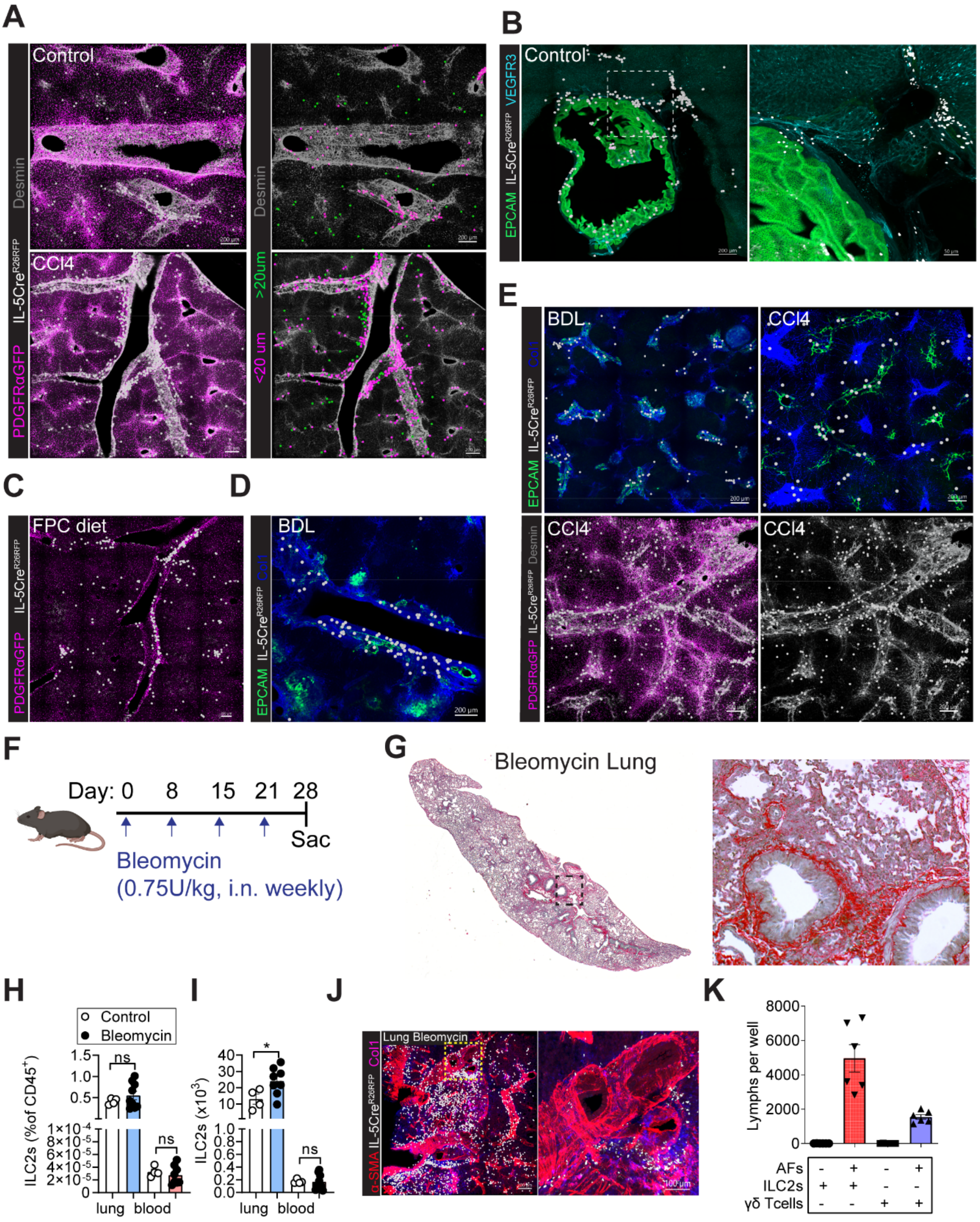

**Figure S3: Type 2 lymphocytes localize to periportal regions at steady state and expand into fibrotic tracts with liver fibrosis (related to Figure 3).**

(A) Representative confocal imaging of liver from IL-5$^+$ lineage tracker mice (IL-5tdtomato-Cre; Rosa26RFP) treated with vehicle or $CCl_4$ for 4 weeks with surface rendering of IL-5$^+$ lymphocytes and indicated staining. Images are representative of three or more mice.

(B) Representative confocal imaging of gallbladder and liver from naïve IL-5$^+$ lymphocyte lineage tracker mice with staining for EpCAM (epithelial duct cells) and VEGFR3 (lymphatics). Higher magnification highlighting accumulation of IL-5$^+$ lymphocytes around the bile ducts. Image is representative of n=3 mice.

(C-E) Representative confocal thick section imaging of liver from IL-5$^+$ lineage tracker mice fed with FPC diet for 16 weeks (C), 14 days post BDL surgery (D, E), treated with $CCl_4$ for 4 weeks (E), and surface rendering of IL-5$^+$ lymphocytes and indicated staining. Images are representative of three or more mice.

(F) Schematic showing Bleomycin administration schedule in IL-5 reporter mice subjected to weekly intranasal applications of Bleomycin (0.75U/kg), relevant to G-J.

(G) Representative Sirius red staining from IL-5tdtomato-Cre mice treated with Bleomycin for 4 weeks. Images are representative of one experiment with n=11 8-10 weeks-old age- and sex-matched mice.

(H and I) Flow cytometry quantitation of percent (H) and total numbers (I) of ILC2s in lung and blood from PDGFRαGFP; IL-33$^{mcherry/+}$ mice treated weekly with Bleomycin or PBS (intranasal application) for 4 weeks.

(J) Representative confocal imaging of cleared thick lung sections with surface analysis for IL-5RFP$^+$ lymphocytes from Il5-tdtomato-Cre mice treated weekly with Bleomycin (intranasal application) for 4 weeks. Images are representative of one experiment with n=11 Il5-tdtomato-Cre mice.

(K) PDGFRα$^+$Sca1$^+$ lung adventitial fibroblasts (AFs) were cultured with lung ILC2s and γδ T cells for 7 days; ILC2s and γδ T cells were counted. Pooled data from 2 independent experiments.

Bar graphs indicate mean (±SE). Unpaired t-test for D, ns= not significant, *$p \leq 0.05$.

### Supplementary Figure 4

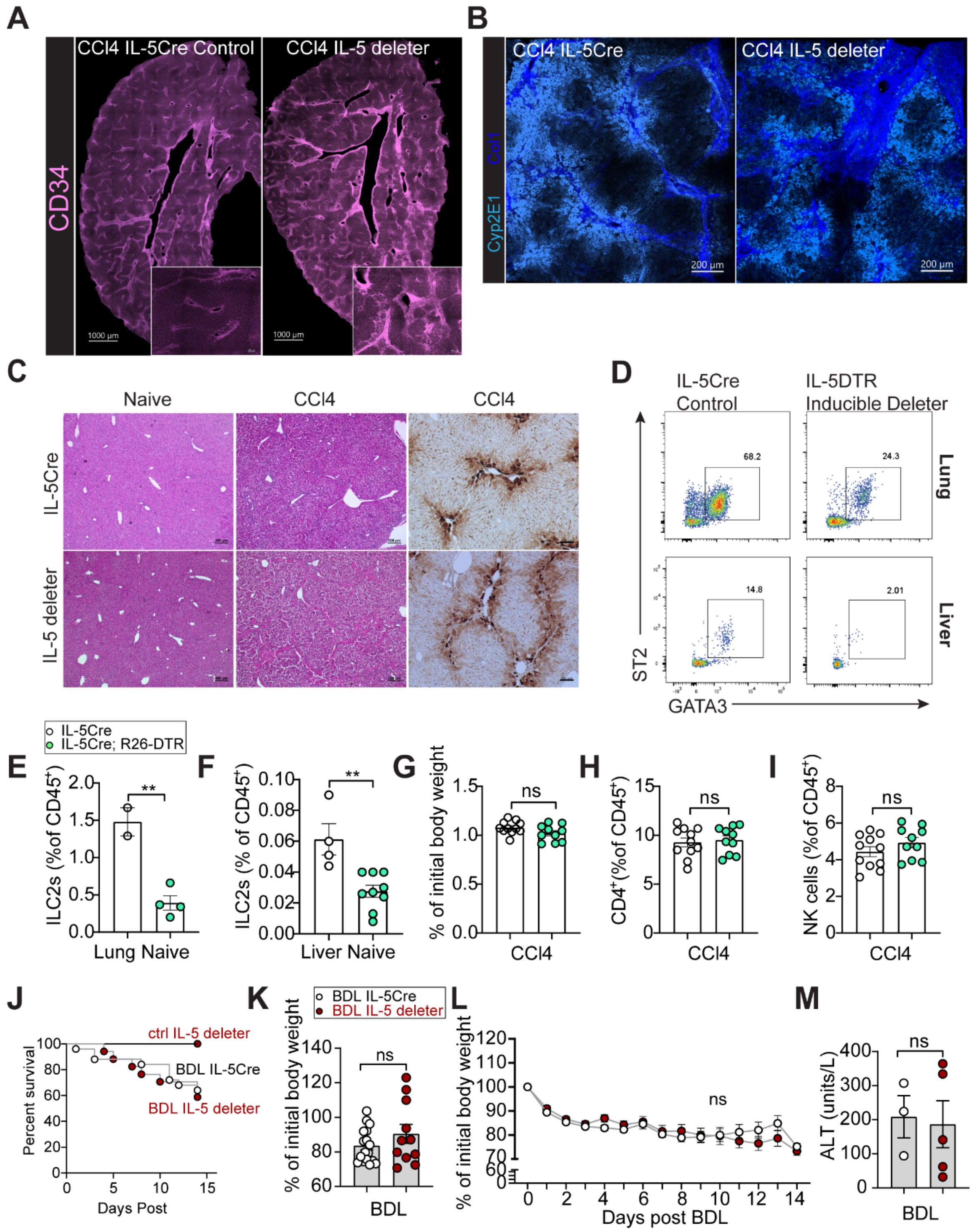

**Figure S4: Loss of IL-5$^{+}$ lymphocytes worsened hepatic fibrosis (related to Figure 5).**

(A) Representative confocal liver tissue sections from control or 4-week $CCl_4$-treated Il5-tdtomato-Cre mice and IL-5 deleter (RRDD; *Il5*$^{Red5/Red5}$*Gt (Rosa)26*$^{DTA/DTA}$) mice with staining for CD34. Images shown are representative of 3-4 mice/group.

(B) Representative confocal thick section imaging of liver from IL-5tdtomato-Cre and IL-5 deleter mice with staining for Collagen I and Cyp2E1 (marker of pericentral hepatocytes). Images are representative of three or more mice.

(C) Representative Sirius red staining and glutamin synthetase (GS) immunohistochemistry from IL-5tdtomato-Cre and IL-5 deleter (RRDD; *Il5*$^{Red5/Red5}$*Gt (Rosa)26*$^{DTA/DTA}$) mice treated with vehicle or $CCl_4$ for 4 weeks. Images are representative of 3 independent experiments with 8-10 weeks-old age- and sex-matched mice.

(D-F) Representative flow cytometry plots of ILC2s ($Lin^{-}Thy1.2^{+}GATA3^{+}ST2^{+}$) (D) and flow quantitation of ILC2 percentages in lung (E) or liver (F) from either IL-5tdtomato-Cre or IL-5$^{DTR}$ treated with $CCl_4$ for 4 weeks.

(G-I) Percent of initial body weight before $CCl_4$ treatment (G) and percent $CD4^{+}$ T cells (H) and percent NK cells (I) in livers of 4-week $CCl_4$-treated IL-5tdtomato-Cre and IL-5$^{DTR}$ mice. Pooled data from 2 independent experiments from 8-10 weeks-old age- and sex-matched mice.

(J-M) Kaplan-Meier-survival curves (J), body weight loss in percentage of initial body weight (K and L) after bile duct ligation (BDL), and ALT levels (M) from bile duct ligated Il5-tdtomato-Cre mice and IL-5 deleter mice. Data for survival curves are pooled from 3 independent experiments with n=25 BDL Il5-tdtomato-Cre mice, n=17 BDL IL-5 deleter mice, n=10 control Il5-tdtomato-Cre mice, and n=4 IL-5 deleter mice. Other data are pooled from 3 independent experiments with n=5-13 8-10 weeks-old age- and sex-matched mice/group.

Bar graphs indicate mean (±SE), Kaplan-Meier survival curves are compared using the log-rank (Mantel-Cox) analysis for J. unpaired t-test, $^{**}p \le 0.01$.

#### Supplementary Figure 5

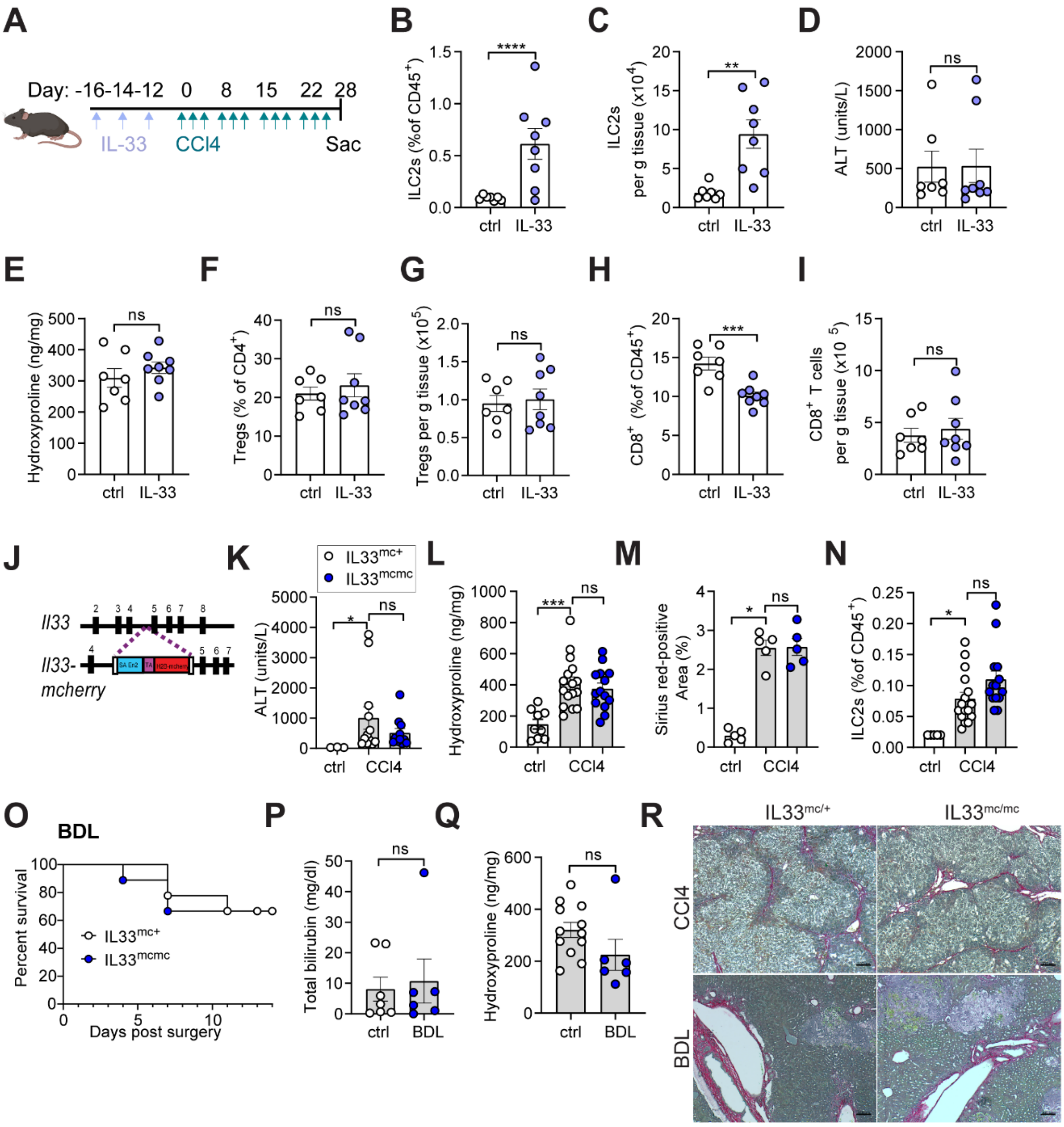

**Figure S5: Neither ILC2 expansion nor IL-33 signalling impacted degree of hepatic fibrosis.**

(A) Schematic of $CCl_4$ and IL-33 treatment (500ng i.p.) for IL-$^{33mcherry/+}$ mice.

(B and C) Flow cytometry quantitation of percent (B) and total numbers (C) of ILC2s in control and IL-33-treated IL-$33^{mcherry/+}$ mice after 4 weeks of $CCl_4$ treatment.

(D and E) Quantification of ALT (D) and hepatic Hydroxyproline levels (E) in control and IL-33-treated IL-$33^{mcherry/+}$ mice after 4 weeks of $CCl_4$ treatment.

(F-I) Flow cytometry quantitation of percent (F) or total numbers (G) of Tregs and percent (H) or total numbers (I) of $CD8^+$ T cells from control and IL-33-treated IL-$33^{mcherry/+}$ mice after 4 weeks of $CCl_4$ treatment.

(J) Diagram of the construction for the IL-33-H2B-mcherry nuclear localization IL-33 reporter.

(K-M) Quantification of ALT (K), hepatic Hydroxyproline (L), and Sirius red staining (M) from IL-$33^{mcherry/+}$ and IL-$33^{mcherry/mcherry}$ mice treated with $CCl_4$ for 4 weeks or controls. Pooled data from 3 independent experiments from 6-12 weeks-old age- and sex-matched mice.

(N) Flow cytometry quantification, showing percent of ILC2s in livers from IL-$33^{mcherry/+}$ and IL-$33^{mcherry/mcherry}$ mice treated with $CCl_4$ for 4 weeks or controls. Pooled data from 3 independent experiments from 6-12 weeks-old age- and sex-matched mice.

(O) Kaplan–Meier survival curves after bile duct ligation (BDL). Data are pooled from 2 independent experiments with n=9 IL-$33^{mcherry/+}$ mice and n=9 IL-$33^{mcherry/mcherry}$ mice.

(P and Q) Quantification of total bilirubin (P) and Hydroxyproline (Q) from IL-$33^{mcherry/+}$ and IL-$33^{mcherry/mcherry}$ mice 14 days post BDL surgery. Data are pooled from 2 independent experiments.

(R) Representative Sirius red staining from IL-$33^{mcherry/+}$ and IL-$33^{mcherry/mcherry}$ mice treated with $CCl_4$ for 4 weeks or 14 days post BDL surgery, as indicated.

Bar graphs indicate mean (±SE), unpaired t-test, *$p \le 0.05$, **$p \le 0.01$, ***$p \le 0.001$, ****$p \le 0.0001$.

### Supplementary Figure 6

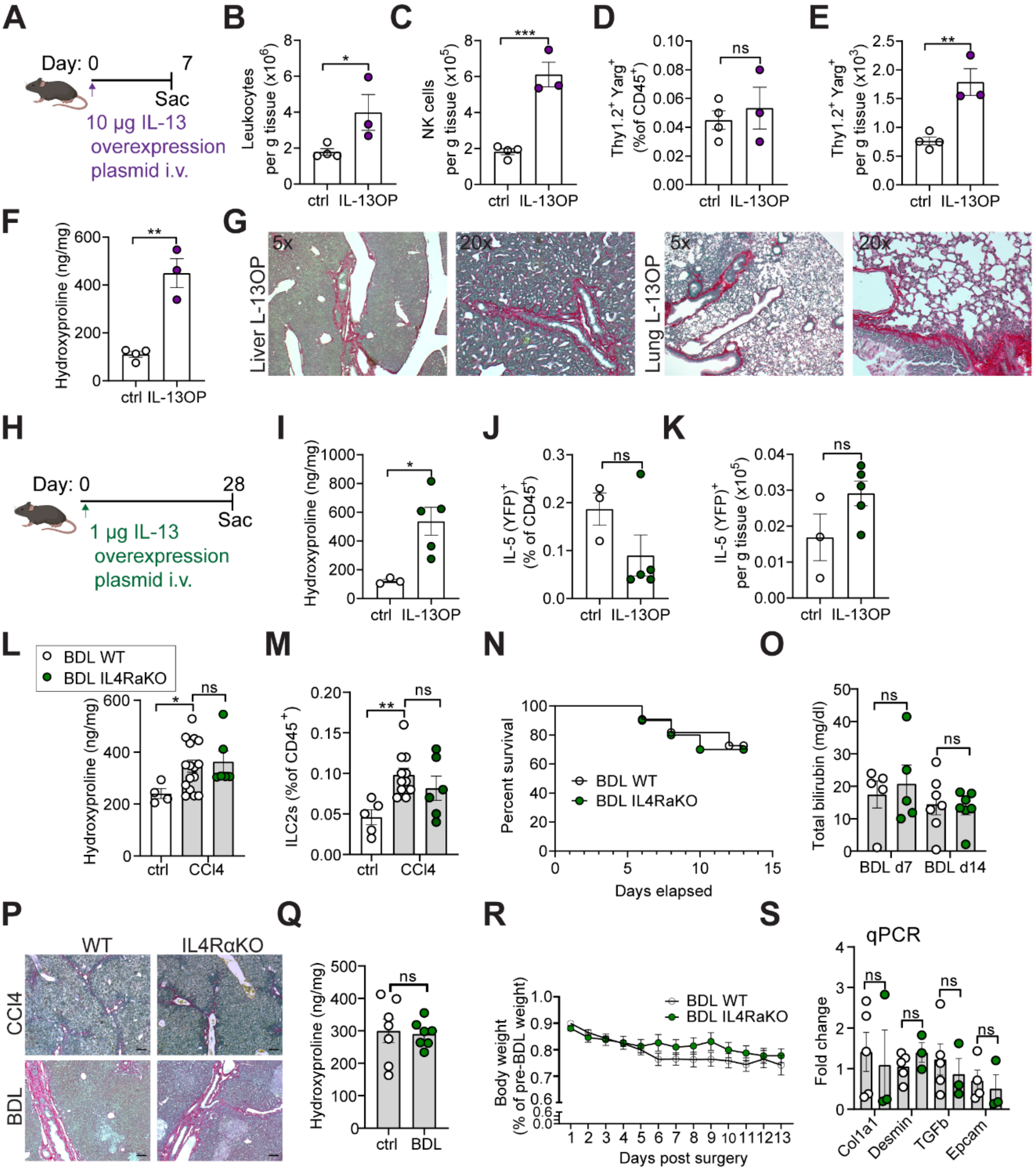

**Figure S6: The IL-4/IL-13 receptor does not regulate hepatic fibrosis in different models of liver injury.**

(A) Schematic showing intravenous (i.v.) treatment with 10µg IL-13 overexpressing plasmid in Arg1 (Yarg); R5 (IL-5); S13 (IL-13) combined triple-reporter (YRS) mice. Mice were harvested after 7 days.

(B-E) Flow cytometry quantitation of total leukocyte numbers (B), total numbers of NK cells (C), percent (D), and total numbers of ILC2s ($Lin^{-}Thy1.2^{+}Arg1^{+}$) (E) in control and $_{plasmid}$IL-13-treated YRS mice.

(F and G) Quantification of Hydroxyproline in control and $_{plasmid}$IL-13-treated YRS mice (F) and representative Sirius red staining from liver and lung of $_{plasmid}$IL-13-treated YRS mice (G). Images are representative of three or more mice.

(H) Schematic showing intravenous (i.v.) treatment with 1µg IL-13 overexpressing plasmid in IL-$5^{+}$ lymphocyte lineage tracker (IL-5tdtomato-Cre; RosaCAGYFP); IL-$33^{mcherry/+}$ double reporter mice. Mice were harvested after 4 weeks.

(I) Quantification of Hydroxyproline levels in control and $_{plasmid}$IL-13-treated IL-$5^{+}$ lymphocyte lineage tracker; IL-$33^{mcherry/+}$ double reporter mice.

(J and K) Flow cytometry quantification of percent (J) and total numbers (K) of IL-$5^{+}$ lymphocytes in control and $_{plasmid}$IL-13-treated IL-$5^{+}$ lymphocyte lineage tracker; IL-$13^{mcherry/+}$ double reporter mice.

(L and M) Quantification of Hydroxyproline (L) and flow cytometry quantification, showing percent of ILC2s (M) in liver from wild-type (WT) or IL-4/IL-13-deficient mice treated with vehicle or $CCl_4$ for 4 weeks.

(N) Kaplan–Meier survival curves after bile duct ligation (BDL) from wild-type and IL-4/IL-13-deficient mice. Data are pooled from 2 independent experiments with n=11 WT mice and n=10 IL-4/IL-13-deficient mice.

(O) Quantification of bilirubin from WT and IL-4/IL-13-deficient mice 14 days post BDL surgery. Pooled data from 2 independent experiments from 8-10 weeks-old age- and sex-matched mice.

(P) Representative Sirius red staining from WT and IL-4/IL-13-deficient mice treated with $CCl_4$ for 4 weeks or 14 days post BDL surgery, as indicated.

(Q) Quantification of Hydroxyproline from WT and IL-4/IL-13-deficient mice 14 days post BDL surgery. Pooled data from 2 independent experiments from 8-10 weeks-old age- and sex-matched mice.

(R) Body weight loss of WT and IL-4/IL-13-deficient mice post BDL surgery.

(S) Total liver gene expression of *COL1A1, DES, TGFB1*, and *EPCAM* normalized to *GAPDH* expression in control or bile duct ligated WT or IL_4/IL-13-deficient mice.

Pooled data from 2 independent experiments from 8-10 weeks-old age- and sex-matched mice.

Bar graphs indicate mean (±SE). Kaplan-Meier survival curves are compared using the log-rank (Mantel-Cox) analysis for N. Unpaired t-test for B-F, I-M, O, Q, and S, and two-Way ANOVA with Sidak post-test for R, $^{*}p \le 0.05$, $^{**}p \le 0.01$, $^{***}p \le 0.001$.

#### Supplementary Figure 7

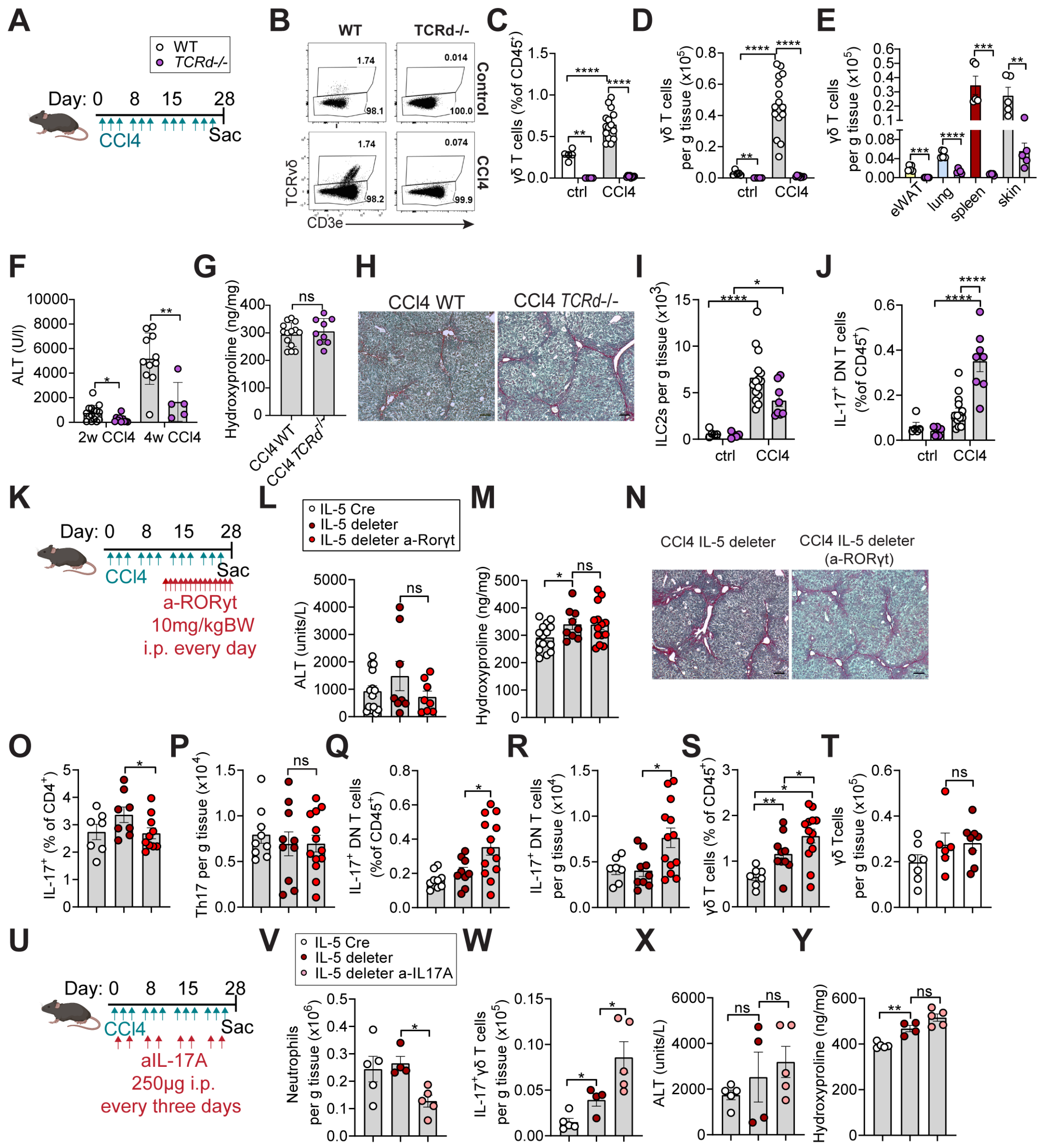


**Figure S7: Redundancy in type 3/17 lymphocytes during hepatic fibrosis (related to Figure 7).**

(A) Schematic showing $CCl_4$ administration schedule in WT and *TCRd*$^{-/-}$ mice injected intraperitoneally (i.p.) with 0.5µl $CCl_4$/g BW three times per week for 4 weeks (4w), relevant to B-J.

(B-D) Representative flow cytometry plots of γδ T cells (B) and quantification of percent (C) or total numbers (D) of γδ T cells in wild-type (WT) and *TCRd*$^{-/-}$ mice treated with vehicle or $CCl_4$ for 4 weeks.

(E) Quantification of total γδ T cell numbers in eWAT, lung, spleen, and skin in naïve adult WT and *TCRd*$^{-/-}$ mice.

(F and G) Quantification of ALT levels (F) and hepatic Hydroxyproline (G) in wild-type (WT) and *TCRd*$^{-/-}$ mice treated with $CCl_4$ for indicated duration. Data pooled from 2 independent experiments with 8-10 weeks-old age- and sex-matched mice.

(H) Representative Sirius red staining from WT and TCRd$^{-/-}$ mice treated with $CCl_4$ for 4 weeks. Images are representative of 2 independent experiments with 8-10 weeks old age- and sex-matched mice.

(I and J) Flow cytometry quantification, showing percent (I) and total numbers of ILC2s (J) in livers from WT and TCRd$^{-/-}$ mice treated with vehicle or $CCl_4$ for 4 weeks. Pooled data from 2 independent experiments from 8-10 weeks-old age- and sex-matched mice.

(K) Schematic showing treatment strategy for $CCl_4$ injected intraperitoneally (i.p.) with 0.5μl $CCl_4$/g BW three times per week for 4 weeks and RORγt antagonist injected with 10mg/kg BW i.p. every day for the last 2 weeks of the experiment with Il5-tdtomato-Cre mice and IL-5 deleter (RRDD; *Il5*$^{\text{Red5/Red5}}$*Gt (Rosa)26*$^{\text{DTA/DTA}}$), relevant to data in L-S.

(L and M) Quantification of ALT levels (L) and Hydroxyproline (M) in 4-week $CCl_4$-treated Il5-tdtomato-Cre mice, IL-5 deleter mice and IL-5 deleter mice treated with RORγt antagonist. Pooled data from 3 independent experiments with 8-10 weeks-old-age- and sex-matched mice.

(N) Representative Sirius red staining from 4-week $CCl_4$-treated IL-5 deleter and IL-5 deleter mice treated with RORγt antagonist. Images are representative of 3 independent experiments with 8-10 weeks-old age- and sex-matched mice.

(O-T) Flow cytometry quantification, showing percent (O) and total numbers (P) of Th17 cells, percent (Q) and total numbers (R) of IL-7$^{+}$ DN T cells, and percent (S) and total numbers (T) of γδ T cells in livers from Il5-tdtomato-Cre mice, IL-5 deleter, and IL-5 deleter mice treated with RORγt antagonist. Pooled data from 3 independent experiments with 8-10 weeks-old age- and sex-matched mice.

(U) Schematic showing $CCl_4$ and neutralizing anti-IL17A administration in Il5-tdtomato-Cre mice and IL-5 deleter mice, injected intraperitoneally (i.p.) with 0.5μl/g BW three times per week and with anti-IL17A antibody every three days throughout the experiment, relevant to U-X.

(V-Y) Flow cytometry quantification of total numbers of neutrophils (V) and IL-17$^{+}$ γδ T cells (W), and quantification of ALT levels (X) and Hydroxyproline (Y) in livers from Il5-tdtomato-Cre mice, IL-5 deleter mice, and IL-5 deleter mice treated with RORγt antagonists. Pooled data are representative of 3 independent experiments with 8-10 weeks-old age- and sex-matched mice.

Bar graphs indicate mean (±SE), unpaired t-test for C-G, I, J, L, M, and one-Way ANOVA for O-S and U-X, ns= not significant, *$p \leq 0.05$, **$p \leq 0.01$, ***$p \leq 0.001$, ****$p \leq 0.0001$.
